# Evolution and inhibition of the FIKK effector kinase family in *P. falciparum*

**DOI:** 10.1101/2024.02.22.581535

**Authors:** Hugo Belda, David Bradley, Evangelos Christodoulou, Stephanie D. Nofal, Malgorzata Broncel, David Jones, Heledd Davies, M. Teresa Bertran, Andrew G. Purkiss, Roksana W. Ogrodowicz, Dhira Joshi, Nicola O’Reilly, Louise Walport, Antoine Claessens, Andrew Powell, David House, Svend Kjaer, Christian R. Landry, Moritz Treeck

## Abstract

Among the ∼200 *Plasmodium* species that infect vertebrates, six infect humans. Of these, *P. falciparum* causes >95% of all ∼500,000 annual fatalities. Phylogenetically, *P. falciparum* belongs to the *Laverania* subgenus, a group of *Plasmodium* species that infect great apes. Common to *Laverania* species is the family of FIKK kinases. One million years ago, a single FIKK kinase conserved in all *Plasmodium* species gained an export element in the *Laverania* subgenus and expanded into the family of ∼20 atypical FIKK kinases, most of which are exported into the host cell. The *fikk* genes are conserved in syntenic loci across the *Laverania*, arguing for a rapid expansion controlling important functions in host cell remodelling and pathogenesis. We provide evidence that the FIKK paralogues evolved specific and mutually exclusive phosphorylation motif preferences, conserved across their *Laverania* orthologues, in a short evolutionary timeframe. Surprisingly, we find that FIKK13 has evolved exclusive tyrosine-phosphorylation preference, which was thought to be absent in *Plasmodium* species. Combining a crystal structure with AlphaFold2 predictions, we identify residues that determine kinase-specificity within the FIKK family in a fast-evolving flexible loop. Finally, we show that all expressed members of the FIKK kinase family can be chemically inhibited *in vitro* using a single compound. Such a pan-specific inhibitor of this kinase family important for virulence could reduce the ability of the parasite to gain escape-mutations and resistance.

## Introduction

Malaria is caused by the infection of red blood cells (RBCs) with *Plasmodium* parasites. ∼200 million infections and 500,000 deaths are observed annually, with severe cases occurring primarily in children under the age of 5^1^. Among the 6 *Plasmodium* species infecting humans, *P. falciparum* causes over 95% of all fatalities. This species remodels RBCs to strongly cytoadhere to the host endothelium causing sequestration of infected RBCs (iRBCs), preventing passage through the spleen in which iRBCs can be recognised and destroyed. While benefiting the parasite, cytoadhesion can lead to severe disease through the formation of blood clots in capillaries, reducing oxygen supply to highly vascularised organs such as the brain, lungs, kidneys, or placenta in pregnant women.

*P. falciparum* exports ∼10% of its proteome into the host cell^2^. Exported proteins fulfil a variety of functions in the iRBC^3^. They facilitate transport and anchoring of the major cytoadhesion ligand *P. falciparum* Erythrocyte Membrane Protein 1 (*Pf*EMP1)^4^ into parasite-derived structures underneath the erythrocyte membrane (knobs)^5^, the creation of new nutrient permeability pathways in the plasma membrane^6,7^ and the formation of intracytoplasmic membranous structures called Maurer’s clefts that help traffic parasite proteins to the host cell surface^8^.

Among the parasite exported proteins is a family of serine/threonine kinases called the FIKK kinases. FIKKs are exclusive to apicomplexan parasites^9^. While most *Apicomplexa* possess one non-exported FIKK kinase (FIKK8 in *P. falciparum*), gene expansion in *P. falciparum* resulted in a family of 21 paralogues, including 2 predicted pseudogenes in the 3D7 reference genome^10^. All FIKK kinases, except for FIKK8, are predicted to be exported into the RBC. The expanded FIKK kinase family is found in all *Plasmodium* species of the *Laverania* subgenus, which includes *P. falciparum* and *Plasmodium* species infecting great apes^11,12^, but no other human-infecting species. No *Plasmodium* species outside the *Laverania* contains predicted exported kinases. 10 of the 19 active *P. falciparum fikk* genes are conserved in syntenic loci in all *Laverania* species and all *fikk* genes are conserved in syntenic loci in at least 4 of the *Laverania* species (Supplementary Table 1, data from PlasmoDB (www.plasmodb.org)^13–15^). The minimum number of FIKK kinases present in any *Laverania* species is 16, but this number may be higher because of low quality genomes regions of some *Laverania* species. This indicates that *fikk* genes rapidly multiplied and diversified early during *Laverania* evolution. The expansion of the FIKK family was followed by a long period of stasis in terms of the *fikk* copy number, suggesting that FIKK kinases individually play important roles in host-pathogen interactions in *Laverania* hosts.

At least one FIKK kinase (FIKK4.1) is important for *Pf*EMP1 surface translocation and cytoadhesion^16^, while FIKK4.2 is important for iRBC rigidification^17^. We previously observed no reduction in growth upon individual conditional deletion of any exported FIKK kinase^16^. This suggested that either exported FIKK kinases play no role in growth under standard cell culture conditions, or that there is a level of redundancy and compensation between FIKK kinases. While it is tempting to speculate an important role for each FIKK based on their conservation across the *Laverania*, redundant functions cannot be ruled out. Determining the degree of redundancy between FIKK kinases is paramount to design experiments understanding their functions during *P. falciparum* infections.

FIKK kinases contain a variable N-terminus that is unique to paralogues but conserved within orthologous groups, and a conserved C-terminal kinase domain containing the eponymous Phe-Ile-Lys-Lys (F-I-K-K) motif. FIKKs lack the glycine triad involved in binding ATP^9,18^ but at least 14 FIKKs have demonstrated activity^16,17,19–26^, indicating that they coordinate ATP in a non-classical manner. The unique ATP-binding pocket along with a small gate-keeper residue^27^ found in most FIKKs may provide opportunities for developing highly specific pan-FIKK inhibitors that target several or all FIKKs simultaneously.

Here, we provide evidence that a core set of FIKKs is under strong positive selection and required for human infection. Our data suggest that FIKK kinases specificity underwent a rapid diversification during the expansion of the kinase family, which is partly due to a fast-evolving loop in the kinase’s substrate-binding region. This diversification appears conserved among distantly related *Plasmodium* species, suggesting evolutionary constraint linked to important functions in host-pathogen interaction with great apes and humans. Finally, we demonstrate that chemical inhibition of the FIKK kinases is achievable and that their highly conserved kinase domain allows for the development of pan-FIKK inhibitors.

## Results

### Potential overlapping and non-overlapping functions of the FIKK kinases

To identify potentially overlapping functions between FIKK kinases, we searched for FIKKs that are expressed at similar timepoints and co-localize. 19 active FIKKs appear to be transcribed in asexual parasite stages (Fig. 1a, Supplementary Table 2), although some only at very low levels. This is in line with our previous observation that some FIKKs are barely detectable as HA-tagged variants^16^. The main subcellular localisations are punctate staining in the RBC cytosol, likely representing Maurer’s clefts or related structures, and the RBC periphery/knobs (Fig. 1b). In addition to FIKK8^16,25^, four FIKKs (FIKK9.2^19^, FIKK3, FIKK9.5^28^ and FIKK5^16^) have reported localisations within the parasite, although antibodies raised against FIKK kinases have not yet been verified using available FIKK knockout lines.

**Fig. 1.**
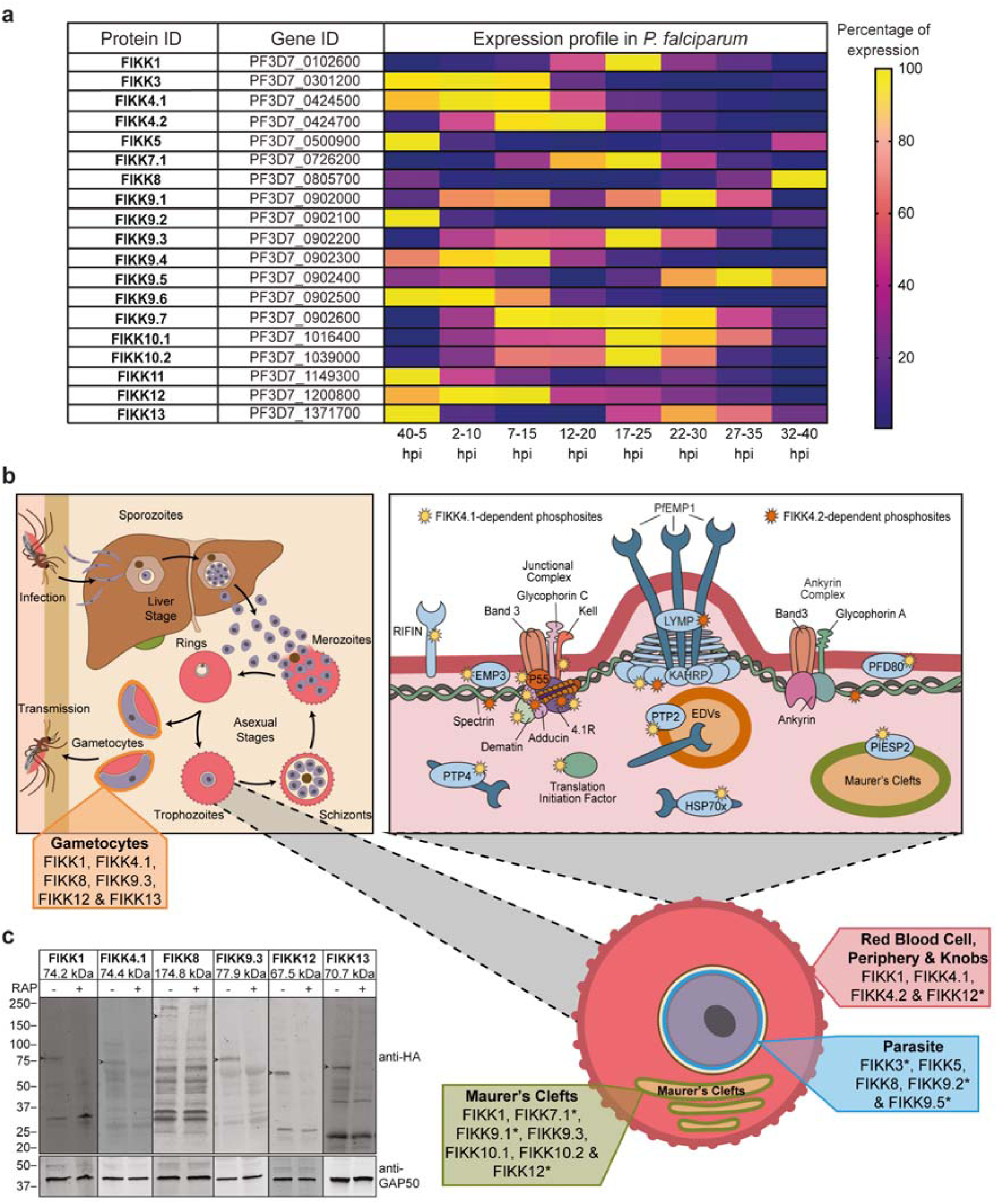
Expression timings and localisations of *P. falciparum* FIKK kinases. **a**, Heatmap built using data from Hoeijmakers *et al.*^30^ RNA-sequencing dataset available on PlasmoDB (www.PlasmoDB.org), showing % expression for each FIKK kinase during the *P. falciparum* asexual replication cycle. yellow = maximum expression; dark blue = minimum expression. TPM (Transcript Per kilobase Millions) numbers used to calculate % expression relative to maximum expression across the 48h lifecycle are available in Supplementary Table 2. **b**, Diagram illustrating *P. falciparum* FIKK kinase expression and localisations in iRBCs. Top left panel: illustration of the *P. falciparum* lifecycle. FIKK1, FIKK4.1, FIKK8, FIKK9.3, FIKK12 and FIKK13 expressed in gametocytes are shown in orange. Bottom right panel: *P. falciparum* iRBC showing the localisation of the FIKK kinases. blue: FIKK3, FIKK5, FIKK8, FIKK9.2 and FIKK9.5 in the parasite; green: FIKK1, FIKK7.1, FIKK9.1, FIKK9.3, FIKK10.1, FIKK10.2 and FIKK12 in Maurer’s clefts; red: FIKK1, FIKK4.1, FIKK4.2 and FIKK12 at the RBC periphery. “*” indicates localisation data from publications from other laboratories. Top right panel: knob structure at the RBC periphery. Yellow stars show FIKK4.1 substrates and orange stars show FIKK4.2 substrates (data from ^16^). **c**, Western blots confirming expression of HA-tagged *P. falciparum* FIKK kinases in gametocytes stage III. GAP50 antibody (bottom) demonstrates equal loading. Arrows show FIKK bands at expected sizes (shown in the labels at the top). A “+” sign indicates rapamycin treatment.

Two of the 21 FIKK kinases (FIKK7.2 and FIKK14) are annotated as pseudokinases in the 3D7 reference genome but not in other genetic backgrounds. This suggests that some FIKKs may still be evolving and dispensable for human infections. To identify other *Pf*FIKKs that may have lost functions in humans, we searched 2,085 available field isolate genomes for FIKK kinases with internal stop codons, or deletions (Supplementary Table 3). Three *fikk* genes show internal stop codons in >1% of all sequenced genomes (*fikk7.2*, *fikk9.2*, *fikk14*). 55.16% (1150/2085) and 2.73% (57/2085) of all field isolates contain a stop codon in *fikk7.2* or *fikk14* genes, respectively. For *fikk7.2*, 93.7% (1078/1150) of mutations are identical (W413*) and are equally distributed between South-East Asia (SEA) and Africa, suggesting an ancient origin. In contrast, *fikk14* shows different premature stop codons throughout the gene, predominantly in African isolates (94.7% (54/57)), indicating that inactivating mutations in *fikk14* are not systematically eliminated by natural selection. Interestingly, 11.44% (137/1197) and 12.05% (107/888) of African and SEA samples, respectively, have *fikk14* deletions, so the preponderance for stop codons in *fikk14* in African isolates is not observed for gene deletions (Supplementary Table 4). *fikk9.2* encodes an active kinase in the 3D7 reference strain, but 4.65% (97/2085) of the field isolate genomes contain an internal stop codon, mainly in SEA isolates (86.6% (84/97)). Collectively, these data suggest that a core set of 18 FIKK kinases are under purifying selection, while three FIKK kinases are under relaxed selection in humans, since inactivating mutations can arise in the field. Relaxed selection could come from redundancy among the FIKK kinases. Alternatively, since these kinases likely evolved in the great ape-infecting ancestors of *P. falciparum*, they may have fulfilled important functions which are now expendable during human infection. An argument for the latter hypothesis is the observation that both *fikk7.2* and *fikk14* are predicted to be functional in *P. praefalciparum*, *P. gaboni* and *P. adleri*^12^.

At least 12 FIKK kinases are likely to be expressed in gametocytes^29^, the sexual stages of the parasite that develop in the RBC and are taken up by a mosquito for onward transmission (Supplementary Table 5). We confirmed the expression of FIKK1, FIKK4.1, FIKK8, FIKK9.3, FIKK12 and FIKK13 in stage III gametocytes using HA-tagged conditional knockout lines^16^ (Fig. 1c). Transcriptomic data also suggest that some FIKK kinases may be expressed during liver infection and/or in parasite stages present in the mosquito, although this has not been experimentally tested (Supplementary Table 5).

Collectively, these data suggest that some FIKK kinases are separated in time and space within the iRBC and therefore likely evolved unique functions. Other FIKK kinases however have similar localisations within the cell and partially overlapping expression timings and could therefore have functional overlaps.

### FIKK4.1 and FIKK4.2 have partially overlapping subcellular localisations

To test whether co-localising FIKKs partially overlap in their function, we explored FIKK4.1 and FIKK4.2 in more depth. These two kinase genes are located in close proximity on chromosome 4, are phylogenetically closely related (Extended Data Fig. 1) and originated from a gene duplication event. Both co-localise at the iRBC periphery by immunofluorescence microscopy (IFA) (Fig. 2a) where they phosphorylate host cytoskeleton and exported parasite proteins (Fig. 1b, top right panel)^16^. FIKK4.1 deletion reduces PfEMP1 surface translocation by ∼50%^16^, but this is not observed upon FIKK4.2 deletion^31^. While this demonstrates that FIKK4.1 deletion cannot be fully compensated by FIKK4.2, a partial substrate overlap and partial rescue cannot be excluded.

**Fig. 2.**
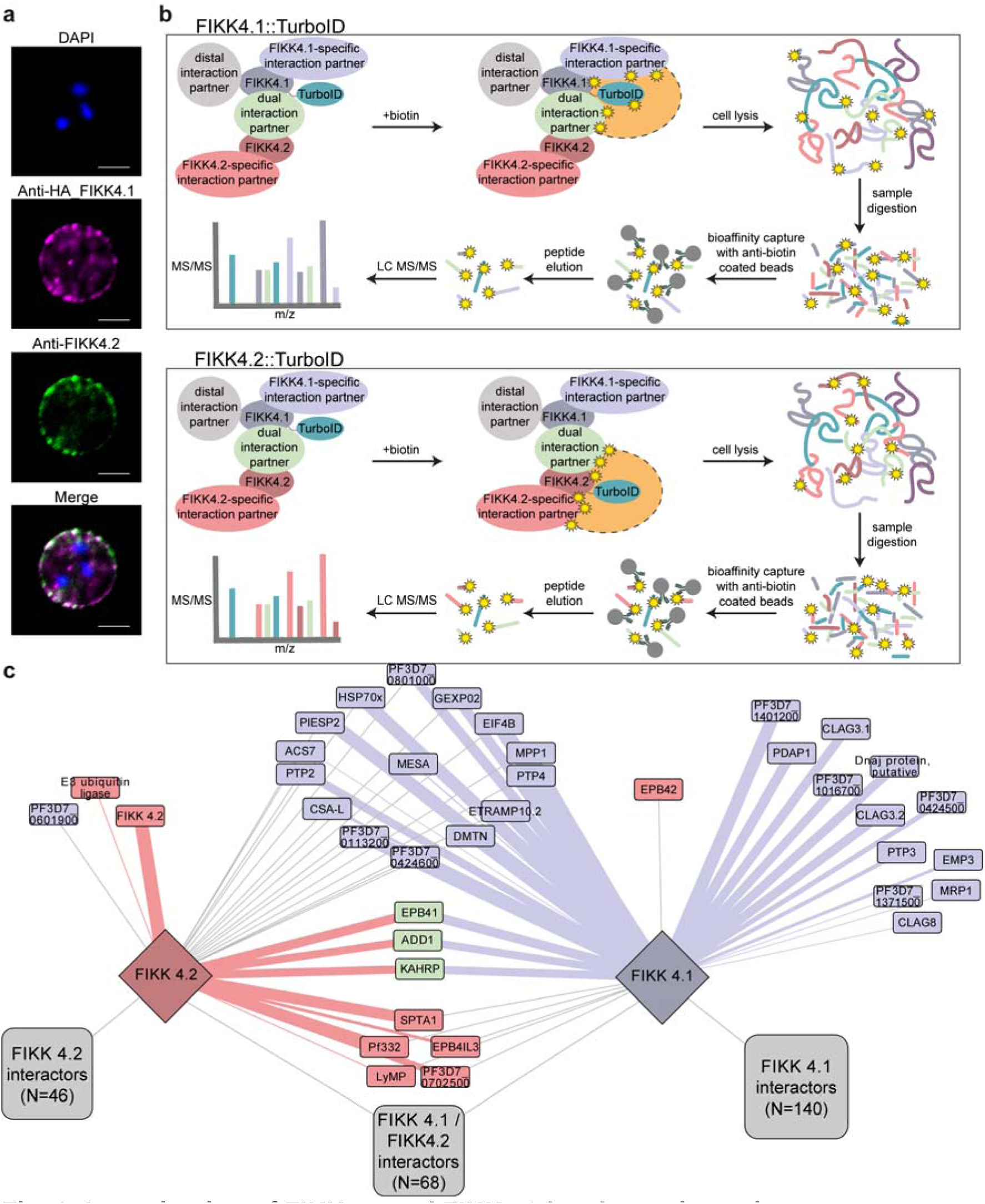
Investigation of FIKK4.1 and FIKK4.2 local protein environment. **a**, Subcellular localisation of FIKK4.1 and FIKK4.2 investigated by immunofluorescence analysis using anti-HA antibodies (magenta) targeting the C-terminally HA-tagged FIKK4.1 and anti-FIKK4.2 antibodies (green). DAPI (blue) is used for nuclear staining. Scale bar = 5µm. **b,** Diagram representing the proximity labelling workflow. FIKK4.1 (top panel) and FIKK4.2 (bottom panel) were tagged with a TurboID biotin ligase. Upon addition of biotin, proteins in the vicinity (represented by an orange area with dashed outline) of the bait are biotinylated on lysine residues (represented by yellow stars). iRBCs were lysed in 8M Urea in 50mM HEPES and proteins were Trypsin-digested into peptides. Biotinylated peptides were enriched using beads coated with two different anti-biotin antibodies and analysed by LC-MS/MS. **c,** Network analysis of FIKK4.1 and FIKK4.2::TurboID data. Connecting lines indicate a protein that is likely in the vicinity of the TurboID-tagged protein. Blue depicts proteins that have been identified as potential FIKK4.1 direct targets in ^16^. Red depicts proteins that have been identified as potential FIKK4.2 direct targets and green depicts proteins that have been identified as potential targets of both FIKK4.1 and FIKK4.2. Thickness of the connection represents how well the phosphorylation site matches the corresponding *in vitro* preferred phosphorylation motifs (of FIKK4.1 or FIKK4.2) from Extended Data Fig. 3 and 4.

To gain high-resolution information on their subcellular localisation, we determined their local protein environment by proximity labelling using TurboID fusion proteins^31,32^ (Extended Data Fig. 2) and mass spectrometry (Fig. 2b). Proteins that are found to be biotinylated by either or both FIKK::TurboID fusions were mapped onto a protein network, and then overlaid with our previous data showing if they could be phosphorylated by either of the two FIKKs^16^. 91 proteins were biotinylated by FIKK4.1::TurboID and FIKK4.2::TurboID and are therefore in close spatial proximity to both kinases (Fig. 2c). However, each of the two fusion proteins also labels a unique subset of proteins indicating that they are not in identical locations. In support of that, we found no evidence of reciprocal biotinylation of FIKK4.1– and FIKK4.2::TurboID fusions (Supplementary Table 6). Phosphorylation of only three proteins is dependent on both FIKK4.1 and FIKK4.2 (α-adducin, protein 4.1 and KAHRP) but the phosphorylated residues are not overlapping. Phosphorylation of all other proteins in proximity of both kinases is exclusively dependent on only one of the two kinases. These data suggest that FIKK4.1 and FIKK4.2 are located in very close, but not direct proximity and evolved to phosphorylate different targets.

### FIKK kinases evolved unique phosphorylation motifs

To gain insights into how FIKK kinases may have evolved to phosphorylate specific targets we determined their preferred phosphorylation motifs. Most eukaryotic protein kinases preferentially phosphorylate S, T, or Y residues within a specific amino acid sequence context (motif). These motifs are broadly classified into acidic, basic, or proline-directed^34^. *P. falciparum* kinases phosphorylate S and T residues within acidic and basic motifs. Phosphorylated Y residues and proline-directed motifs are rarely found^35^ and predicted tyrosine kinases are lacking from the genome. We attempted to recombinantly express the kinase domains of all predicted active *Pf*FIKKs (Fig. 3a) (Extended Data Fig. 3, 4 and Supplementary Table 7) and assessed substrate specificity on S, T and Y residues using OPAL libraries (Oriented Peptide Array Library)^36^. Of the 19 FIKKs, only FIKK9.6 and FIKK9.7 were refractory to bacterial expression.

**Fig. 3.**
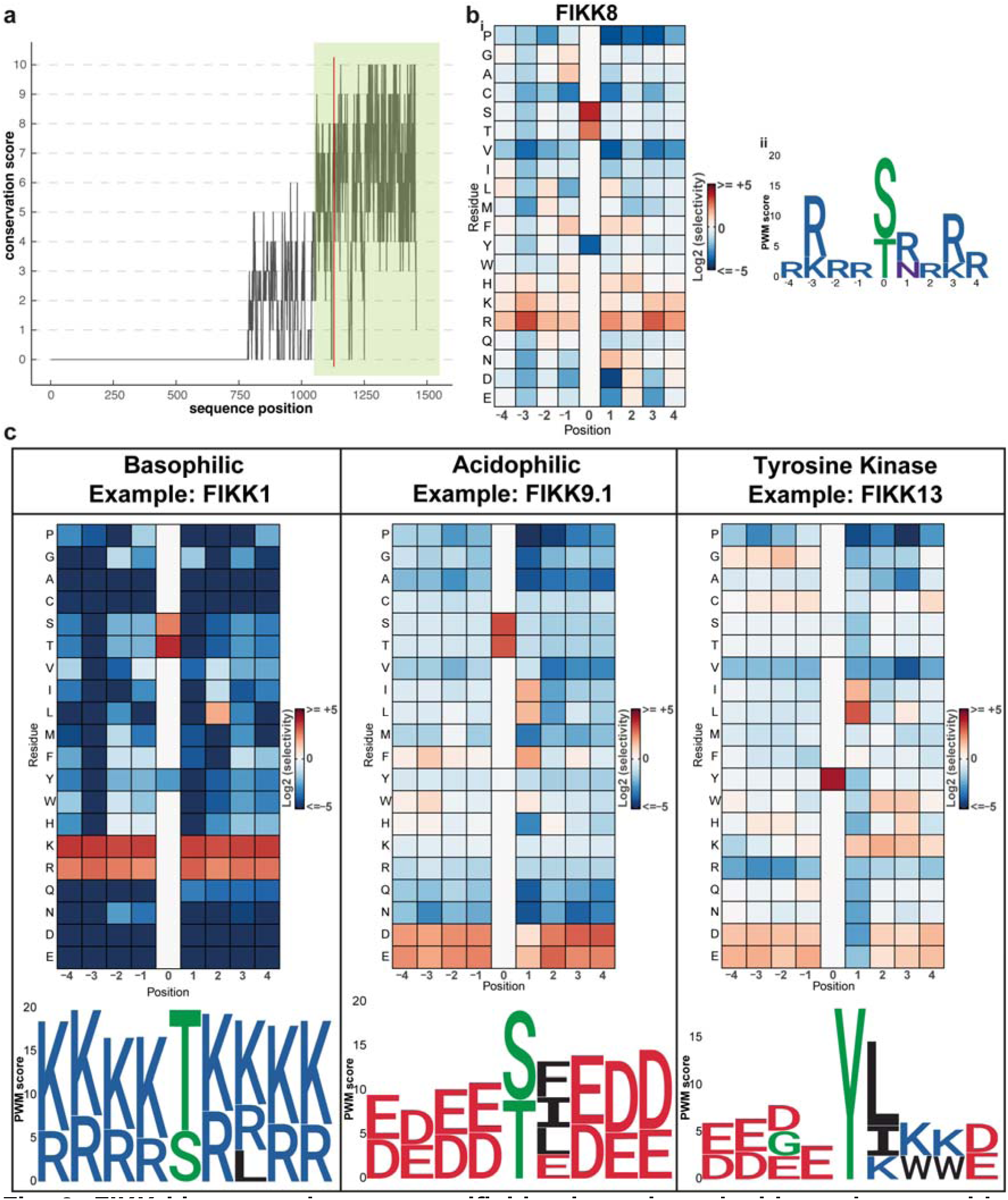
FIKK kinases substrate specificities investigated with random peptide libraries. **a**, Amino acid sequence conservation in *P. falciparum* FIKK kinases was assessed using the PRALINE Multiple Sequence Alignment Software^46^. Conservation values reflect the normalised average of BLOSUM62 scores for each alignment column and range from 0 (low conservation) to 10 (high conservation). Sequence position is with respect to *P. falciparum* FIKK8 as the reference sequence. Green shading illustrates the FIKK kinase domain. The eponymous F-I-K-K motif is represented in red. **b_i_**, Extended Data Fig. 5 data represented as a heatmap. ^32^P incorporation values were normalised to 20 (number of possible natural amino acids) and are shown as Log2(x) where negative values (blue cells) indicate disfavoured amino acids and positive values (red cells) indicate favoured amino acids. **b_ii_**, PWM logo generated with FIKK8 raw OPAL data. PWMs depict the preference of the kinase for all 20 amino acids at every substrate position. For ease of visualisation, the PWM logo displays amino acids with scores above an arbitrary threshold of 2.5 (see Material and Methods). Amino acid colours are set as: Acidic negatively charged (D, E) Red; Basic positively charged (R, K, H) Blue; Polar uncharged (N, Q) Purple; Nonpolar (A, I, L, M, F, V, P) Black; Phosphorylatable or Special (S, T, Y, C, G) Green. **c,** Heat map representation of OPAL data for basophilic FIKK1 (left panel), acidophilic FIKK9.1 (middle panel) and tyrosine kinase FIKK13 (right panel). PWM logos generated from raw OPAL data are displayed below the corresponding heatmaps. OPAL membrane images are available in Extended Data Fig. 8.

As previously reported^22^, FIKK8, which likely represents the closest relative to the ancestral kinase from which all FIKKs evolved in the *Laverania,* shows a preference for basic residues (Fig. 3b_i_, Extended Data Fig. 5). Position Weight Matrices (PWMs) indicate especially strong preference for arginine and/or lysine residues in position P-3 and P+3 (Fig. 3b_ii_). Eleven FIKKs prefer basic and positively charged amino acids surrounding the phosphorylated residue, while five FIKKs favour acidic motifs (Fig. 3c, Extended Data Fig. 6, 7). Within both groups, nuanced preferences emerge. FIKK1 strongly prefers a hydrophobic residue in the P+2 position, distinguishing it from other FIKK kinases. FIKK9.3 and FIKK9.4 show a mix of basic and acidic residues in the motif; here assignment to a group was based on the dominant charge.

Interestingly, we observed several FIKKs that phosphorylate peptides with a central Y. For some of these (FIKK5, FIKK8, FIKK9.1 or FIKK12) S/T residues in the flanking regions of the central Y may explain the signal, while others (FIKK3, FIKK4.2, FIKK9.2, FIKK9.3, FIKK9.4, FIKK9.5 and FIKK11) exhibited dual specificity. FIKK13 showed exclusively Y phosphorylation activity (Fig. 3c). This is a surprising result considering the absence of known *bona-fide* tyrosine kinases in *Plasmodium* or indeed any *Apicomplexa*^37^. This result suggests that FIKK13 has evolved from a S/T kinase into a tyrosine kinase, potentially to interact with specific host-cell proteins. To validate this result, we screened a DNA-encoded cyclic peptide library (RaPID selection, Extended Data Fig. 9a)^38–41^ and enriched four cyclic peptides that bind to FIKK13 (FIKK13_2 K_D_=310±290nM, FIKK13_3 K_D_=7±5nM, FIKK13_4 K_D_=17±0.4nM and FIKK13_5 K_D_=120±156nM) (Extended Data Fig. 9b and Supplementary Table 8). One peptide (FIKK13_4: cyclic-d(Y)PLRFLSKYHC(S-)G-CONH_2_) was identified as an *in vitro* substrate for FIKK13 and phosphorylation depended on the tyrosine residue (Extended Data Fig. 9c, d). This further supports FIKK13 function as a tyrosine kinase.

In summary, FIKK kinases evolved divergent substrate specificities from a basophilic ancestor, thereby expanding the repertoire of proteins the parasite can regulate. The motif diversity of the FIKK kinases highlights the rapid evolution of this relatively young protein family^14^, likely due to selection pressure to subvert the host machinery. This contrasts with ancient kinase families such as CK1, MAPK or PKA kinases that possess more highly conserved phosphorylation motifs^42–45^.

### FIKK phosphorylation motifs are conserved in distantly related *Laverania* species

To further confirm the specificity of FIKK13 for tyrosine residues, we expressed its orthologue from *P. gaboni*, the most distantly related *Laverania* species to *P. falciparum*, which is estimated to have diverged ∼1 million years ago. If *Pg*FIKK13 is evolving under purifying selection against changes to specificity, it should possess a similar tyrosine-based preferred phosphorylation motif. We also expressed *Pg*FIKK1 and *Pg*FIKK9.1 as examples of a basophilic and acidophilic kinases, to test whether the preferences for charges are retained. Strikingly, the motifs of the *P. falciparum* FIKKs are nearly identical to their *P. gaboni* orthologues (Fig. 4a, Extended Data Fig. 10). FIKK kinase orthologues are more similar between species than paralogues within the same species (Fig. 4b). Considering the strong motif preference observed for *Pf* and *Pg* FIKK1, 9.1 and 13, this suggests that FIKK kinase substrate specificity is probably conserved across all *Laverania* species. The analysis also shows almost equal divergence between FIKK paralogues in terms of their sequence identity (Extended Data Fig. 11). Therefore, in most cases, the precise evolutionary relationship between paralogues cannot be resolved with confidence, in agreement with the rapid and early diversification of the FIKK kinase family. Exceptions are the pair of recent paralogues FIKK9.5 and FIKK9.6, and the FIKK4.1-FIKK4.2-FIKK10.2 clade that are both predicted with high confidence (Fig. 4b, Extended Data Fig. 11). Together with the broad diversity of preferred phosphorylation motifs and their deep conservation between orthologues, it appears that the ancestor to FIKK8 rapidly diversified into the 16+ exported copies that we see in the *Laverania* subgenus.

**Fig. 4.**
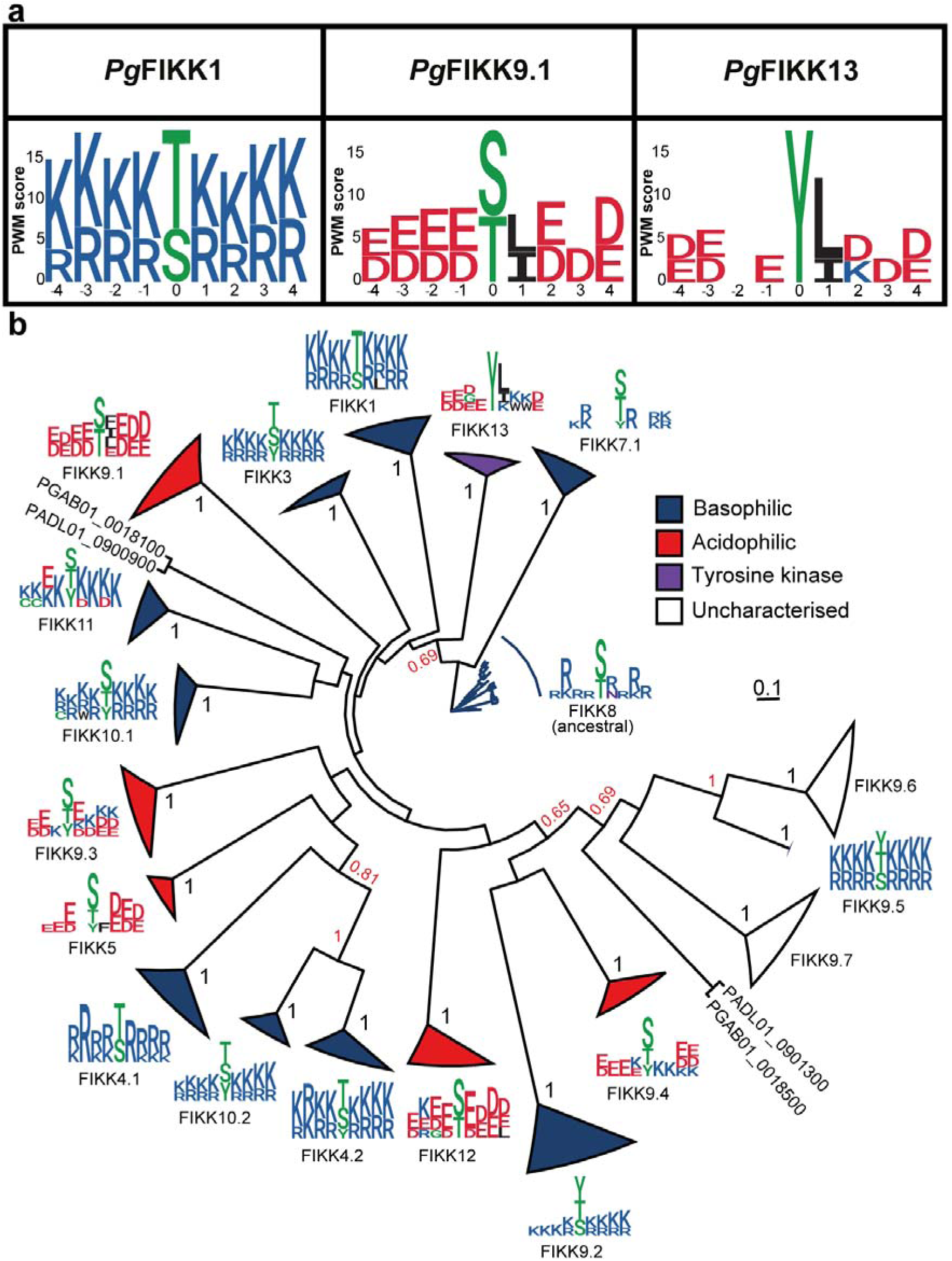
FIKK kinases substrate specificities are conserved among *Laverania* species. **a**, PWM logo generated with *Pg*FIKKs raw OPAL data. See Fig. 3 caption. **b,** Maximum-likelihood phylogenetic tree of *Laverania* FIKK kinase sequences built using FIKK8 kinases and two avian malaria FIKK kinases (*P. relictum* FIKK kinase PRELSG_0112400 and *P. gallinaceum* FIKK kinase PGAL8A_00108200) as an outgroup. 100 bootstrap replicates were generated to assess branch support^50^. All orthologue clades have maximum branch support (1 out of 1). Branches between paralogues are highlighted in red if they are > 0.5. Triangle length represents the divergence between FIKK kinase sequences within a specific clade. Colour code identifies the kinases substrate specificities as follows, blue = basophilic, red = acidophilic, purple = tyrosine kinase, white = uncharacterised. Sequence logos for each clade are given for the *P. falciparum* kinase copy.

### Divergent substrate motifs between FIKK kinases in similar subcellular localisations allow distinct regulation of targets

FIKK1, FIKK4.1 and FIKK4.2, which all localise to the RBC periphery (Fig. 1), share a basic preferred phosphorylation motif, although the specificity maps differ slightly. FIKK1 prefers a hydrophobic leucine residue in the P+2 position, while FIKK4.1 has a strong preference for an arginine residue in the P-3 position (Fig. 3, Extended Data Fig. 6).

To test whether the *in vitro* phosphorylation motifs identified here match the targets we previously identified by conditional FIKK deletion and phosphoproteomics in cell culture^16^, we performed activity assays on membranes containing 215 peptides predicted to be targets of FIKK1, FIKK4.1 and FIKK4.2. We also included 89 peptides that are targeted by FIKK10.2, a basophilic FIKK kinase which localises to the Maurer’s clefts, and 93 peptides from host cell proteins found to be more phosphorylated upon *P. falciparum* infection (Fig. 5a) (see Supplementary Table 9 for peptides sequences). The membranes were incubated with either recombinant FIKK1, FIKK4.1, FIKK4.2 or FIKK10.2 kinase domains and [γ-32P]-ATP (Fig. 5a, Extended Data Fig. 12a). Phosphorylation motifs from the OPAL libraries of randomised peptides largely correspond to the motifs of natural peptides that are strongly phosphorylated in this assay (Fig. 5b, Extended Data Fig. 12b). This is shown in Fig. 5c for FIKK1 where its basophilicity is confirmed and highly phosphorylated peptides feature a leucine at the P+2 position – a FIKK1 signature – making it the most highly specific kinase from this dataset.

**Fig. 5.**
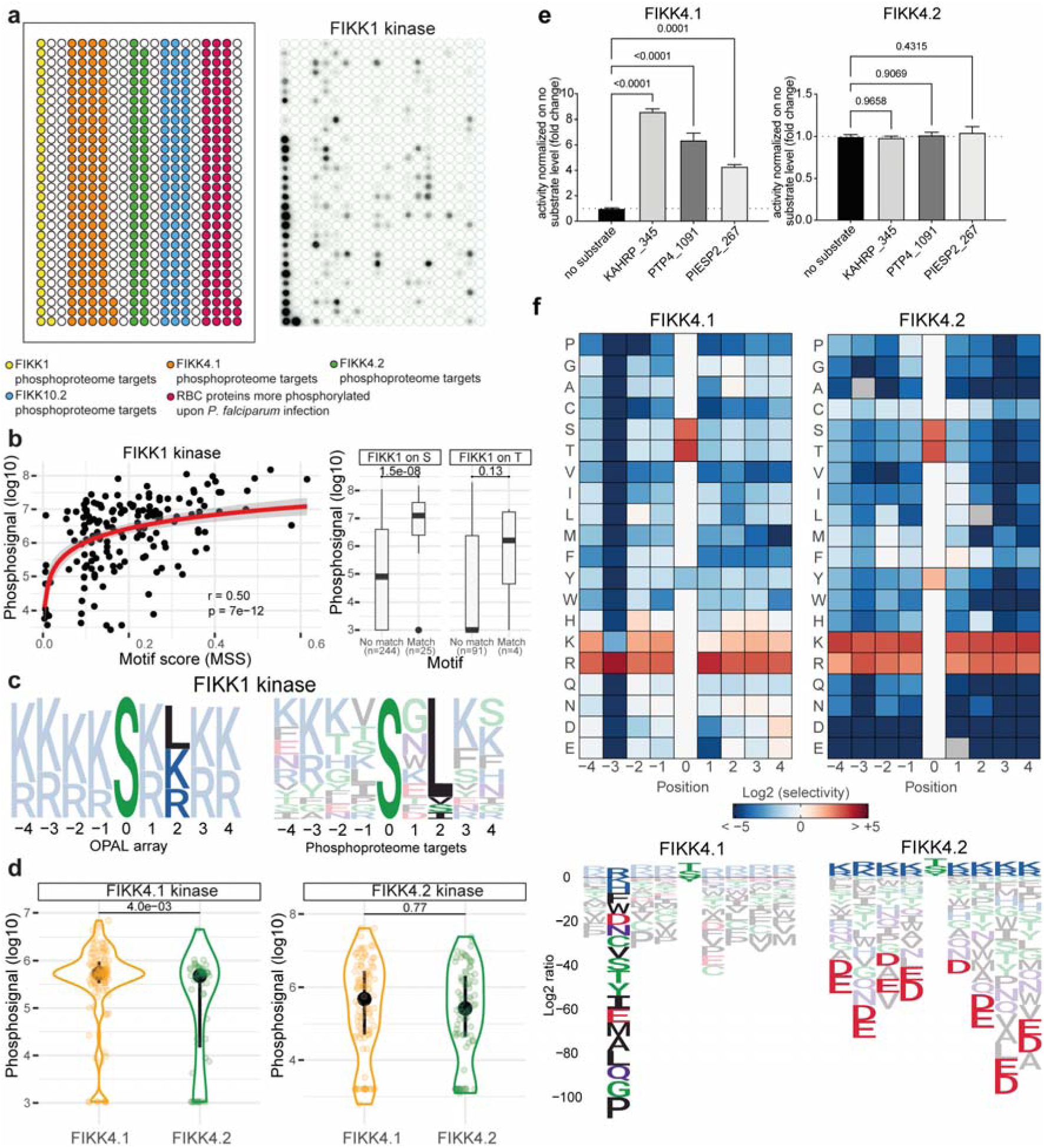
Investigation of FIKK1, FIKK4.1 and FIKK4.2 substrate specificities. **a**, Left: graphic representation of FIKK kinase phosphoproteome peptides membrane. Each dot is filled with only one peptide species. Yellow, peptides corresponding to FIKK1 targets; Orange, peptides corresponding to FIKK4.1 targets; Green, peptides corresponding to FIKK4.2 targets; Blue, peptides corresponding to FIKK10.2 targets; Red, peptides corresponding to host cell proteins found more phosphorylated upon infection by *P. falciparum*. See Supplementary Table 9 for a full list of peptides with sequences. Right: Activity of FIKK1 kinase against the phosphoproteome peptides membrane. **b,** Left: correlation of FIKK1 kinase activity on the phosphoproteome peptide membrane (log10-transformed) against the FIKK1 motif score (matrix similarity score) for each peptide (n=163). Pearson’s correlation for the y = log(x) curve. Right: Difference in FIKK1 phosphorylation signal (log10-transformed) between peptides without or with a match to the FIKK1 motif, for peptides with an S (n=269, Cohen’s D = 1.2, p=1.5e-08, Wilcoxon test, one-sided) or T phosphoacceptor (n=95, Cohen’s D = 0.57, p=0.13, Wilcoxon test, one-sided). **c,** Left: specificity logo of FIKK1 kinase for favoured amino acids, derived from the randomised OPAL peptides. Right: FIKK1 specificity logo derived from natural peptides that are phosphorylated by FIKK1 above background levels on the peptide membrane. **d,** Left: FIKK4.1 kinase activity (log10-transformed) against predicted target peptides of FIKK4.1 (orange) and of FIKK4.2 (green) (n=174, Cohen’s D = 0.57, p=4.0e-03, Wilcoxon test, one-sided). Right: FIKK4.2 kinase activity (log10-transformed) against predicted target peptides of FIKK4.1 (orange) and of FIKK4.2 (green) (n=174, Cohen’s D = 0.10, p=0.77, Wilcoxon test, one-sided). **e,** Recombinant FIKK4.1 and FIKK4.2 kinase domains activity on substrates KAHRP_345, PTP4_1091 and PIESP2_267. The results are represented as the mean±SEM fold change compared with the no substrate luminescent signal obtained using the ADP-Glo assay. Statistical significance was determined using a one-way ANOVA followed by Dunnett’s multiple comparison post-test. **f,** PWM logos for FIKK4.1 and FIKK4.2 made using data from Extended Data Fig. 3. Here, values are Log2 transformed so that a positive value depicts favoured amino acids and a negative value depicts disfavoured amino acids. See Fig. 3 caption for colour code. See Extended Data Fig. 13 for Log2 transformed PWM logos for all recombinant FIKK kinases tested.

FIKK4.1 strongly phosphorylates FIKK4.1 target peptides, while FIKK4.2 cannot clearly discriminate between FIKK4.1 and FIKK4.2 substrates (Fig. 5d). However, three peptides previously identified as FIKK4.1 substrates^16^ (KAHRP_345: GSRYS**S**FSSVN, PTP4_1091: HTRSM**S**VANTK and PIESP2_267: EIRQE**S**RTLIL) are exclusively phosphorylated by FIKK4.1 but not FIKK4.2 (Fig. 5e). Analysing disfavoured amino acids in the OPAL libraries data (Extended Data Fig. 13) reveals a notable difference between FIKK4.1 and FIKK4.2, with FIKK4.2 disfavouring negatively charged amino acids, whereas FIKK4.1 can accommodate more variety, except in position P-3 (Fig. 5f). This aligns with FIKK4.1’s strong arginine preference at P-3, a specificity determinant present in all three peptides tested in Fig. 5d. Collectively, these data show that the FIKK kinases evolved distinct phosphorylation motifs allowing the specific regulation of targets in specific subcellular contexts.

### FIKK kinase domain crystal structure informs on specificity determinant residues

We determined the crystal structure of the kinase domain of FIKK13 harbouring a mutation of the catalytic Asp-379 (D379N) to prevent autophosphorylation (Extended Data Fig. 14a) introducing microheterogeneity during production in *E. coli*. Crystallisation was facilitated by two anti-FIKK13 nanobodies generated through llama immunisation (Extended Data Fig. 14c). The kinase domain of FIKK13 was co-crystallised with the non-hydrolysable ATP-analogue ATP□S and adopts, despite the low sequence identity, the classical bi-lobal fold known from eukaryotic protein kinase (ePKs)^51^ with a few notable additions, shown and further described in Extended Data Fig. 14c. An alignment of the AlphaFold2 model and the experimental structure of the FIKK13 kinase domain revealed significant overlap [RMSD = 0.826Å] (Fig. 6a) affirming the accuracy of AlphaFold2 models, not only for FIKK13 but likely for other FIKK kinase domains.

**Fig. 6.**
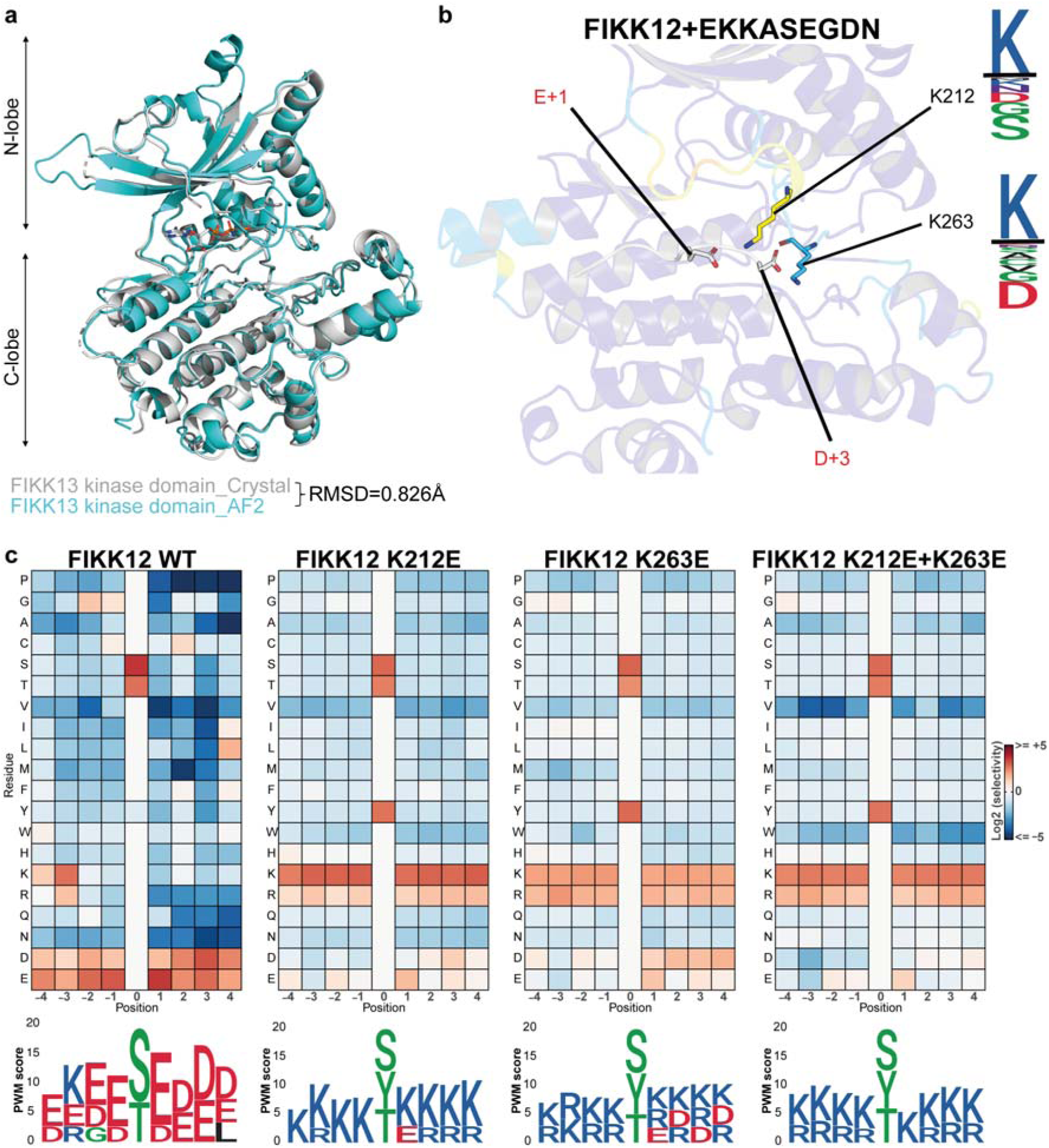
Mutating specificity determinant residues identified using FIKK13 D379N structure allows for changes in FIKK substrate specificity. **a**, Overlay of FIKK13 D379N kinase domain crystal structure with ATP□S (grey) and FIKK13 kinase domain AlphaFold2 structure prediction (cyan). Root Main Square Deviation (RMSD) was calculated using PyMol^58^. **b,** A target peptide (EKKASEGDN) of FIKK12 was modelled into the substrate-binding groove of the FIKK12 AF2 structure (see Methods). The K212 and K263 kinase residues are predicted to bind to the peptide at the +1 and +3 positions. The sequence logos show the residue conservation between FIKK12 *Plasmodium* sequences (top), and basophilic *Plasmodium* sequences (bottom). **c,** FIKK12 wild type and FIKK12 mutants phosphorylation activity on OPAL membranes represented as heatmaps (see Fig. 3bi caption). Below is represented the PWM logos (see Fig. 3bii caption).

We were unable to predict the basis for the tyrosine specificity of FIKK13 but used AlphaFold2 models for all FIKKs to predict specificity determinants. Modelling interactions of potential FIKK target peptides^16^ or preferred phosphorylation motifs (Fig. 3, Extended Data Fig. 6, 7) predicted specific residues in several FIKKs as specificity determinants: FIKK1 (E517 and E522), FIKK1 (V321), FIKK9.1 (K240), FIKK12 (K212 and K263) (Fig. 6b, Extended Data Fig. 15). To test the predictions, we reversed the charges of the amino acids (positively charged K mutated to negatively charged E and *vice versa*) or replaced the hydrophobic V321 in FIKK1 with a charged D. A single mutation in the FIKK12 kinase domain (K212E or K263E) shifted the substrate specificity from acidophilic to basophilic for all positions in the preferred phosphorylation motif (Fig. 6c). The double mutation (K212E + K263E) achieved total conversion to basophilicity. For FIKK1 and FIKK9.1, the changes in substrate specificity were more subtle with an increased overall preference for oppositely charged residues but no complete inversion (Extended Data Fig. 16a). Mutation of the V321 residue in FIKK1, homologous to K263 in FIKK12, was sufficient for the loss of the leucine specificity at P+2 for this kinase (Extended Data Fig. 16b). A similar effect could be observed for FIKK1 E517K+E522K, but not for the single mutants. Therefore, it could be that E517 and E522 combined are required for optimal positioning of the peptide leading to loss of the L+2 specificity when mutated. Thus, a single mutation in the kinase domain can dramatically change the preferred phosphorylation motif of FIKK12, and to a lesser extent that of FIKK1 and FIKK9.1. In contrast to canonical kinases, where peptide specificity is largely determined by cognate subpockets on the kinase domain^52–55^, the determinants identified here map to kinase loop regions. These loop regions are rapidly evolving (Extended Data Fig. 17) and likely flexible given their low pLDDT scores in the AF2 models^56,57^.

### Identification of pan-FIKK specific inhibitors *in vitro*

The structural analysis of the kinase domain ATP-binding site revealed some features conserved among the FIKK kinases that distinguish them from most eukaryotic kinases: 1) The glycine-loop found in ePKs, known to position ATP for catalysis, is not present in the FIKKs. An equivalent loop exists in the FIKKs, but it has a low degree of conservation among family members and is unstructured in the experimental FIKK13 structure However, a basic residue (K/R) at the position of Lys-205 is conserved throughout the FIKKs (Extended Data Fig. 18) and could help position the ATP and hence play a role in catalysis. 2) The FIKKs possess the eponymous F-I-K-K motif that plays a role in the binding of ATP, or as in this case, ATP□S. The invariant Phe-228 (Extended Data Fig. 18) is stacked upon the adenine in the back of the nucleotide-binding pocket (Extended Data Fig. 14d). The bulky and hydrophobic nature of the Phe sidechain reduces the size of the back-pocket with the equivalent residues in ePKs often having small side chains such as Ala, or in rare instances Val^61^. ATP coordination is likely supported by Lys-230, equivalent to Lys-72 in PKA^51^, which coordinates the phosphates of the nucleotide, thereby sensing nucleotide pocket occupation and forms a salt-bridge with the conserved Glu-261 on the C-helix, (Glu-91 in PKA), a hallmark of active ePKs^51^. 3) Most FIKK kinases possess a small gatekeeper residue not found in most human kinases^62^. This fundamental, and conserved, difference in nucleotide pocket composition could enable drug development specifically targeting the FIKK family of kinases.

We first tested six different Staurosporine analogues, which inhibit the majority of human kinases (>85%) by competing with ATP^63–65^ on recombinant FIKK8. None inhibited FIKK8 activity (Fig. 7a), highlighting the distinctive features of the FIKK kinase domain. A screen of the PKIS kinase inhibitor library (containing 868 ATP analogues), developed for human kinases^66,67^, identified 12 compounds that inhibited the FIKK8 kinase domain activity by =/>75% at 10µM concentration (Fig. 7b). The IC_50_s of these 12 compounds ranged between 11nM and 332nM (Supplementary Table 10). Further biochemical screening revealed structure-activity relationships (SAR) for several analogues, including some close structural analogues with weak FIKK potency that could serve as useful negative control compounds. From this compound set, three compounds were prioritised as inhibitor tools for FIKKs. GW779439X and GSK2181306A are potent FIKK inhibitors from different chemical series. GSK3184025A was selected as a very weakly active (>10µM) compound from the same chemical series as GW779439X (Extended Data Fig. 19). Most recombinant FIKK kinase domains were inhibited *in vitro* by either GW779439X or GSK2181306A (Fig. 7c), while no inhibition was observed with the inactive compounds GSK3184025A.

**Fig. 7.**
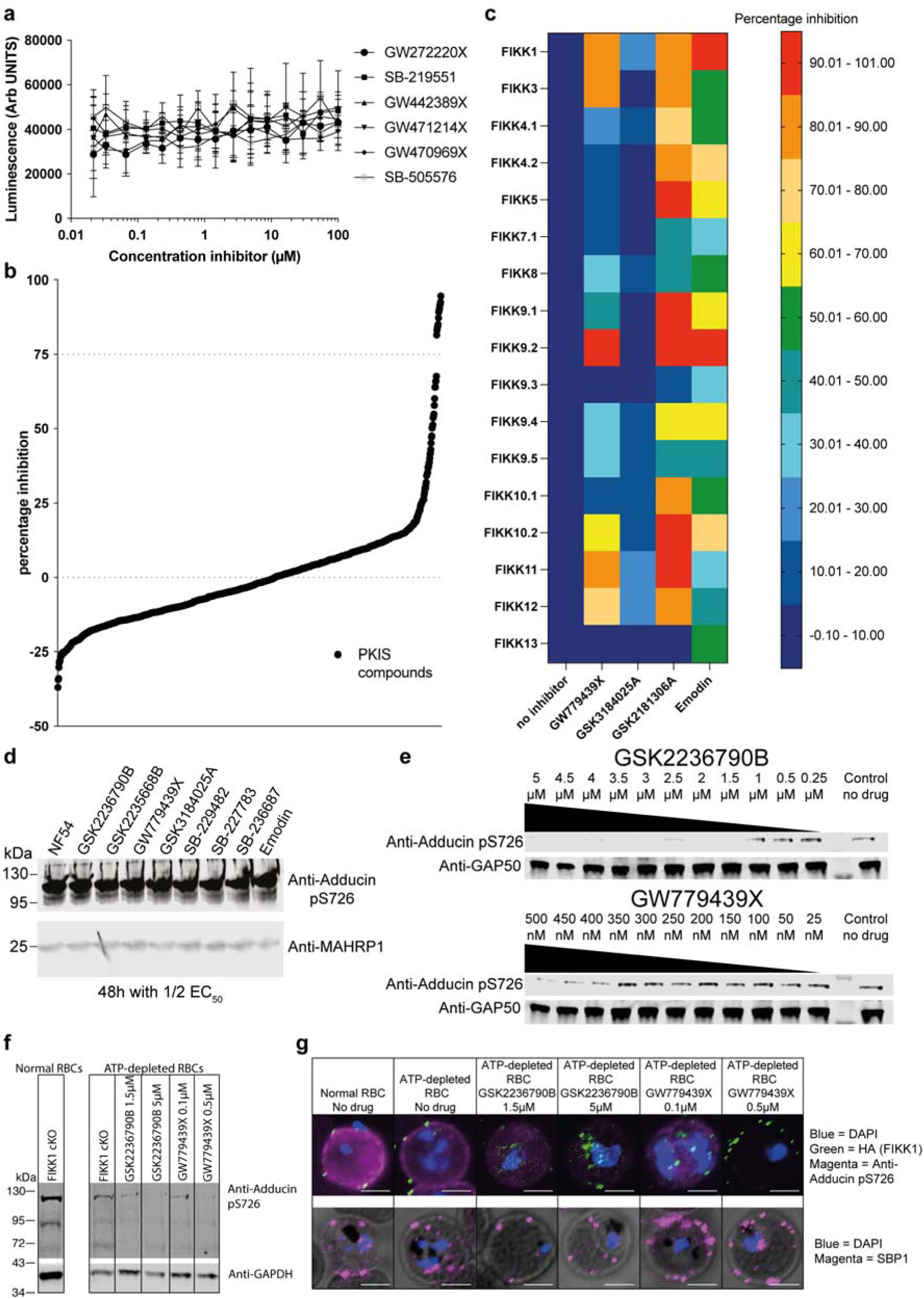
PKIS library screen allows for the identification of several pan-FIKK kinases inhibitors which target at least one FIKK kinase in ATP-depleted iRBCs. **a**, FIKK8 activity in the presence of increasing concentrations of Staurosporine analogues (GW272220X, SB-219551, GW442389X, GW471214X, GW470969X and SB-505576). n = 6 technical replicates for each inhibitor. Shown is the mean±SEM. **b,** Ranked plot showing the results of the PKIS library screen on recombinant FIKK8. A threshold of >75% inhibition was arbitrarily set and identified the 12 most potent PKIS compounds on recombinant FIKK8 kinase domain (n = 2). Each data point represents the mean percentage inhibition in both replicates. **c,** Heatmap representing inhibition (%) of selected compounds on recombinant FIKK kinase domains (n=3 biological replicates). **d,** Western blot showing adducin S726 phosphorylation in iRBCs treated with SAR-identified compounds and Emodin at 1⁄2 EC_50_ for 48 hours. The MAHRP1 antibody (bottom) demonstrates equal loading. **e,** Western blot showing adducin S726 phosphorylation in RBCs pre-treated with 1228µM iodoacetamide and 2046µM inosine, infected with wildtype NF54 *P. falciparum* and treated with different concentration of either GSK2236790B or GW778439X. The GAP50 antibody demonstrates equal loading. **f,** Western blot showing adducin S726 phosphorylation in RBCs pre-treated with 1228µM iodoacetamide and 2046µM inosine, infected with FIKK1 condKO DMSO-treated *P. falciparum* and treated with different concentration of either GSK2236790B or GW778439X. GAPDH antibody demonstrates equal loading. **g,** Immunofluorescence assays showing adducin S726 phosphorylation and protein export in ATP-depleted iRBC treated with different concentrations of either GSK2236790B or GW778439X. Protein export is investigated with anti-HA antibodies targeting the C-terminal HA-tag fused to FIKK1 kinase domain and with anti-SBP1 antibodies. DAPI (blue) is used as a nuclear staining. Scale bar = 5µm.

FIKK9.3 and FIKK13 were not inhibited by any compound. The S/T/Y kinase inhibitor Emodin, known to inhibit the *P. vivax* FIKK^24^ and *P. falciparum* FIKK8^25^ showed potency against all *Pf*FIKK kinases (Fig. 7c). These results support the feasibility of pan-FIKK inhibition and provide further support that FIKK13 is indeed a tyrosine kinase.

### Pan-FIKK inhibitors inhibit FIKK1 in cell culture

In live parasite cultures, GW779439X, GSK2177277A and GSK2181306A inhibited parasite growth [EC_50_s = 0.31±0.01µM, 0.40±0.01µM and 0.16±0.01µM respectively] (Extended Data Fig. 20a), but not phosphorylation of serine 726 on human adducin (Fig. 7d), which depends on FIKK1^16^. As none of the exported *Pf*FIKK kinases were previously found to be individually essential for parasite growth^16^, we hypothesised that the compounds engage one or multiple kinases other than the FIKKs at the concentrations used, preventing us from demonstrating their activity on the FIKKs in cell culture. Indeed, *P. knowlesi,* which only expresses one non-exported FIKK kinase not expected to be essential for parasite growth, is equally susceptible to all the compounds (Extended Data Fig. 20b).

High ATP concentrations in the RBC (2-5mM)^69^ may compete with the ATP analogues and prevent proof of concept of FIKK inhibition in cell culture. As we could not increase drug concentrations to test on-target activity without killing parasites, we reduced the ATP concentration in the RBC^70^ by pre-treating RBCs with 1228µM iodoacetamide and 2046µM inosine. This resulted in substantial ATP depletion without preventing adducin S726 phosphorylation or parasite development (Extended Data Fig. 21). Under these conditions, adducin S726 phosphorylation, but not protein export (SBP1^71^), was inhibited by the compounds (Fig. 7e, f, g), suggesting that they impair FIKK1 and potentially other FIKK kinases activity. Taken together, these data show that the compounds identified as pan-recombinant FIKK inhibitors are active on at least one of the parasite FIKK kinases in live cell culture. This sets a framework for screening better inhibitors working at physiological levels of ATP in RBCs.

## Discussion

*Plasmodium* species of the *Laverania* are thought to have evolved ∼1 million years ago from the bird-infecting *Plasmodiae*^12^. ∼50,000 years ago, *P. falciparum* emerged as a human parasite^14^, with a severe population bottleneck in the last 5,000-10,000 years^12^. Several gene families important for host-pathogen interaction evolved specifically in the *Laverania* but their function remains largely elusive. Pseudogenisation of genes within these families in different *Laverania* species suggests that some genes may be remnants of their evolutionary past or indicate a level of redundancy that relaxes selection on current gene copies. Here, we provide strong evidence that most FIKK kinases in *P. falciparum* have diversified in function, are likely essential in human infections and appear under stringent selection within the *Laverania* clade. However, a few kinases are found as pseudogenes in patient isolates, indicating these may be remnants from an ancestor not required for infection of modern humans. We observe notable differences in pseudogenisation between geographical backgrounds, suggesting that the environment might impact FIKK relevance. This is interesting in the light of a recent study which found an association between a SNP in the *fikk4.2* gene and sickle-cell trait which protects from severe *P. falciparum* malaria and is highly prevalent in people of African descent^72^.

The expansion and diversification of the FIKK kinase family into several members with likely different functions required specialisation of each kinase. This was achieved by different expression timing, subcellular localisation and, as we show here, the evolution of highly specific phosphorylation motifs. By combining molecular docking and mutational analyses, we identified strong specificity determinants for FIKK12, and residues with more moderate effects for FIKK1 and FIKK9.1. These map to rapidly evolving loop regions on the kinase domain, perhaps explaining why the FIKK family was able to functionally diversify rapidly (∼1 million years) in terms of its phosphorylation motif specificity.

Strikingly, we show that FIKK13 is a bona fide tyrosine kinase in *P. falciparum* and *P. gaboni*, and therefore probably in other *Laverania* species. Additionally, several FIKK kinases show dual specificity. This suggests that tyrosine phosphorylation of host, and/ or exported parasite proteins by *Laverania* secreted kinases is not only carried out by hijacking human kinases, as believed so far^73,74^. Thus, some exported FIKK kinases have likely evolved to specifically interfere with critical host signalling pathways that rely on tyrosine phosphorylation. This could be specifically important for the infection of nucleated cells such as erythroid precursors^75^ and/or liver cells^76^. While a secreted dual specificity kinases (S/T and Y) has been described in the related *Toxoplasma* parasite^77^, the evolution of apparently exclusive tyrosine kinase specificity in FIKK13 from a S/T kinase family has not previously been observed. This is an important finding as it implies that in other species, bona fide tyrosine kinases might have evolved from a recent S/T kinase ancestor, similar to what is observed here for the FIKK family. However, predicting tyrosine kinase activity solely based on sequence or structure remains elusive. The determinants of tyrosine specificity appear to be more difficult to determine than for canonical kinases and it is a remaining challenge to combine computational and experimental approaches to understand the precise molecular relationship between kinase sequence and specificity for all FIKKs and all substrate positions.

While the diversification of the substrate specificity is underpinned by evolution of the peptide binding area, several conserved features of the FIKK kinases required for ATP-binding may allow the generation of inhibitors which target several or all FIKKs simultaneously. The Phe in the F-I-K-K motif that is strictly conserved across the FIKK kinases, appears to be involved in an unusual coordination of ATP in the kinase active site. In combination with a small gatekeeper residue common across the FIKKs, this feature may allow the design of compounds that specifically inhibit the FIKKs. Here, we identify *in vitro* pan-specific FIKK inhibitors that can interfere with FIKK1 activity in live parasites upon reducing ATP-levels in the RBC, demonstrating that pan-FIKK inhibition is an achievable goal, if inhibitors more specific over human enzymes can be found. Since FIKK kinases are with a high likelihood critical for parasite survival in the host, their collective inhibition represents an interesting strategy for combination therapies. Resistance through mutations in single genes is readily observed against most current drugs^78^, which would not easily be possible for compounds that inhibit a whole family of proteins. The crystal structure solved in this work will allow further investigation of the FIKK family chemical inhibition.

## Extended Data Figures List

**Extended Data Fig. 1**. Phylogenetic tree of *Pf*FIKK kinases rooted on FIKK8 sequences.

**Extended Data Fig. 2**. CRISPR/Cas9 strategy to generate FIKK::TurboID fusion proteins and validation.

**Extended Data Fig. 3**. Alignment of *P. falciparum* FIKK protein sequences allows for accurate determination of the FIKK kinase domain starting amino acid.

**Extended Data Fig. 4**. Coomassie-stained gel of purified recombinant FIKK kinase domains.

**Extended Data Fig. 5**. FIKK8 OPAL membrane.

**Extended Data Fig. 6**. Basophilic FIKK kinases preferred phosphorylation motifs.

**Extended Data Fig. 7**. Acidophilic FIKK kinases preferred phosphorylation motifs.

**Extended Data Fig. 8**. FIKK1, FIKK9.1 and FIKK13 OPAL membranes.

**Extended Data Fig. 9**. Identification of a tyrosine-based cyclic peptide as a substrate for FIKK13.

**Extended Data Fig. 10**. Heat map representation of OPAL arrays raw data for *P. gaboni* FIKK1, FIKK9.1 and FIKK13.

**Extended Data Fig. 11**. Protein sequence identity matrix of *P. falciparum* FIKK kinases.

**Extended Data Fig. 12**.FIKK4.1, FIKK4.2 and FIKK10.2 activity on the phosphoproteome peptides libraries.

**Extended Data Fig. 13**. Log2 transformed PWM logos for all recombinant FIKK kinases tested.

**Extended Data Fig. 14**. FIKK13 D379N dead mutant crystal structure informs on ATP binding.

**Extended Data Fig. 15**. Target peptides of FIKK1, FIKK9.1, or FIKK12 modelled into the substrate-binding groove of the FIKK AF2 structures.

**Extended Data Fig. 16**. Substrate specificity assessment of FIKK1 and FIKK9.1 kinase mutants using OPAL arrays.

**Extended Data Fig. 17**. Sequence conservation of FIKK specificity determinants.

**Extended Data Fig. 18**. Multiple sequence alignment of various kinase domains.

**Extended Data Fig. 19**. Structure-Activity Relationship assay identifies closely related compounds with different behaviours towards recombinant FIKK8 kinase domain.

**Extended Data Fig. 20**. The three most potent *in vitro* FIKK inhibitors kill *Plasmodium* parasites in culture.

**Extended Data Fig. 21**. Optimisation of ATP-depletion conditions.

## Supplementary Tables List

**Supplementary Table 1**. FIKK orthologues in *Laverania*.

**Supplementary Table 2**. FIKK transcript levels during *P.falciparum* asexual replication cycle.

**Supplementary Table 3**. STOP codon in *fikk* genes in 2085 field isolate genomes.

**Supplementary Table 4**. *fikk* genes deletions in 2085 field isolate genomes.

**Supplementary Table 5**. Transcription evidence for PfFIKK kinases in Gametocytes and mosquito stages according to Malaria Cell Atlas data.

**Supplementary Table 6**. Table of proteins in the vicinity of FIKK4.1 and/or FIKK4.2 identified by TurboID-based proximity labelling.

**Supplementary Table 7**. Start sites of recombinantly expressed FIKK kinase domains.

**Supplementary Table 8**. SPR binding affinities for FIKK13 peptides.

**Supplementary Table 9**. Phosphoproteome peptide library composition.

**Supplementary Table 10**. Half maximal inhibitory concentration (IC50) of the PKIS compounds identified as inhibitors of FIKK8 recombinant kinase domain.

**Supplementary Table 11**. Material used for generation of FIKK_TurboID parasite lines.

**Supplementary Table 12**. Recodonised sequences used for FIKK kinase domains expression in *E. coli*

**Supplementary Table 13**. Processed mass spectrometry proximity labelling data.

**Supplementary Table 14**. FIKK13 RaPID selection Next Generation Sequencing Data.

**Supplementary Table 15**. Flow cytometry data for EC50s determination of FIKK inhibitors in *P. falciparum* and *P. knowlesi*.

**Supplementary Table 16**. Flow cytometry data for ATP-depletion optimisation experiment.

## Material and Methods

### FIKK orthologues in *Laverania*

To assess the number of FIKK orthologues in each *Laverania* species, the word “FIKK” was entered into the search engine of the PlasmoDB website (www.PlasmoDB.org) (release 66) selecting *Plasmodium adleri* G01, *Plasmodium billcollinsi* G01, *Plasmodium blacklocki* G01, *Plasmodium falciparum* 3D7, *Plasmodium gaboni* SY75, *Plasmodium praefalciparum* G01 and *Plasmodium reichenowi* CDC genomes. To assess syntenicity, the JBrowse genome browser of PlasmoDB was used, selecting the “Syntenic Sequences and Genes (Shaded by Orthology)” track. The *P. falciparum* 3D7 genome was used as a reference to evaluate whether the chromosome sequences from the other *Laverania* species were complete. ‘no genome information’ signifies that chromosome sequence was not available in the database, probably due to degradation of telomeric regions.

### Field Genomes FIKK Pseudogenisation analysis

To identify genetic variants in *fikk* genes, a global dataset of clinical *P. falciparum* samples was examined, using the Pf3K project release 5 (www.malariagen.net/projects/parasite.pf3k)^79^. Out of the 2483 *P. falciparum* clinical samples of diverse geographical origin, 2085 with high quality data were selected (>80% of the genome covered with 10 or more reads). For each isolate genome, *fikk* pseudogenes were defined by the presence of at least one internal STOP codon variant with an alternative allele frequency greater than 0.5 (Alt reads divided by total number of reads of that position). To identify natural genomic deletions that include *fikk* genes, deleted genes were defined as 95% of the gene sequence with coverage under 3 reads. The large majority of deleted FIKK kinases had zero reads over the entire length of the gene, with the rest of the genome being over 10X coverage (typically ∼50X). As a complementary approach, we made use of the microarray transcriptomic data from Mok *et al.*^80^. From the 2085 genome samples, 693 transcriptomes from the same isolates were also available. A negative Log2 value was defined as ‘no expression’.

### FIKK percentage of expression heatmap

Expression data were taken from Hoeijmakers *et al.* RNA-sequencing dataset available on PlasmoDB release 66 (www.PlasmoDB.org). The dataset gives a TPM (Transcript Per kilobase Millions) value for eight different time windows throughout *P. falciparum* 48 hours asexual replication cycle ([40-5hpi]; [2-10hpi]; [7-15hpi]; [12-20hpi]; [17-25hpi]; [22-30hpi]; [27-35hpi]; [32-40hpi]). Percentage of expression was calculated for each *P. falciparum* FIKK kinase using the following formula 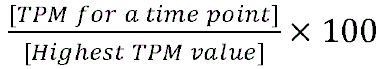. Percentage of expression values were then plotted in GraphPad PRISM10 and represented as a heatmap with dark blue cells representing no expression and yellow cells representing 100% expression.

### Human cells

Human RBCs were acquired from the National Health Service Blood and Transplant (NHSBT) service.

### In vitro maintenance and synchronisation of *Plasmodium* parasites

Human erythrocytes infected with *P. falciparum* asexual stages were cultured at 37°C in complete medium. Complete medium consists of 1L RPMI-1640 medium supplemented with 5g Albumax II (ThermoFischer Scientific) to act as a serum substitute, 0.292g L-glutamine, 0.05g hypoxanthine, 2.3g sodium bicarbonate, 0.025g gentamycin, 5.957g HEPES and 4g dextrose. A haematocrit of 1-5% was used and the blood was from anonymous donors provided through the UK Blood and Transfusion service. According to standard procedures, parasites were grown in a gas atmosphere consisting of 90% N_2_, 5% CO_2_ and 5% O ^81^. Thin blood smear fixed in 100% methanol, air-dried and stained with Giemsa were routinely used to assess parasitemia and developmental stages by light microscopy. *P. knowlesi* parasites in the asexual RBC stages were cultured in complete medium supplemented with 10% human serum as described previously^82^. Parasite cultures were synchronised by Percoll (GE Healthcare) for isolation of mature schizont stages parasites. Purified schizonts were incubated in complete medium at 37°C with fresh RBCs for 4 hours in a shaking incubator. Any remaining schizonts were removed with a second Percoll purification leaving only tightly synchronised ring-stage parasites in the flask.

### Gametocyte induction, culture, FIKK gene excision and harvest

An adapted version of previously described techniques was used to obtain synchronous gametocytes^83^. Briefly, highly synchronous ring-stage parasites at 8-10% parasitemia were stressed by retaining half the spent culture medium and replenishing the rest with fresh complete medium. The following day, the stressed cultures were spun and the spent culture medium was replaced with complete media. Cultures were left shaking until the following day when all the schizonts had ruptured and reinvaded. A certain proportion of the reinvaded rings should have then committed to gametocytogenesis. This committed parasites were then split into two flasks and treated for 4 hours at 37°C with either 100nM rapamycin (Sigma) or dimethyl sulfoxide (DMSO) (0.1% [vol/vol]) as described previously^84^. Parasites were then washed three times with complete medium and cultured in complete medium supplemented with 10% human serum. From this point onwards, parasite culture medium was exchanged daily with pre-warmed complete medium supplemented with 10% human serum and heparin at 20 units/ml to prevent asexual growth. When a majority of stage III gametocytes could be seen on Giemsa smears, the cultures were submitted to Percoll purification allowing isolation of sexual stages which were lysed in 5x SDS-sample buffer for Western Blot analysis of FIKK kinases expression.

### Immunoblotting

Parasites submitted to Western Blot analysis were first harvested by Percoll purification. 1µl of parasite pellets were then resuspended in 15µl PBS, lysed with 5x SDS-sample buffer (25mM TrisHCl pH 6.8, 10% SDS, 30% Glycerol, 5% β-mercaptoethanol, 0.02% bromophenol blue) and denatured at 95°C for 5 minutes. Samples were then subjected to SDS-PAGE, transferred onto Transblot Turbo™ Mini-size nitrocellulose membrane (Biorad) and blocked overnight in 5% skimmed milk in PBS with 0.2% Tween-20 at 4°C. For FIKK kinases expression in gametocytes (Fig. 1c), the membranes were probed with rat anti-HA high affinity (clone 3F10, Roche, 1:1,000) and rabbit anti-GAP50^85^ (a gift from Julian Rayner, 1:2,000) antibodies. For Western blots investigating Adducin S726 phosphorylation (Fig. 7 and Extended Data Fig. 21), the membranes were probed with rabbit anti-Adducin pS726 (Abcam, 1:1,500), rabbit anti-MAHRP1 (a gift from J. Rayner and L. Parish; 1:2,000), rabbit anti-GAP50 (1:2,000) or mouse anti-GAPDH (1:10,000) (Monoclonal antibody 7.2 (anti-GAPDH) which was obtained from The European Malaria Reagent Depository (http://www.malariaresearch.eu). Source: Dr. Jana McBride^86^). For Western blots assessing protein biotinylation by FIKK4.1 and FIKK4.2::TurboID (Extended Data Fig. 2b), the membranes were probed with rabbit anti-MAHRP1 (1:2,000) and mouse anti-V5 (Abcam, 1:1,000). Following primary antibody staining, the membranes were incubated with the relevant secondary fluorochrome-conjugated antibodies (LI-COR, 1:20,000) or IRDye® 800CW Streptavidin (1:2,000). The antibody reactions were carried out in 5% skimmed milk in PBS with 0.2% Tween-20 for 1 hour in the dark and membranes were washed 3 times between each antibody staining in PBS with 0.2% Tween-20. After a final wash with PBS, the antigen-antibody reactions were visualised using the Odyssey infrared imaging system (LI-COR Biosciences).

### Transcription evidence of *Pf*FIKK kinases in sexual and mosquito stages

Data were obtained from the Malaria Cell Atlas (www.malariacellatlas.org)^29^. The SmartSeq2 cell view was used and an FIKK kinase was considered expressed if at least two sample analysed showed an expression above 0.

### Generation of FIKK::TurboID parasite lines

FIKK::TurboID parasite lines were generated using CRISPR/Cas9. Briefly, suitable gRNAs for FIKK4.1 and FIKK4.2 were identified using the Eukaryotic Pathogen CRISPR guide RNA/DNA Design Tool (grna.ctegd.uga.edu)^87^. A pair of complementary oligonucleotides corresponding to the 19 nucleotides closest to the identified PAM sequence was synthesised (IDT), phosphorylated using T4 polynucleotide kinase, annealed and ligated into pDC_Cas9_hDHFRyFCU^32^ digested with BbsI. To generate compatible, sticky ends between the annealed primer pairs encoding the gRNAs and the BbsI digested vector, the forward oligonucleotide had 5’-ATTG added to the 19 nucleotides corresponding to the gRNAs whereas the compatible oligonucleotide had a 5’-AAAC overhang added (See Supplementary Table 11). This way, gRNAs targeting *fikk4.1* and *fikk4.2* genes were assembled using oligonucleotide pairs gRNA_4.1_310For/Rev and gRNA_4.2_235For/Rev respectively. Repair templates containing a 5’HR, a recodonised sequence, a linker, a TurboID-coding sequence, a V5-tag and a 3’HR flanked by two XhoI restriction sites were ordered from GeneArt (Supplementary Table 11). For transfection, 60µg repair template plasmid was linearised with XhoI for 4 hours at 37°C before inactivation at 80°C for 20 minutes. 20µg of gRNA plasmid was added and the plasmid mixture was ethanol precipitated, washed and resuspended in 10µl sterile TE buffer (10mM Tris, 1mM EDTA). In parallel, highly synchronised segmented schizonts (48 h.p.i) of NF54::DiCre parasites^88^ were collected by Percoll-enrichment and washed once with complete medium. The DNA constructs in TE buffer were mixed with 90µl P3 Primary cell solution (Lonza) and used to resuspend 20µl segmented schizonts which were subsequently transferred to a transfection cuvette. Transfections were performed by electroporation using the FP158 programme from an Amaxa 4D Electroporator machine (Lonza). Following transfection, the parasites were transferred to pre-warmed flasks containing 2ml complete medium and 300µl fresh uRBCs. After 40 minutes of gentle shaking at 37°C, 8 ml complete medium were added to the flask. Transfected parasites were incubated for 24 hours, then selection was performed with 2.5nM WR99210 (Jacobus Pharmaceuticals) for four days. Following establishment of the transgenic lines, correct modification of the parasite genome was confirmed by PCR using the primers described in Extended Data Fig. 2 and Supplementary Table 11.

### Phylogenetic tree

All FIKK amino acid sequences were retrieved from the UniProtKB^89^. Heavily truncated sequences (<200 amino acids) were removed manually. The full-length protein sequences were then aligned using the MAFFT L-INS-i algorithm^90^. Alignment positions where more than 20% of sequences contain a gap (-gt 0.8) were removed from the multiple sequence alignment (MSA) using trimAl software^91^. A maximum likelihood (ML) estimate of the FIKK phylogeny was generated with IQ-TREE2 software^92^, using the ModelFinder parameter (-m MFP) to automatically detect the best evolutionary model^93^. Branch support for the ML phylogeny was assessed using 100 replicates of the Felsenstein bootstrap (-b 100)^50^. The phylogenetic tree was visualised using the ggtree package in R^94^ after removing homologues to the FIKK 7.2 and FIKK14 pseudogenes.

### Immunofluorescence assays

Air-dried blood films were fixed for 5 min in ice-cold methanol and subsequently rehydrated in PBS for 5 min. Slides were blocked in 3% (w/v) bovine serum albumin (BSA) in PBS containing kanamycin (50µg/ml) for 1 h and subsequently incubated with primary antibodies in 1% (w/v) BSA in PBS containing kanamycin (50µg/ml) for 1 h at room temperature. Primary antibodies dilutions were as follow: high affinity rat anti-HA (clone 3F10, Roche; 1:1,000), mouse anti-FIKK4.2 (1:1,000) (Monoclonal antibody 126 (anti-FIKK4.2) was obtained from The European Malaria Reagent Depository (http://www.malariaresearch.eu). Source: Dr. Odile Mercereau-Puijalon^17^), rabbit anti-phosphoAdducin S726 (1:1,500) (Abcam), rabbit anti-SBP1 (1:10,000) (gift from T. Spielmann^95^). After three washes with PBS, the coverslips were incubated with the relevant Alexa Fluor secondary antibodies (1:2,000 in PBS with 1% BSA) at room temperature for 1 h in the dark. After three final washes with PBS, the slides were mounted with Prolong Gold antifade reagent (Invitrogen) containing the DNA dye 4□, 6-diamidino-2-phenylindole (DAPI), covered with a coverslip and sealed with nail polish. Images were taken using a Ti-E Nikon microscope using a ×100 TIRF objective at room temperature equipped with an LED-illumination and Orca-Flash4 camera. The images were processed using Nikon Elements software (Nikon).

### Proximity labelling experiments

We first performed a NF54-FIKK4.2::TurboID comparison at the peptide level. We then repeated the assay with NF54, FIKK4.1::TurboID and FIKK4.2::TurboID. Both experiments were performed following the same protocol and data from both experiments were combined.

#### Cell culture and lysis

For all experiments, NF54 WT parasites, used as controls, FIKK4.1::TurboID and FIKK4.2::TurboID parasites were tightly synchronized to a 4-hours window using Percoll. For each line, parasites were grown in biological triplicate in 200ml of complete medium containing biotin (0.2mg/L Η 819nM) at at least 10% parasitaemia in 2ml of blood, each replicate being cultured in blood coming from different donors. iRBCs were harvested at late schizont stage (44-48 hpi) using Percoll. Subsequently, parasites were washed 3 times with 50ml of complete medium and 5 times with 5ml PBS. Parasites were then lysed in 8M urea in 50mM HEPES pH8.0 containing protease inhibitors (cOmplete, Roche). Samples were further solubilised by sonication with a microtip sonicator on ice for 3 rounds of 30 seconds at an amplitude of 30%. Lysates were then clarified by centrifugation at 15,000 rpm for 30 minutes at 4°C. The protein concentrations were then calculated using a BCA protein assay kit (Pierce), first diluting 20µl aliquots from all lysates 1:25 in H_2_O to reduce the concentration of urea and then following the instructions provided in the kit.

#### Protein digestion

4mg of each lysate was then reduced with 5mM dithiothreitol (DTT) for 1 hour at room temperature and subsequently alkylated in the dark with 10mM iodoacetamide for 30 minutes at room temperature. Following alkylation, the lysates were diluted with 50mM HEPES pH8.0 to <2M urea and digested overnight with trypsin (Promega) at 1:50 (enzyme:protein) at 37°C.

#### Sep-Pak desalting

Samples were cooled on ice for 10 minutes before being acidified with trifluoroacetic acid (TFA; ThermoFischer Scientific) to a final concentration of 0.4% (vol/vol) and left on ice for 10 more minutes. All insoluble material was removed by centrifugation (15,000 rpm, 10 minutes, 4°C) and the supernatants were desalted on Sep-Pak C18 1cc Vac cartridges (Waters) in conjunction with a vacuum manifold. The columns were first washed with 3ml acetonitrile, conditioned with 1ml of 50% acetonitrile and 0.5% acetic acid in H_2_O, and then equilibrated with 3ml of 0.1% TFA in H_2_O. The acidified samples were loaded, desalted with 3ml of 0.1% TFA in H_2_O, washed with 1ml of 0.5% acetic acid in H_2_O and finally eluted in 1.3ml of 50% acetonitrile and 0.5% acetic acid in H_2_O. Each sample was then dried by vacuum centrifugation.

#### Charging protein G agarose beads with anti-biotin antibodies

60µl of protein G agarose bead slurry (ThermoFischer Scientific) were taken per sample. Beads were washed three times with 10 bead volumes of Biosite buffer^96^ (50mM Tris-HCl pH8.0, 150mM NaCl, 0.5% Triton x100, pH7.2-7.5) at 4°C. According to supplier recommendations, protein G agarose beads were functionalised with 100µg antibodies / 100µl slurry with two different anti-biotin antibodies (150-109A, Bethyl Laboratories; ab53494, Abcam) by adding 300µg of each antibody to the beads which were incubated rotating overnight at 4°C.

#### Immunoprecipitation

Samples were dissolved in 1.5ml Biosite buffer on ice and pH adjusted with 1-5µl 10M NaOH to 7-7.5 at 4°C. Any undissolved material was removed by spinning at 15,000rpm, 10 minutes, 4°C and peptide BCA assay (Pierce) was performed on the supernatant to know the peptide concentration in each sample. Protein G agarose beads functionalised with anti-biotin antibodies were washed three times with 10 bead volumes (3ml) Biosite buffer and equal amount of peptides per sample was added onto the antibody loaded beads (60µl slurry per sample). The mixture was incubated rotating for 2 hours at 4°C. Beads were pelleted at 1,500xg for 2 minutes at 4°C and washed three times with 500µl Biosite buffer, once with 500µl 50mM Tris-HCl pH8.0 and three times with 500µl H_2_O. Peptides were eluted from the beads by adding 50µl of 0.2% TFA, gently shaken and spun at 1,500xg for 2 minutes at 4°C. Elution was repeated 4 times for a total volume of 200µl.

#### Stage-tip desalting

All samples were desalted before LC-MS/MS using Empore C18 discs (3M). Briefly, each stage-tip was packed with one C18 disc, conditioned with 100µl of 100% methanol, followed by 200µl of 1% TFA. The samples were loaded onto the stage-tip in 200µl of 0.2% TFA, washed twice with 300µl of 1% TFA and eluted with 40µl of 40% acetonitrile + 0.1% TFA. The desalted peptides were vacuum dried in preparation for LC-MS/MS analysis.

#### LC-MS/MS

Samples were loaded onto Evotips according to manufacturer’s instructions. After a wash with 0.1% formic acid in H_2_O, samples were loaded onto an Evosep One system coupled to an Orbitrap Fusion Lumos (ThermoFisher Scientific). A PepSep 15cm column was fitted onto the Evosep One and a predefined gradient for a 44 minutes method was used. The Orbitrap Fusion Lumos was operated in data-dependent mode with a 1 second cycle time, acquiring IT HCD MS/MS scans in rapid mode after an OT MS1 survey scan (R=60,000). The target used for MS1 was 4E5 ions whereas MS2 target was 1E4 ions. The maximum ion injection time utilised for MS2 scans was 300ms, the HCD normalised collision energy was set at 32 and the dynamic exclusion was set at 15 seconds.

#### Data processing

Acquired raw files were processed with MaxQuant v1.5.2.8^97^.

The Andromeda^98^ search engine was used to identify peptides from the MS/MS spectra against *Plasmodium falciparum* (PlasmoDB_v46^13^) and *Homo sapiens* (UniProt, 2020^89^). Acetyl (Protein N-term), Biotin (K), Oxidation (M) were selected as variable modifications whereas Carbamidomethyl (C) was selected as a fixed modification. The enzyme specificity was set to Trypsin with a maximum of 3 missed cleavages. Minimum peptide length was set to 6 amino acids. Biotinylated peptides search in MaxQuant was enabled by defining a biotin adduct (+226.0776) on lysine residues as well as three diagnostic ions: fragmented biotin (m/z 227.0849), immonium ion harbouring biotin with a loss of NH_3_ (m/z 310.1584) and an immonium ion harbouring biotin (m/z 327.1849).

The precursor mass tolerance was set to 20ppm for the first search (used for mass re-calibration) and to 4.5ppm for the main search. The datasets were filtered on posterior error probability (PEP) to achieve a 1% false discovery rate on protein, peptide and site level. Other parameters were used as pre-set in the software. ‘Unique and razor peptides’ mode was selected to allow identification and quantification of protein in groups (razor peptides are uniquely assigned to protein groups and not to individual proteins). Intensity-based absolute quantification (iBAQ) in MaxQuant was performed using a built-in quantification algorithm^97^ enabling the ‘Match between runs’ option (time window 0.7 minutes) within replicates.

#### Data analysis

The MaxQuant output files were processed with Perseus v1.5.0.9^99^.

Modified peptides data were filtered to remove contaminants and IDs originating from reverse decoy sequences. iBAQ intensities were log2 transformed and peptides with less than one valid value in total were removed. Non-biotinylated peptides (background) were also removed from the datasets. Additionally, peptides with intensities only in the NF54 samples were removed as they are likely to represent background binding to the beads. Replicates were grouped for each condition (NF54 and FIKK4.2::TurboID for the first experiment and NF54, FIKK4.1::TurboID and FIKK4.2::TurboID for the second experiment) and only peptides with at least two valid values in at least one group were conserved for further analysis. Data for the first experiment (NF54 – FIKK4.2::TurboID) and the second experiment (NF54 – FIKK4.1::TurboID – FIKK4.2::TurboID) are available in Supplementary Table 13.

### Network

The network representation of the TurboID data (Fig. 2c) was generated using Cytoscape v3.10.1^100^. Proximal proteins were included in the network if they contained at least one peptide that was biotinylated in 2 or more of the 3 biological replicates from either the FIKK4.1 or FIKK4.2 TurboID assays. All proteins in the vicinity of FIKK4.1 or FIKK4.2 were annotated as potential kinase targets if they were found to be less phosphorylated upon knock-out (KO) of the respective kinase, using data published in ^16^. Regulated phosphosites on candidate substrates were scored against the FIKK 4.1 or FIKK4.2 kinase specificity models presented in Extended Data Fig. 6, using a simple scoring function that outputs a normalised summation between 0 (minimum) and 1 (maximum)^101^. Data on protein proximity, target status, and motif scores are given in Supplementary Table 6.

### Recombinant protein expression and purification

The DNA sequences coding for *P. falciparum* 3D7 and *P. gaboni* SY75 FIKK kinase domains were obtained from PlasmoDB (https://plasmodb.org/plasmo/)^13^ and were codon optimised for *E. coli* expression (IDT) (https://eu.idtdna.com/CodonOpt) (see Supplementary Table 12 for recodonised FIKK kinase sequences). For FIKK4.2, blocks of low complexity repeat sequences and the short low complexity downstream sequence (amino acids 403-928) were removed as per ^17^. Sequences were subsequently inserted into a pET-28a vector (Novagen) to produce a N-terminal thrombin cleavage His_6_ tag fusion (MGSS**HHHHHH**SSG<u>LVPRGS</u>H*MASMTGGQQMG*RGS, where the sequence in bold is the His_6_ tag, the underlined sequence is the thrombin site and the sequence in italics is the T7 tag). The insert sequence was verified by DNA sequencing. For expression in *E. coli*, BL21-Gold (DE3) cells (Stratagene) were transformed with pET-28a-FIKK vectors, grown over 2 days at 18°C in ZYM-5052 media supplemented with 50µg.ml^-1^ kanamycin and harvested by centrifugation. In a typical preparation, 10g of cells were resuspended in 100ml lysis buffer (50mM Tris-HCl pH 7.5, 500mM NaCl, 1mM TCEP, 20mM imidazole, 10mM MgSO_4_, 10% glycerol and 2 protease inhibitor cocktail tablets (cOmplete, EDTA free, Roche)), lysed by sonication and clarified by centrifugation at 20,000g for 30 min at 4°C. The supernatant was loaded into a 1ml HisTrap column (GE Healthcare) and the bound proteins were eluted in 50mM Tris-HCL pH 7.5, 500mM NaCl, 1mM TCEP, 300mM imidazole and 10% glycerol. After concentration, the samples were loaded on a Hi-Load Superdex 200 16/600 column (GE Healthcare) equilibrated with 50mM Tris-HCl pH7.5, 250mM NaCl, 1mM TCEP and 10% glycerol. The fractions containing the different recombinant FIKK kinase domains were analysed by SDS-PAGE stained by Coomassie.

### Peptides arrays

Oriented Peptide Array Libraries (OPAL) and phosphoproteome peptide libraries synthesis was performed by the Francis Crick Institute Peptide Chemistry Science Technology platform as described previously^36,102^. Briefly, peptide arrays were synthesised on an Intavis ResResSL automated peptide synthesiser (Intavis Bioanalytical Instruments, Germany) by cycles of N(a)-Fmoc amino acids coupling via activation of the carboxylic acid groups with diisopropylcarbodiimide in the presence of ethylciano-(hydroxyamino)-acetate (Oxyma pure) followed by removal of the temporary α-amino protecting group by piperidine treatment. Subsequent to chain assembly, side chain protection groups are removed by treatment of membranes with a deprotection cocktail (20ml 95% trifluoroacetic acid, 3% triisopropylsilane and 2% H_2_O) for 4 hours at room temperature, them washing (4x dichloromethane, 4x ethanol, 2x H_2_O and 1x ethanol) prior to being air dried. For the phosphoproteome peptide libraries, the final product is a cellulose membrane containing a library of 11-mer peptides. Sequences of the peptides can be found in Supplementary Table 9. For the OPAL libraries, the final product is a cellulose membrane containing a library of 9-mer peptides with the general sequences: A-X-X-X-X-S-X-X-X-X-A; A-X-X-X-X-T-X-X-X-X-A or A-X-X-X-X-Y-X-X-X-X-A. For each peptide, one of the 20 naturally occurring proteogenic amino acids was fixed at each of the 8 positions surrounding the phosphorylated residue (S, T or Y), with the remaining positions, represented by X, degenerate (approximately equimolar amount of the 16 amino acids excluding cysteine, serine, threonine and tyrosine). Cellulose membranes were placed in an incubation trough and moisten with 5 ml ethanol. They were subsequently washed twice with 50ml kinase buffer (20mM MOPS, 10mM magnesium chloride and 10mM manganese chloride, pH7.4, Alfa Aesar) and incubated overnight in 100ml reaction buffer (kinase buffer + 0.2mg/ml BSA (BSA Fraction V, Sigma) + 50□g/ml kanamycin). The next day, the kinase buffer was removed and the membranes were incubated at 30°C for 1 hour in 30ml blocking buffer (kinase buffer + 1mg/ml BSA + 50μg/ml kanamycin). After incubation, the blocking buffer was replaced with 30ml reaction buffer supplemented with 300μl 10mM ATP and 125μCi [γ-32P]-ATP (Hartmann Analytics, Germany). The reaction was started by adding 100nM of the recombinant FIKK kinase domain studied and left to incubate for 20 min at 30°C with gentle agitation. After incubation, the reaction buffer was removed and the membranes were washed 10 x 15 min with 100ml 1M NaCl, 3 x 5 min with 100ml H_2_O, 3 x 15 min with 5% H_3_PO_4_, 3 x 5 min with 100ml H_2_O and 2 x 2 min with 100ml ethanol. The membranes were left to air dry before being wrapped up in plastic film and exposed overnight to a PhosphorScreen. The radioactivity incorporated into each peptide was then determined using a Typhoon FLA 9500 phosphorimager (GE Healthcare) and quantified with the program ImageQuant (version 8.2, Cytiva LifeScience). Data corresponding to the “signal above background” was used.

### Position Weight Matrices (PWMs) generation from OPAL data

PWMs were constructed from the raw OPAL data using a standard approach presented in ^103,104^. First, raw OPAL values for S, T and Y amino acids were replaced with average (median) values for each corresponding peptide position to control for the possibility of spurious phosphorylation in flanking region. The OPAL values were then normalised per position to give a mean PWM score of 1 per amino acid and a total score of 20 per position. The raw OPAL data from S, T and Y libraries was then combined to generate a S/T/Y PWM. This was achieved by summing OPAL scores – after correcting flanking S/T/Y scores – from each of the peptide libraries. The OPAL data was then normalised as before to yield a mean PWM score of 1 per amino acid and a total PWM score of 20 per position. The relative scores between S, T and Y at position was calculated by taking the ratio of the total OPAL scores for the S, T and Y libraries. For ease of visualisation, the PWM logos display only amino acids with the scores above the arbitrary threshold of 2.5 using the software package ggseqlogo^105^. These PWM scores were then log2-transformed to generate heatmaps of the matrix specificity scores.

### FIKK13 peptide RaPID selection

In vitro selections were carried out with Bio-His-FIKK13 following previously described protocols. Briefly, initial DNA libraries (including 6-12 degenerate NNK codons) were transcribed to mRNA using T7 RNA polymerase (37 °C, 16 hr) (Thermo Scientific) and ligated to a puromycin linker primer ([5’Phos]CTCCCGCCCCCCGTCC[SP18][SP18][SP18][SP18][SP18]CC[Puromycin]) using T4 RNA ligase (30 min, 25 °C) (New England Biolabs). First round translation was performed on a 150 µL scale, with subsequent rounds performed on a 5 µL scale. Translations were carried out (30 min, 37 °C then 12 min, 25 °C) using a custom methionine(-) Flexible In vitro Translation system composed by PURExpress^TM^ (ΔRF123) Kit (New England Biolabs) solution B, an in-house solution A (50 mM HEPES-KOH pH 7.6, 2 mM ATP, 2 mM GTP, 1mM CTP, 1mM UTP, 20 mM creatine phosphate, 100 mM potassium acetate, 2 mM spermidine, 6mM magnesium acetate, 1.5 mg/ml E. coli tRNA mix (Roche), 14 mM DTT) and additional ClAc-D-Tyr-tRNA^fMet^ (25 µM). Ribosomes were then dissociated by addition of EDTA (18 mM final concentration, pH 8) and library mRNA reverse transcribed using MMLV RTase, Rnase H Minus. The reaction mixture was buffer exchanged into selection buffer (50 mM Tris Ph 7.5, 50 mM NaCl, 2mM DTT, 10 mM MgCl_2_, 1.5 µM ADP, 0.1% Tween) using 1 mL homemade columns containing pre-equilibrated Sephadex resin (Cytiva) before the addition of 2X blocking buffer (50 mM Tris pH 7.5, 250 mM NaCl, 2 mM DTT, 10 mM MgCl_2_, 1.5 µM ADP, 0.1% Tween, 4 mg/mL sheared salmon sperm DNA (Invitrogen), 0.1% acetyl-BSA final (Invitrogen)). Libraries were incubated with negative selection beads (Dynabeads M-280 streptavidin (Life Technologies)) (3×30 min, 4 °C) followed by incubation with bead-immobilised His-bio-FIKK13 (200 nM, 4 °C, 30 min) before washing (3×1 bead volume selection buffer, 4 °C) and elution of retained mRNA/DNA/peptide hybrids in PCR buffer (95 °C, 5 min). Library recovery was assessed by quantitative real-time PCR relative to a library standard, negative selection, and the input DNA library. Recovered library DNA was used as the input library for the subsequent round. Following 6 rounds of selection, double indexed libraries (Nextera XT indices) were prepared and sequenced on a MiSeq platform (Illumina) using a v3 chip as single 151 cycle reads. Sequences were ranked by total read numbers and converted into their corresponding peptides sequences for subsequent analysis (Supplementary Table 14).

Library DNA:

5’-TAATACGACTCACTATAGGGTTAACTTTAAGAAGGAGATATACATATG (NNK)nTGCGGCAGCGGCAGCGGCAGCTAGGACGGGGGGCGGAAA

Bead preparation:

To assess the binding capacity of biotinylated FIKK13 to streptavidin, Bio-His-FIKK13 was incubated with different quantities of magnetic streptavidin beads (Invitrogen) for 30 min. Beads were then washed three times with cold selection buffer and protein elution was performed by boiling the beads at 95 °C for 5 minutes. Samples were then run in an SDS-PAGE gel and stained with Coomassie. Bead capacity was calculated quantifying the gel bands with FIJI.

### FIKK13 cyclic peptide synthesis

Peptides were synthesised using NovaPEG Rink Amide resin as C-terminal amides by standard Fmoc-based solid phase synthesis as previously described, using a Liberty Blue Peptide Synthesis System (CEM), a SYRO I (Biotage) or a Activotec P-11 peptide synthesiser. Following synthesis, the N-terminal amine was chloroacetylated by reaction with 0.5 M chloromethylcarbonyloxysuccinimide (ClAc-NHS) in DMF (1 hour, RT). Resin was washed (5 x DMF, 5 x DCM) and dried *in vacuo*.

Peptides were cleaved from the resin and globally deprotected with TFA/triisopropyl silane/1,2-ethanedithiol/H_2_O (92.5:2.5:2.5:2.5) for 3 hours at room temperature. Following filtration, the supernatant was concentrated by centrifugal evaporation and precipitated with cold diethyl ether. Crude peptides were resuspended in DMSO/H_2_O (95:5) and, following basification with triethylamine to pH 10, were incubated with rotation for 1 hour at room temperature. Peptides were then acidified with TFA and purified by HPLC (Shimadzu) using a Merck Chromolith column (200 x 25 mm) with a 10-50% gradient of H_2_O/acetonitrile containing 0.1% TFA. Pure peptides were lyophilised and dissolved in DMSO for further use. Peptide stock concentrations were determined by absorbance at 280 nm based on their predicted extinction coefficients.

### Surface plasmon resonance

Single cycle kinetics analysis by SPR was carried out using Biacore S200 and a Biotin CAPture kit, series S (Cytiva). Bio-His-FIKK13 was immobilised on the chip to yield a response of approximately 1400 RU. 50 mM Tris Ph 7.5, 250 mM NaCl, 2mM DTT, 10 mM MgCl2, 1.5 µM ADP, 0.02% Tween and 0.1 % DMSO was used as running buffer and experiments were perfomed at 25 °C. Samples were run with 100 s contact time and data were analysed using the Biacore S200 analysis software. Data represent the average ± standard deviation of at least two independent replicates. SPR data are available in Supplementary Table 8.

### ADP-Glo Assay

Recombinant FIKK kinase domains activity was measured using the ADP-Glo kinase assay (Promega), which quantifies the amount of ADP produced during the kinase reaction. Briefly, the kinase reactions were conducted at room temperature for 1 hour by mixing 100nM recombinant FIKK kinase domain with 10µM ATP and 10µM substrate when specified, in 40µl kinase buffer (20mM MOPS, 10mM magnesium chloride and 10mM manganese chloride, pH 7.4, Alfa Aesar). When kinase inhibition by ATP analogues was assessed, compounds (diluted in DMSO, final concentration δ1%) were tested at 10µM, or otherwise specified, by incubation for 15 minutes with the recombinant kinase domain prior to addition of ATP ± substrate. ADP-Glo reagent (40µl) was added to stop the kinase reaction and deplete the unconsumed ATP. After incubation at room temperature for another hour, 80µl kinase detection reagent was added and incubated for 30 minutes at room temperature. Luminescence was measured using the multi-mode microplate reader FLUOstar Omega (BMG Labtech).

### Protein sequence identity matrix

As described above, *Plasmodium falciparum* FIKK amino acid sequences were retrieved from UniProt and aligned using the MAFFT L-INS-i algorithm^90^.

Heavily gapped alignment positions (more than 20% gapped) were filtered out of the multiple sequence alignment (MSA) using the trimAl software^91^. The sequence identity of this processed alignment was then calculated using seqidentity (,normalise=TRUE) function in the R package bio3d^106^.

### Phosphoproteome libraries analysis

Position weight matrices (PWMs) were calculated for each FIKK kinase as described above. These data were then cross-referenced with the phosphoproteome peptides presented in Fig. 5. Specifically, each peptide in the array was scored for its match to the FIKK preferred phosphorylation motif, using a simple matrix similarity score (MSS) of the PWM against the peptide sequence^101^. This function outputs a normalised score that has a minimum of 0 and a maximum of 1. In each case, a Pearson’s correlation coefficient (PCC) is calculated between the motif score (x) and a log10 transformation of the phosphorylation signal from the phosphoproteome peptide array (y), from curves of the form y = log(x).

Peptides from the library were divided into a motif ‘match’ and ‘no match’ with respect to any given FIKK specificity matrix. This was based on a null distribution of randomised peptide sequences for phosphosites not affected by the FIKK knockout tested previously^16^. Peptides with a motif score (MSS) below an empirical *p-value* of 0.05 were considered a ‘Match’. Peptides with a MSS above 0.05 in *p-value* were considered ‘No Match’.

The sequence logo of phosphoproteome targets (e.g. in Fig. 5c) represents the relative frequency of amino acids among peptides phosphorylated above background levels (log10(signal >4.0) for the FIKK kinase of interest. Sequence logos were generated using ggseqlogo^105^.

### Expression and purification of *P. falciparum* FIKK13 kinase domain proteins for crystallisation

Codon-optimised DNA encoding the kinase domain of *Pf*FIKK13 residues 149-561 (PlasmoDB PF3D7_1371700) was cloned into pET-47b to produce an HRV 3C cleavable His_6_ N-terminal fusion (MA**HHHHHH**SAA<u>LEVLFQ⇩GP</u>G) with the HRV 3C cleavage site underlined. The kinase inactive D379N mutant was generated by site directed mutagenesis using the pET-47-*Pf*FIKK13^149–561^ construct as a template. Both constructs were verified by DNA sequencing.

The *Pf*FIKK13^149–561^ and *Pf*FIKK13^149–561_D379N^ were expressed in *E. coli* strain BL21 (DE3) Gold (Agilent). Bacterial cultures were grown in TB at 30°C to an OD_600_=1.2-1.5 and isopropyl-β-D-thiogalactoside (IPTG) was added to a final concentration of 0.5mM to the culture grown at 25°C overnight. Cell pellets were harvested, resuspended in lysis buffer A (50mM HEPES pH7.5, 20mM Imidazole, 0.5M NaCl, 10mM MgCl_2_, 10%(v/v) Glycerol, 1mM TCEP) supplemented with 1U/ml Universal nuclease (Pierce) and 1 Protease inhibitor tablet (cOmplete, Roche) per 50ml solution, and lysed by sonication. The bacterial lysate was centrifuged for 30 minutes at 80,000xg. The supernatant was applied to a 5ml HisTrap column (Cytiva) and washed with 10 CV of buffer A. Fractions containing *Pf*FIKK13^149–561^ and *Pf*FIKK13^149–561_D379N^ were pooled separately and incubated overnight at 4°C with HRV 3C protease to remove the 6xHis-tag. The next day, *Pf*FIKK13^149–561^ and *Pf*FIKK13^149–561_D379N^ were concentrated and purified by size-exclusion chromatography using buffer B (50mM HEPES pH7.5, 250mM NaCl, 2mM MgCl_2_, 1mM TCEP and 5%(v/v) glycerol) as running buffer.

### Generation of nanobodies recognising *Pf*FIKK13^149–561^

A healthy llama (Arla) was immunised with 3 doses of 200µg of purified *Pf*FIKK13^149–561^ using GERBU as adjuvant followed an established protocol with animal handling carried out by trained personnel under the Home Office Project Licence PA1FB163A. The three immunisations took place on Day 0, 28 and 56 and a 150ml blood sample was harvested 10 days after the 3^rd^ immunisation. The PBMCs were isolated and total RNA extracted using described methods^107^. Total RNA was reverse transcribed using dT18-oligos. The VHH was PCR-amplified using primers CALL01 and CALL02 and a band of ∼700bp excised. The 700bp band was used as template to re-amplify the VHH using primers VHH-Sfil2 (5’-GTCCTCGCAACTGCGGCCCAGCCGGCCATGGCTCAGGTGCAGCTGGTGGA-3’) and VHH-Not2 (5’-GGACTAGTGCGGCCGCTGAGGAGACGGTGACCTGGGT-3).

The PCR product was digested with *SfiI* and *NotI* enzymes (NEB) and ligated into a pHEN2 vector modified with a triple c-Myc tag^108^ which was used to transform electrocompetent TG1 cells (Lucigen). The resulting Nb-library consisted of 5×10^7^ independent colonies. Phage particles were prepared by super-infection with VCS13 helper phage (Agilent). The phage preparation was concentrated by adding 1/5^th^ volume of 20%(W/v) PEG6000, 2.5M NaCl and subsequent centrifugation at 4000xg for 30min.

Nanobodies specific for *Pf*FIKK13^149–561^ were selected against biotinylated *Pf*FIKK13^149–561^ immobilised on either Streptavidin-coated magnetic beads (Dynabeads™ M280-Streptavidin, ThermoFisher Scientific) or Pierce™ NeutrAvidin™ Coated plates (#15123 ThermoFisher Scientific). Individual clones were isolated by ELISA using soluble Nbs as primary antibody and detecting with the anti-C-myc antibody 9E10, followed by anti-mouse-HRP conjugated antibodies (Agilent). The chromogenic TMB substrate (ThermoFisher Scientific) was added, and colour development was quenched with 1M HCl. The absorbance was read at 450nm with 620nm used as baseline. ELISA-positive Nb clones were sequenced. A total of 7 different families of Nbs was isolated. Further work was performed with Nb2G9 and Nb9F10.

### Expression and purification of Nb2G9 and Nb9F10

Nb2G9 (protein sequence: QVQLVESGGGSVQAGGSLRLSCAASGRTFSSYSMAWFRQAPGKERENVAVISWSGSTSYYAESVKGRFTISRDNAKNTVYLQMNSLKPEDTAVYYCAAGPRTTPQAMGAVEYDYWGQGTQVTVSS) was found to be compatible with Nb9F10 (protein sequence: QEQLVESGGGLVQAGGSLTLSGASSGGTFETYAMGWFRQAPGKEREFAAAVSWSGGSAHYADSVKGRFTISRDKVKNTVYLQMNSLKPEDTAVYYCAADRSYGSSWYHYPEDALDAWGQGTQVTVSS) in simultaneously binding *Pf*FIKK13^149–561^ (data not shown). The DNA encoding Nb2G9 and Nb9F10 were cloned into a modified pET-21b vector to produce a N-term pelB secretion signal fusion (MKYLLPTAAAGLLLLAAQPA⇩MA) with a C-term TEV cleavable Avi•Tag/His_8_ (AA<u>ENLYFQ⇩G</u>*LNDIFEAQKIEWHE***HHHHHHHH** where the underlined sequence is the TEV cleavage site, the Avi•Tag in italics and the 8xHis-tag in blod).

Nb2G9 and Nb9F10 were expressed in *E. coli* strain Rosetta2(DE3). Bacterial cultures were grown in TB at 37°C to a density of OD_600_=1.2-1.5 and protein expression was induced with 0.5mM IPTG at 30°C overnight. The bacteria were pelleted by centrifugation at 5,000xg for 30min. The clarified TB medium containing nanobodies was adjusted to pH8.0, 20mM NaCl and loaded onto a 5ml HiTrap Excel column (Cytiva) pre-equilibrated in PBS. Bound proteins were eluted with PBS containing 400mM imidazole pH8.0. The C-term Avi/His_8_ tag was removed by TEV protease cleavage and subsequent size-exclusion chromatography.

### FIKK13/Nb2G9/Nb9F10 complex formation

Purified *Pf*FIKK13^149–561_D379N^ and Nb2G9 and 9F10 were mixed in a 1:1.2:1.2 molar ratio and loaded on a Superdex 75 10/300 Increase column (Cytiva) equilibrated in 20mM HEPES pH7.5, 150mM NaCl, 2mM MgCl_2_, 0.5mM TCEP to remove the excess nanobodies. The fractions corresponding to the FIKK13^149–561_D379N^/Nb2G9/Nb9F10 complex peak were pooled, concentrated to 7mg/ml and used for crystallisation experiments. Additionally, the FIKK13^149–561_D379N^/Nb2G9/Nb9F10/ATPγS samples were prepared by mixing purified FIKK13^149–561_D379N^/Nb2G9/Nb9F10 and ATPγS in a 1:1.1 molar ratio.

### Crystallisation of *Pf*FIKK13^149–561_D379N^ with Nb2G9, Nb9F10 and ATPγS

Crystallisation trials were set up using samples at ∼7mg/ml. Initial crystals of the complex between *Pf*FIKK13^149–561_D379N^ and Nb2G9 and Nb9F10 nanobodies were grown in 20%(w/v) PEG3350 and 0.2M Sodium thiocyanate (Peg Ion HT screen condition B1, Hampton Research) and further optimised. Crystals were grown in sitting drops by vapor diffusion at 20°C, cryoprotected by stepwise addition of PEG4000 or ethylene glycol to a final concentration of 25% (v/v), and flash-cooled to 100K by direct immersion in liquid nitrogen.

The initial apo (without ATPγS) structure was solved at low resolution by molecular replacement using an AlphaFold search ensemble generated at Diamond Light Source using data obtained from crystals grown in 18%(w/v) PEG3350, 150mM Sodium thiocyanate and 10mM Calcium chloride as an additive. The model was rebuilt and refined before molecular replacement into a higher resolution dataset obtained from crystals grown in 21%(w/v) PEG3350 and 0.1M Sodium thiocyanate.

Crystals of *Pf*FIKK13^149–561_D379N^ in complex with Nb2G9 and Nb9F10 bound to ATPγS were obtained from a condition containing 0.1M lithium chloride, 10%(v/v) Ethylene glycol, 20%(w/v) PEG6000 and 0.1M HEPES pH7.0 (Ligand Friendly Screen condition C9, Molecular Dimensions) and seeding with apo crystals.

### Kinase-peptide models generation

Kinase-peptide models were generated using the HADDOCK 2.4 webserver^109^ applied to AlphaFold2 predictions of the FIKK kinase domain^110,111^. Docking was executed using default parameters with the following alterations. First, residues with a minimum relative solvent accessibility (RSA) of 5% could be considered as accessible. Second, the peptide sequence was designated as the molecule type ‘Peptide’ and defined to be fully flexible at every position. No ‘active’ or ‘passive’ residues were chosen as ambiguous interactions restraints (AIRs)^112^, but an unambiguous interaction restraint was specified between the phosphoacceptor (S, T, or Y) side chain oxygen and the hydroxyl oxygen of the catalytic aspartate residue (D166 in PDB: 1ATP), at a maximum distance of 5.5 Angstroms.

### PKIS screen and Structure Activity Relationship (SAR) assays

PKIS and SAR compounds at 1mM in 60nl DMSO were plated in white, opaque, flat-bottomed, 384-well microplates (Greiner Bio-one). Columns 6 and 18 of the microplates served as controls. Column 6 = positive control (Recombinant FIKK8 kinase domain + *P_o_* peptide (RRRAPSFYRK)^22^ + ATP without compounds) Column 18 = negative control (Recombinant FIKK8 kinase domain + *P_o_* peptide – ATP). 3µl kinase at 40nM in kinase reaction buffer (20mM MOPS, 10mM magnesium chloride and 10mM manganese chloride, pH 7.4, Alfa Aesar) was dispensed in each well using a Multidrop™ Combi Reagent dispenser (ThermoFischer Scientific). Recombinant FIKK8 kinase domain was left to incubate in the presence of compounds for 15 minutes at room temperature. 3µl *P_o_* peptide + ATP at 20µM each, diluted in kinase reaction buffer, was dispensed in each well except in column 18 in which 3µl of peptide without ATP was dispensed. Kinase reaction was left to occur for 1 hour and was stopped with 6µl ADP-Glo reagent. Kinase activity was assessed after addition of 12µl Kinase Detection reagent by measuring luminescence on a multi-mode microplate reader FLUOstar Omega (BMG Labtech).

### *In vitro* measurement of compounds IC_50_

*In vitro* half-maximal inhibitor concentrations of PKIS and SAR compounds was determined by testing recombinant FIKK8 kinase domain activity in the presence of increasing concentrations of compounds. White, opaque, flat-bottomed 384-well microplates (Greiner Bio-One) containing compounds in a range of concentrations starting from 25µM to 0.4nM (1 in 3 serial dilutions) were ordered from GSK in Stevenage. Compounds were dispensed in the microplates at the required concentrations in 60nl DMSO. Kinase activity in the presence of the compounds was measured as described above. Briefly, 3µl recombinant FIKK8 kinase domain at 40nM in kinase reaction buffer was dispensed in each well using a Multidrop™ Combi Reagent dispenser (ThermoFischer Scientific). Recombinant FIKK8 kinase domain was left to incubate in the presence of compounds for 15 minutes at room temperature. 3µl *P_o_* peptide + ATP at 20µM each, diluted in kinase reaction buffer, was dispensed in each well. Kinase reaction was left to occur for 1 hour and was stopped with 6µl ADP-Glo reagent. Kinase activity was assessed after addition of 12µl Kinase Detection reagent by measuring luminescence on a multi-mode microplate reader FLUOstar Omega (BMG Labtech). Data were analysed using GraphPad Prism version 10 and IC_50_s were calculated from a four-parameters logistical fit of the data.

### FIKK inhibitors EC_50_s determination

Half maximal effective concentration (EC_50_) of the different compounds tested was determined by flow cytometry. Two-fold dilutions of the compounds were plated in triplicate in 96 well-plates. 200µl parasite solution containing 1% NF54 parasitemia and 2% haematocrit was added to each well and plates were incubated for 72 hours at 37°C in a sealed gassed chamber. After incubation, parasite growth was assessed by flow cytometry. Samples (20µl) were fixed in 2% paraformaldehyde (PFA) + 0.2% glutaraldehyde (GA) in PBS for 1 hour in the dark at 4°C. Fixative was subsequently washed out with PBS and samples were stained with SYBR Green for 30 minutes in the dark at 37°C. After a final wash, parasitemia was counted by flow cytometry on a BD LSR Fortessa flow cytometer (Becton Dickinson) using the FACS Diva software. Data were analysed using the FlowJo 10 analysis software (Becton Dickinson).

### ATP-depletion

Irreversible depletion of ATP in RBCs was carried out by incubating uRBCs for 2 hours at room temperature in PBS containing various concentrations of inosine and iodoacetamide^70^ (see Extended Data Fig. 21 for inosine and iodoacetamide concentrations used). ATP-depleted uRBCs were then washed three times with PBS and ATP-depletion was evaluated for each dilution using the CellTiter-Glo→ Luminescent Cell Viability assay (Promega) following the instructions provided in the kit (see Extended Data Fig. 21 for ATP-depletion assessment). ATP-depleted uRBCs were put in the presence of Percoll-purified mature schizont stage parasites at 1% haematocrit in complete medium in a shaking incubator at 37°C for 4 hours. Parasites were allowed to grow for 48 hours before samples were taken for immunofluorescence, Western blot and flow cytometry assessment of parasite growth as described above.

## Data availability

The mass spectrometry proteomics data have been deposited to the ProteomeXchange Consortium via the PRIDE^113^ partner repository with the dataset identifier PXD048966. The crystal structure of *Pf*FIKK13^149–561_D379N^ with Nb2G9, Nb9F10 and ATPγS is available from the Protein Data Bank under the accession code …. Gene sequences and annotations for *P. falciparum* 3D7 were acquired from PlasmoDB.org (v46)^13^ and human sequences were acquired from Uniprot.org (2023)^89^. RNA sequencing data from Hoeijmakers *et al.* available on PlasmoDB was also used. The Pf3K project dataset used to identify genetic variants in fikk genes is available at the following address (www.malariagen.net/projects/parasite.pf3k)^79^. Source data in the form of unprocessed gels and western blots corresponding to Figs. 1c, 7d, 7e, 7f and Extended Data Figs. 2a, 2b, 4, 21c are available with the article.

## Code availability

No custom code deemed central to the conclusions to this manuscript has been used in this study.

## Supporting information

Supplementary Table 1_FIKK orthologues in Laverania

Supplementary Table 2. FIKK transcript levels during the P.falciparum asexual replication cycle

Supplementary Table 3. STOP codon in fikk genes in 2085 field isolate genomes

Supplementary Table 4. fikk genes deletions in 2085 field isolate genomes

Supplementary Table 5. Transcription evidence for PfFIKK kinases in Gametocytes and mosquito stages according to Malaria Cell Atlas data

Supplementary Table 6. Table of proteins in the vicinity of FIKK4.1 and/or FIKK4.2 identified by TurboID-based proximity labelling

Supplementary Table 7. Start sites of recombinantly expressed FIKK kinase domains

Supplementary Table 8. SPR binding affinities for FIKK13 peptides

Supplementary Table 9. Phosphoproteome peptide library composition

Supplementary Table 10. Half maximal inhibitory concentration (IC50) of the PKIS compounds identified as inhibitors of FIKK8 recombinant kinase domain

Supplementary Table 11. Material used for the generation of FIKK_TurboID parasite lines

Supplementary Table 12. Recodonised sequences used for FIKK kinase domains expression in E. coli

Supplementary Table 13. Processed mass spectrometry proximity labelling data

Supplementary Table 14. FIKK13 RaPID selection Next Generation Sequencing Data

Supplementary Table 15. Flow cytometry data for EC50s determination of FIKK inhibitors in P. falciparum and P. knowlesi

Supplementary Table 16. Flow cytometry data for ATP-depletion optimisation experiment

## Acknowledgments

We thank members of the Treeck, Blackman, Knuepfer and Sateriale labs for critical discussions. We also thank the Crick Science Technology Platforms (STPs) (Proteomic and Flow cytometry) for their outstanding technical support and training. We thank Dr. Julian Rayner and L. Parish for the MAHRP1 and GAP50 antibodies, Dr. Tobias Spielmann for the SBP1 antibody and the European Malaria Reagent Depository for the FIKK4.2 and GAPDH antibodies. We thank Dr. Ellen Yeh for sharing insights on the proximity labelling experiments in malaria infected RBCs. We thank GSK for its commitment to support fundamental discovery research through the establishment of the Crick-GSK LinkLabs partnership. We would also like to thank Hong Lin, Barney Jones and Professor Gary Stephens, University of Reading, for expert help with generation of nanobodies under the authority of PA1FB163A. Special thanks to PlasmoDB and VEuPathDB for providing critical resources. M.T. received funding from the ERC (ERC Grant number: 101044428) and the Francis Crick Institute (Grant No. CC2132). M.T.B and L.J.W. were supported by the Francis Crick Institute (CC2030). The Francis Crick Institute and its Science Technology platforms receive core funding from Cancer Research UK, the UK Medical Research Council and the Wellcome Trust (Grant No. CC0199). S.D.N. is funded by an Early-Career Award Wellcome Trust grant (225686/Z/22/Z). C.R.L. and D.B. were supported by a Canadian Institutes of Health Research Foundation grant number 387697 and a Human Frontier Science Program research grant RGP34/2018 to C.R.L. C.R.L. holds the Canada Research Chair in Cellular Systems and Synthetic Biology. D.B. was supported by an EMBO Long-Term Fellowship (LTF) (ALTF 1069-2019).

## Authors contributions

H.B. performed the parasite genetic manipulations and phenotypic analysis, the kinase activity assays on peptides and membranes. D.B. performed the bioinformatics analysis with input from C.R.L.. S.D.N. and H.B. performed the gametocyte experiments. H.B., H.D. and M.B. processed the proteomic samples. H.B., D.B. and M.B. analysed the proteomic data. H.B., E.C. and D.J. expressed and purified the recombinant proteins. H.B. and D.B. analysed the substrate specificity data. M.T.B. performed the cyclic peptide screen and SPR under supervision from L.W. D.Joshi generated the peptide libraries and synthetic peptides under supervision of N.O’R.. E.C., A.G.P., R.W.O. and S.K. generated protein crystals and solved the protein crystal structure. D.B designed the FIKK mutants. A.C. performed the field isolates genome analysis. H.B. performed the inhibitor screens under the supervision of A.P. and D.H. H.B. and M.T. conceived the study. H.B., D.B., S.D.N., H.D., M.T.B., S.K. and M.T. designed figures. H.B., D.B., S.D.N., M.T.B., A.C., S.K., C.R.L and M.T. wrote the original manuscript. All authors were involved in critically reviewing and editing the manuscript.

---

**Extended Data Fig. 1.**
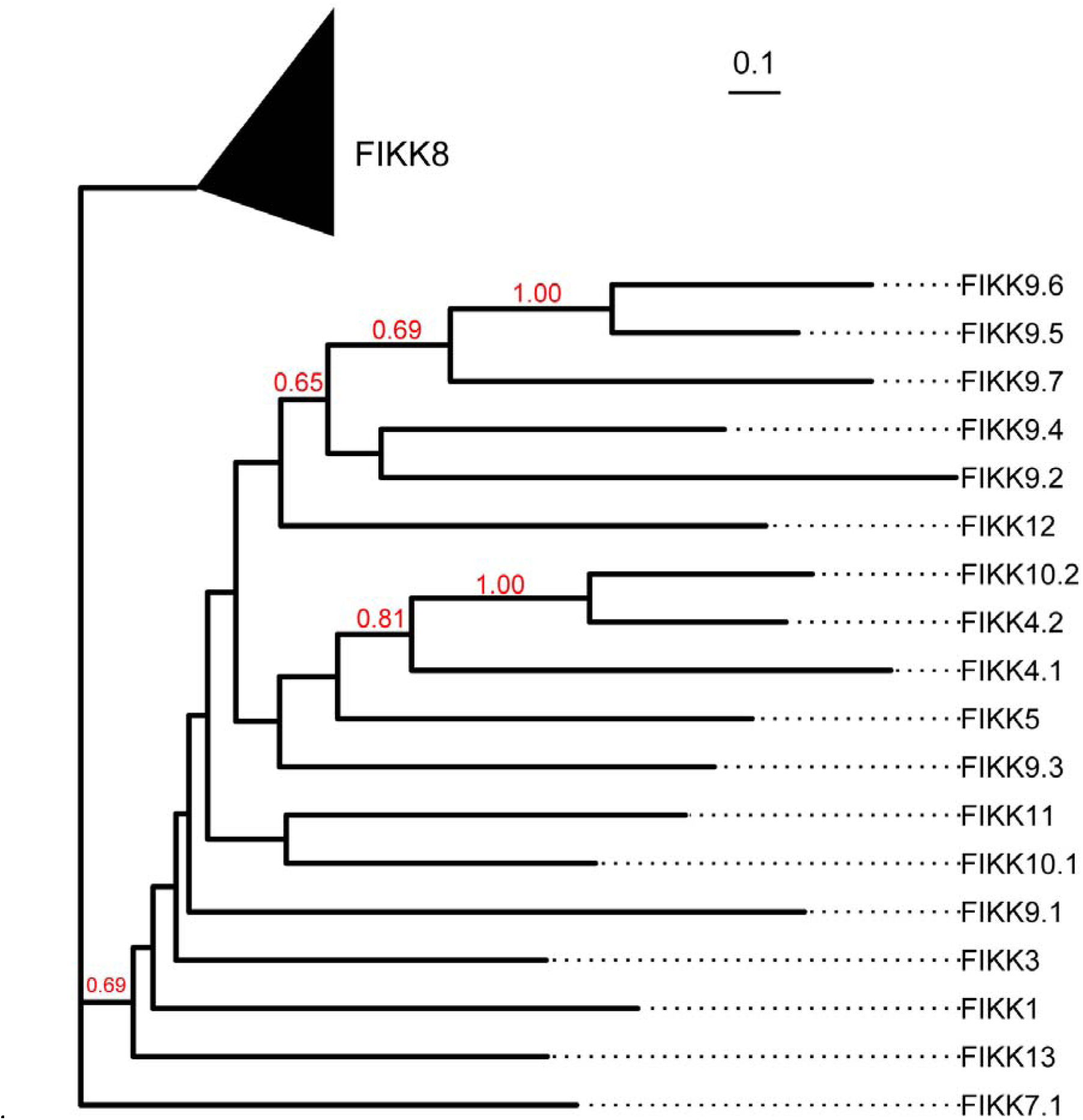
Phylogenetic tree of *Pf*FIKK kinases rooted on FIKK8 sequences. Maximum-likelihood phylogenetic tree of *P. falciparum* FIKK kinase sequences (see Methods). The tree was rooted using known FIKK8 sequences across *Plasmodium* species. Branch support was assessed using 100 bootstrap replicates^50^ and is shown for branches with support > 0.5.

**Extended Data Fig. 2.**
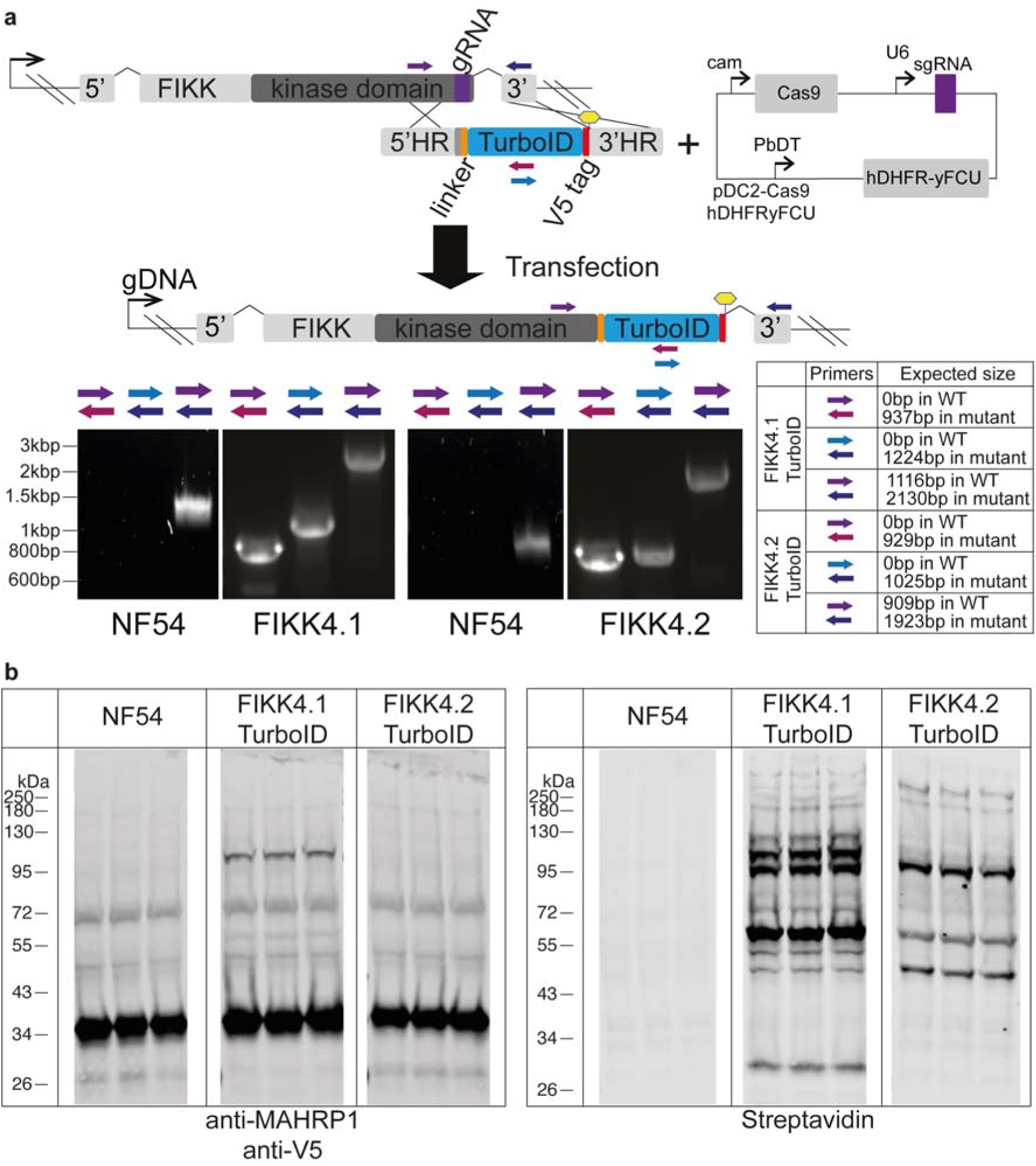
CRISPR/Cas9 strategy to generate FIKK::TurboID fusion proteins and validation. **a**, Diagram illustrating the CRISPR/Cas9 strategy^33^ used to insert a TurboID_V5 cassette at the C-terminal end of the *fikk* genes. Homology regions used to edit the genome are denoted by 5’ and 3’HR and the Cas9 guide is denoted as a purple cassette. Yellow hexagons denote a stop codon. Primers used to investigate integration into the correct endogenous loci along with the presence of WT parasites into the mutant population are shown. Expected band size for PCR reactions are indicated. Material used to generate and validate FIKK::TurboID lines can be found in Supplementary Table 11. **b,** Western blots of cloned parental NF54, FIKK4.1::TurboID (111kDa) and FIKK4.2::TurboID (180kDa) fusion lines cultured in biotin-containing medium for the duration of the parasite asexual lifecycle (48h) probed with anti-V5 and streptavidin-fluorophore. Anti-MAHRP1 (29kDa) antibody is used as a loading control.

**Extended Data Fig. 3.**
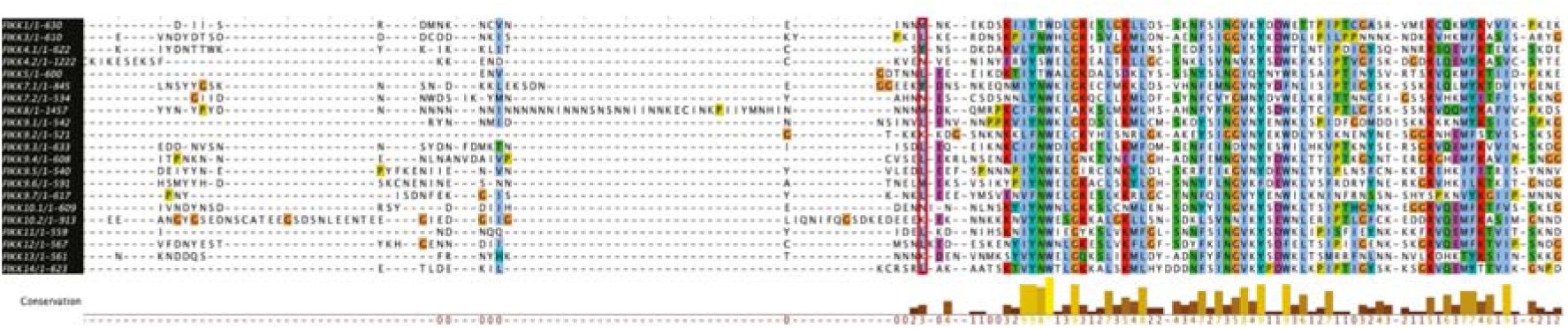
Alignment of *P. falciparum* FIKK protein sequences allows for accurate determination of the FIKK kinase domain starting amino acid. Alignment of all FIKK kinase sequences from *P. falciparum* using the T-Coffee multiple sequence alignment program^47^ available in the Jalview software^48^. Encircled in red are the amino acids chosen as a starting point for recombinant expression of *P. falciparum* FIKK kinase domains. The ClustalX colour scheme was used to assign colour to amino acids with the following criteria: Blue – Hydrophobic (A, I, L, M, F, W, V, C); Red – Positively charged (K, R); Magenta – Negatively charged (D, E); Green – Polar (N, Q, S, T); Orange – Glycine (G); Yellow – Proline (P); Cyan – Aromatic (H, Y); White – unconserved amino acids. Below the alignment is indicated the conservation score which measures the number of physicochemical properties conserved for each column of the alignment. Its calculation is based on ^49^. Conserved columns are indicated by * (score of 11), conservation score then ranges between 10 (high conservation) and 0 (no conservation). Hyphens denote gaps.

**Extended Data Fig. 4.**
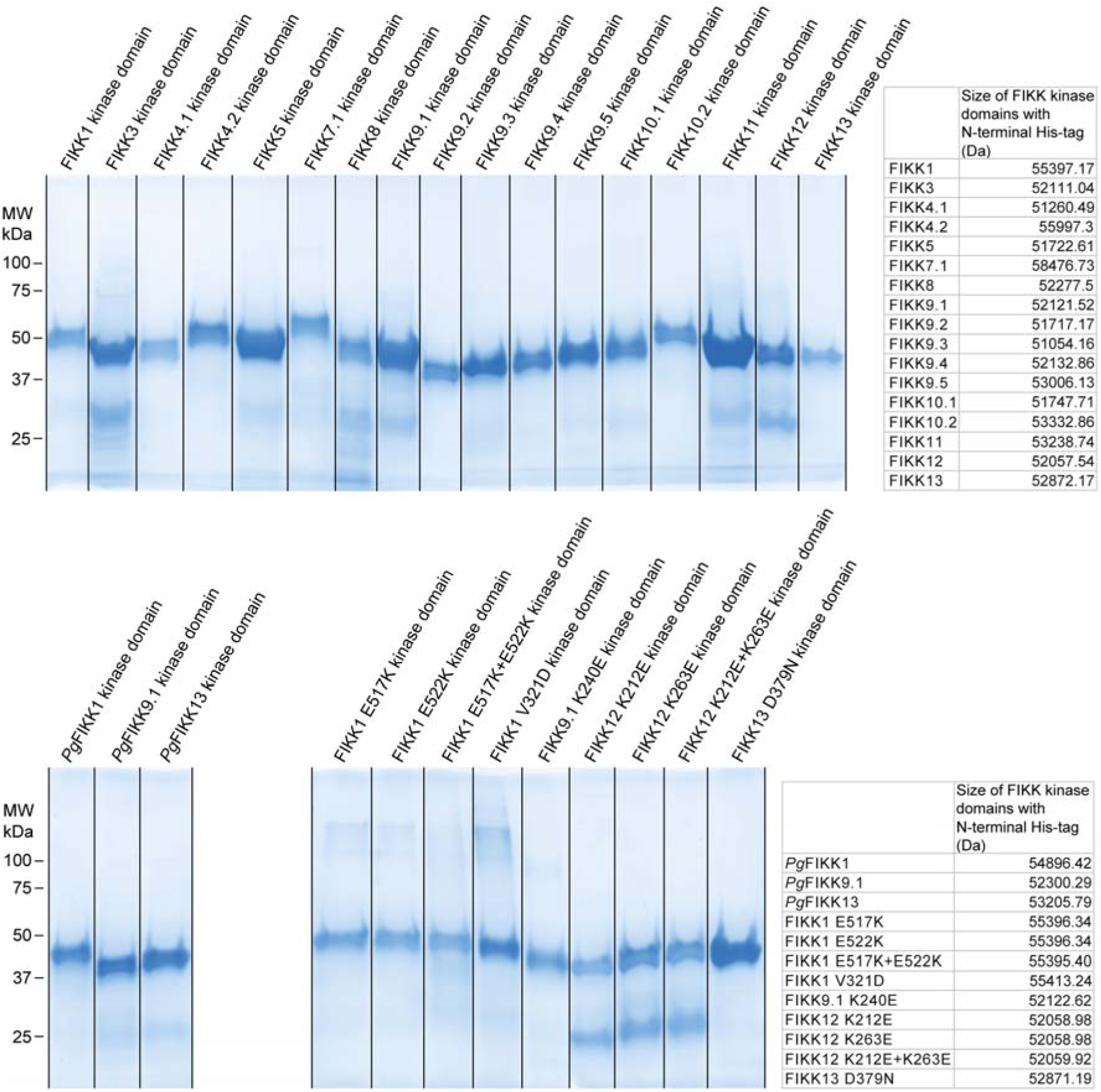
Coomassie-stained gel of purified recombinant FIKK kinase domains. Protein ladder is depicted on the left-hand side of the gel in kilodaltons (kDa). Predicted sizes of the purified recombinant kinase domains with N-terminal His-tag are indicated in Dalton in the table.

**Extended Data Fig. 5.**
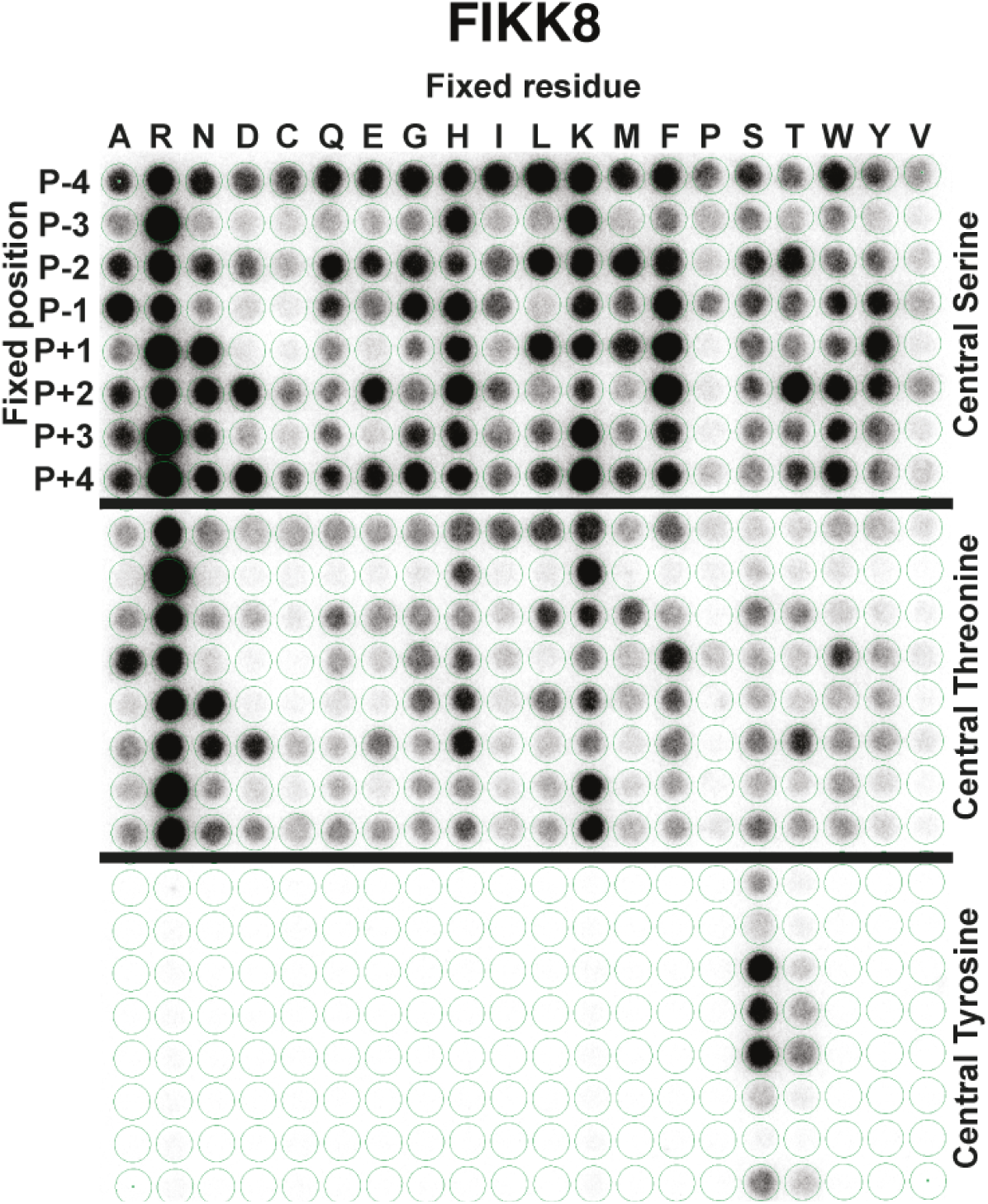
FIKK8 OPAL membrane. An OPAL membrane constituted of 9-mer peptides with the general sequences A-X-X-X-X-S-X-X-X-X-A (top panel), A-X-X-X-X-T-X-X-X-X-A (middle panel) or A-X-X-X-X-Y-X-X-X-X-A (bottom panel) was used to assess FIKK8 preferred phosphorylation motif. X represents any natural amino acid except for S, T, Y or C. For each peptide, one the 20 naturally occurring amino acids is fixed at each one of the 8 positions surrounding the phosphorylatable residue (S, T or Y). The membrane was incubated in the presence of recombinant FIKK8 kinase domain and [LJ-32P]-ATP. After several washes, the membrane was exposed overnight to a phosphorscreen. The radioactivity incorporated into each peptide was determined by scanning the phosphorscreen with a phosphorimager giving the radiograph visible in this figure. Plotting the intensity pattern of the array enables the identification of preferred phosphorylation motifs (Fig. 3bii) and reveals amino acids that are less favoured in a peptide sequence.

**Extended Data Fig. 6.**
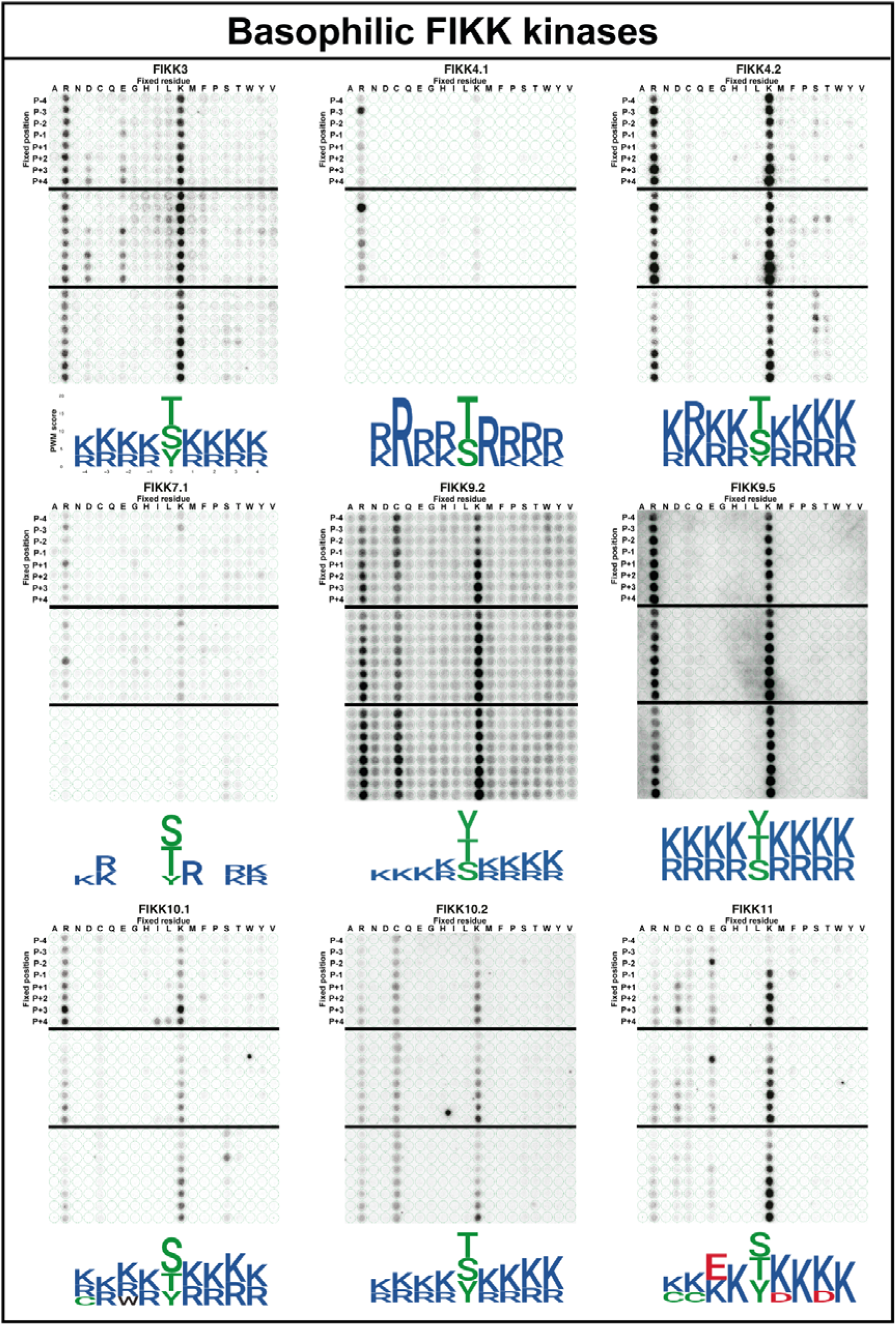
Basophilic FIKK kinases preferred phosphorylation motifs. See Extended Data Fig. 5 caption.

**Extended Data Fig. 7.**
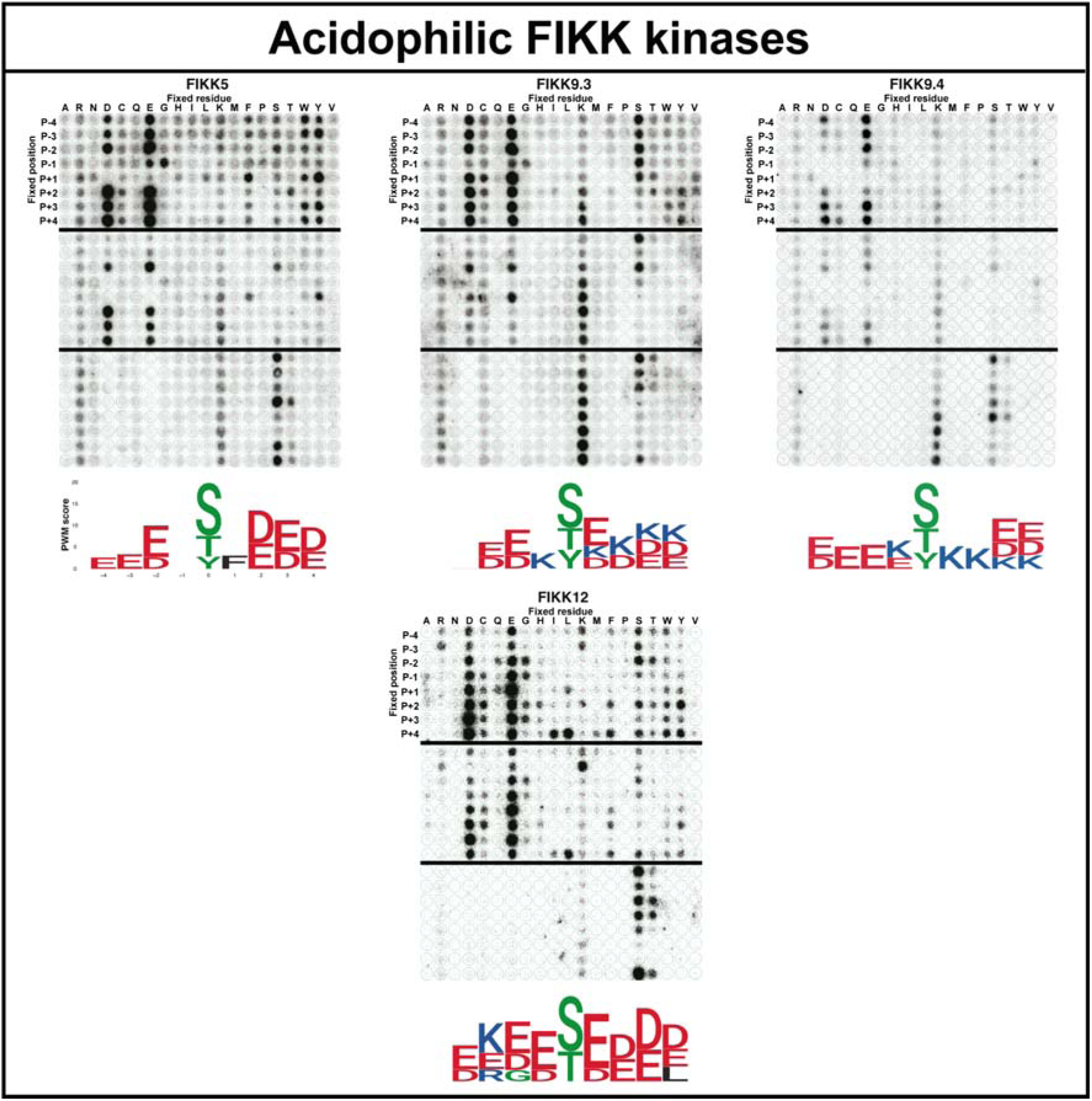
Acidophilic FIKK kinases preferred phosphorylation motifs. See Extended Data Fig. 5 caption.

**Extended Data Fig. 8.**
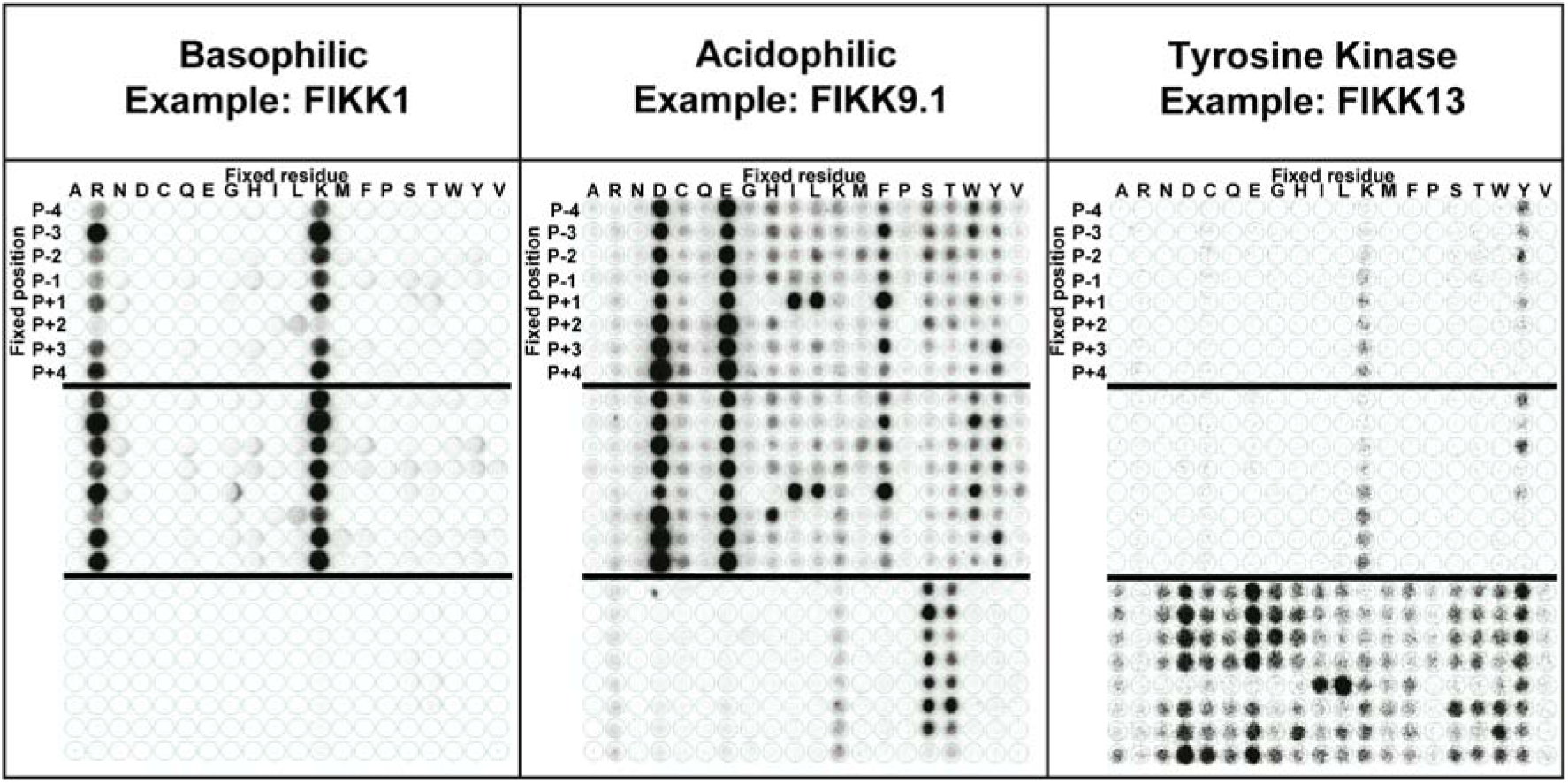
FIKK1, FIKK9.1 and FIKK13 OPAL membranes. See Extended Data Fig. 5 caption.

**Extended Data Fig. 9.**
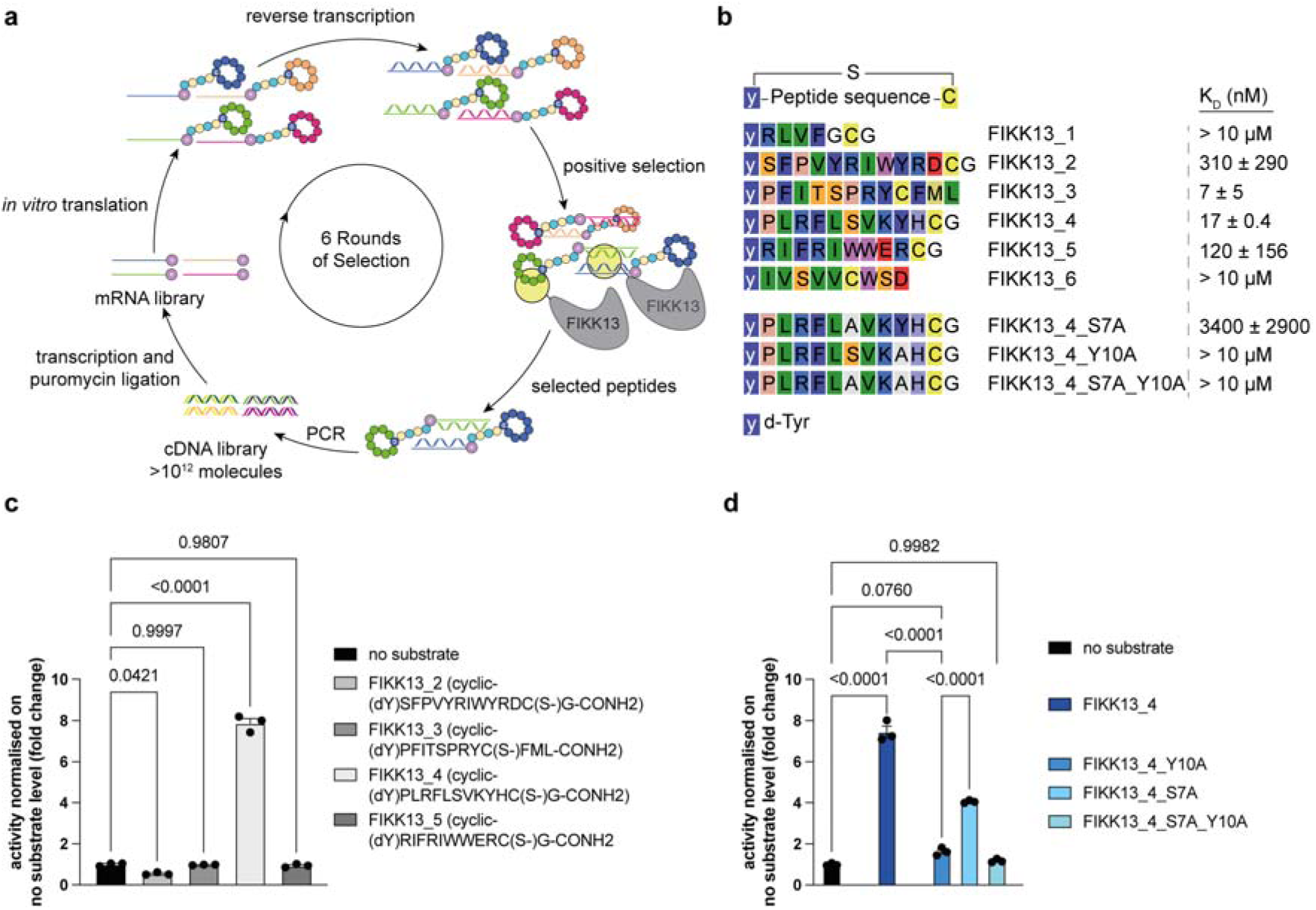
Identification of a tyrosine-based cyclic peptide as a substrate for FIKK13. **a**, Scheme of the FIKK13 RaPID selection. **b,** Sequences and binding affinities of the different peptides recovered after 6 rounds of selection and different variants of the parent peptides. Peptides were initialised with d-Tyr and cyclised via a thioether bond between the N-terminus and the cysteine side chain. Binding affinities were measured by SPR (Supplementary Table 8) and show average ± standard deviation of at least 2 independent replicates. **c,** FIKK13 kinase domain phosphorylating activity on cyclic peptide identified in panel **b**. The results are represented as the mean±SEM fold change compared to the no substrate luminescent signal obtained using the ADP-Glo assay. Statistical significance was determined using a one-way ANOVA followed by Dunnett’s multiple comparison post-test. n=3 biological independent replicates. **d,** FIKK13 kinase domain phosphorylating activity on FIKK13_4 mutant peptides. The results are represented as the mean±SEM fold change compared to the no substrate luminescent signal obtained using the ADP-Glo assay. Statistical significance was determined using a one-way ANOVA followed by Šidák’s’s multiple comparison post-test. n=3 biological independent replicates.

**Extended Data Fig. 10.**
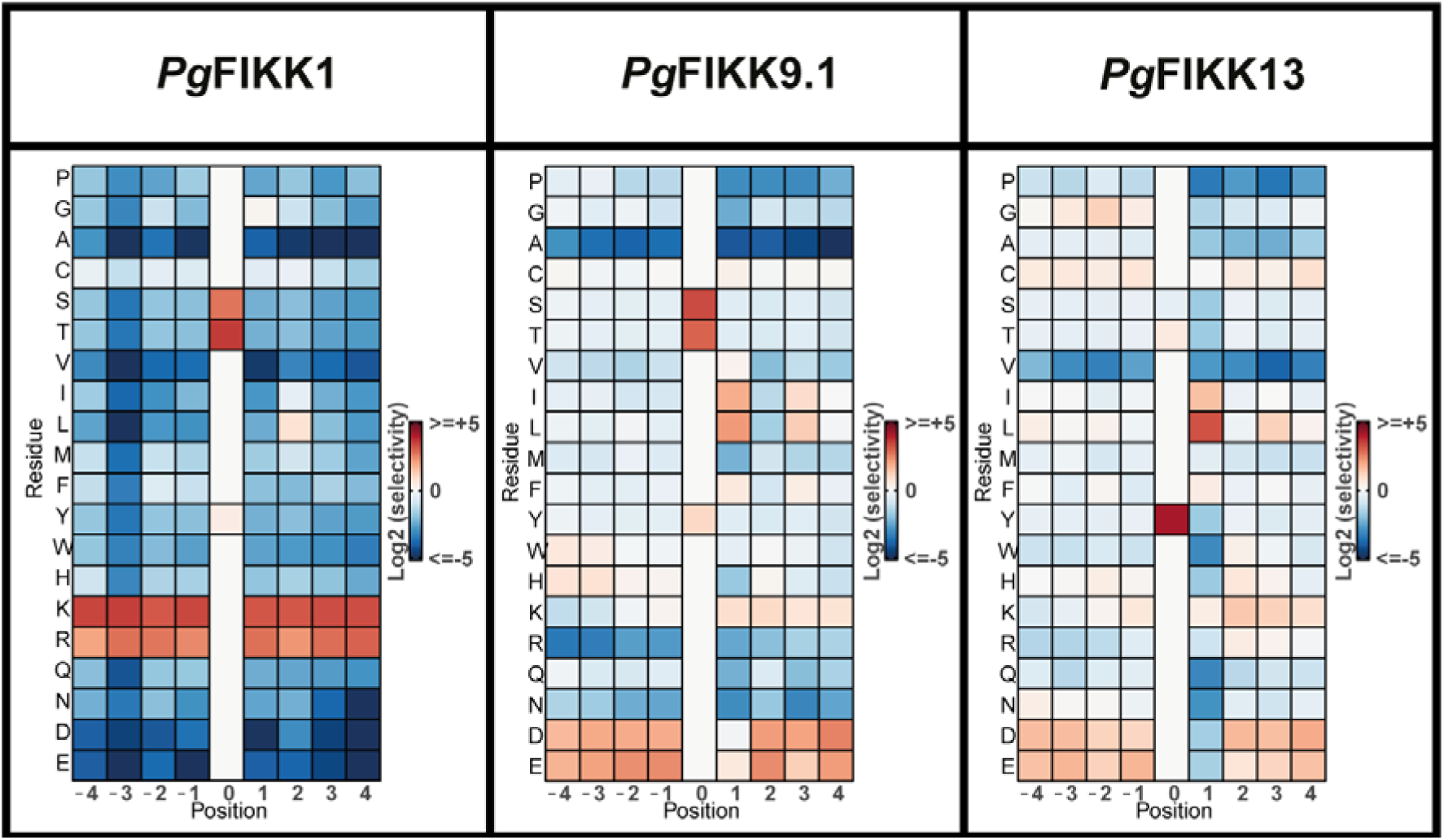
Heat map representation of OPAL arrays raw data for *P. gaboni* FIKK1, FIKK9.1 and FIKK13. Heat map representation of OPAL array raw data for *P. gaboni* FIKK1, FIKK9.1 and FIKK13. See Fig. 3 caption.

**Extended Data Fig. 11.**
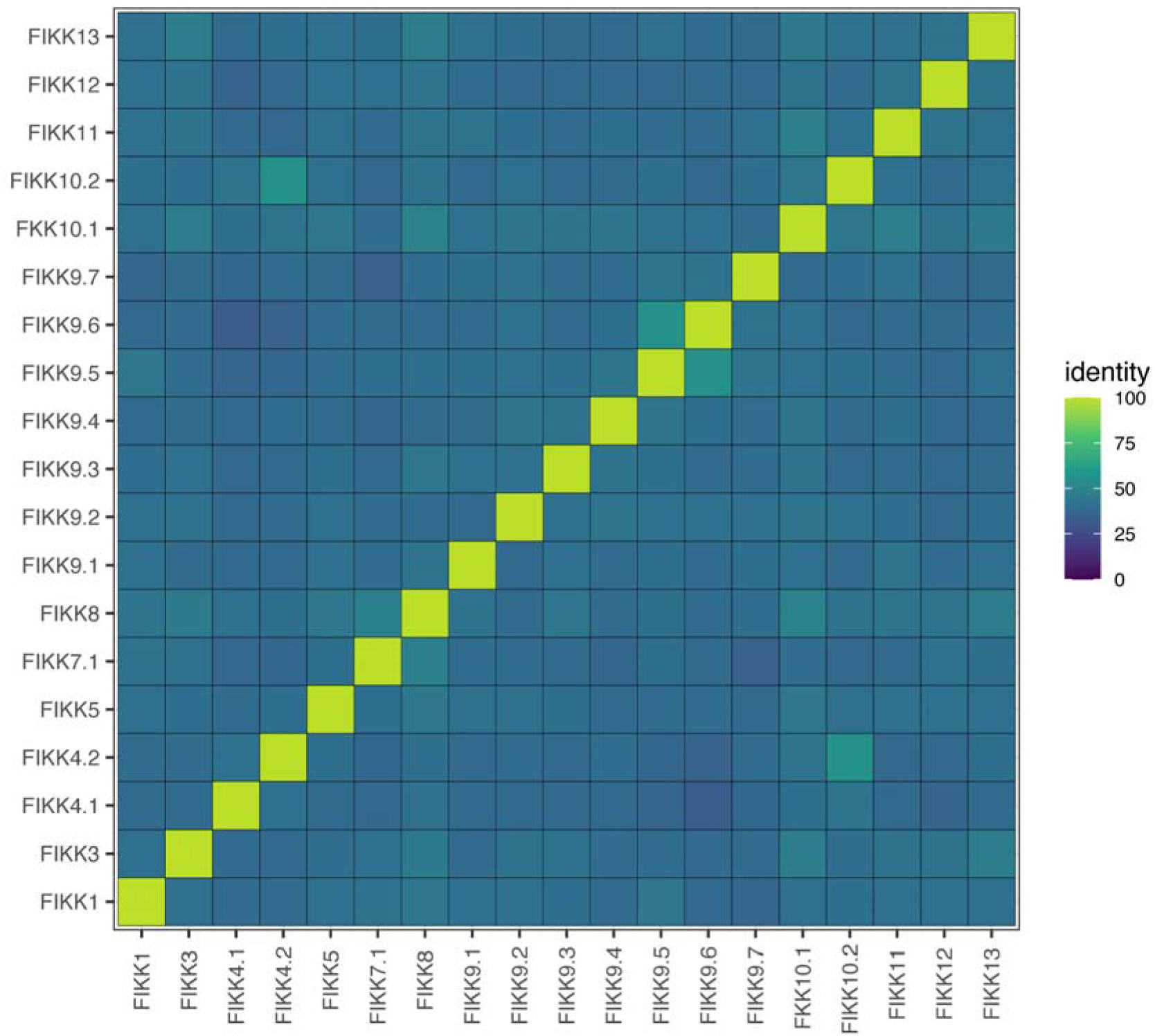
Protein sequence identity matrix of *P. falciparum* FIKK kinases. Amino acid sequence identity on the basis of the *P. falciparum* multiple sequence alignment after removing poorly aligned regions.

**Extended Data Fig. 12.**
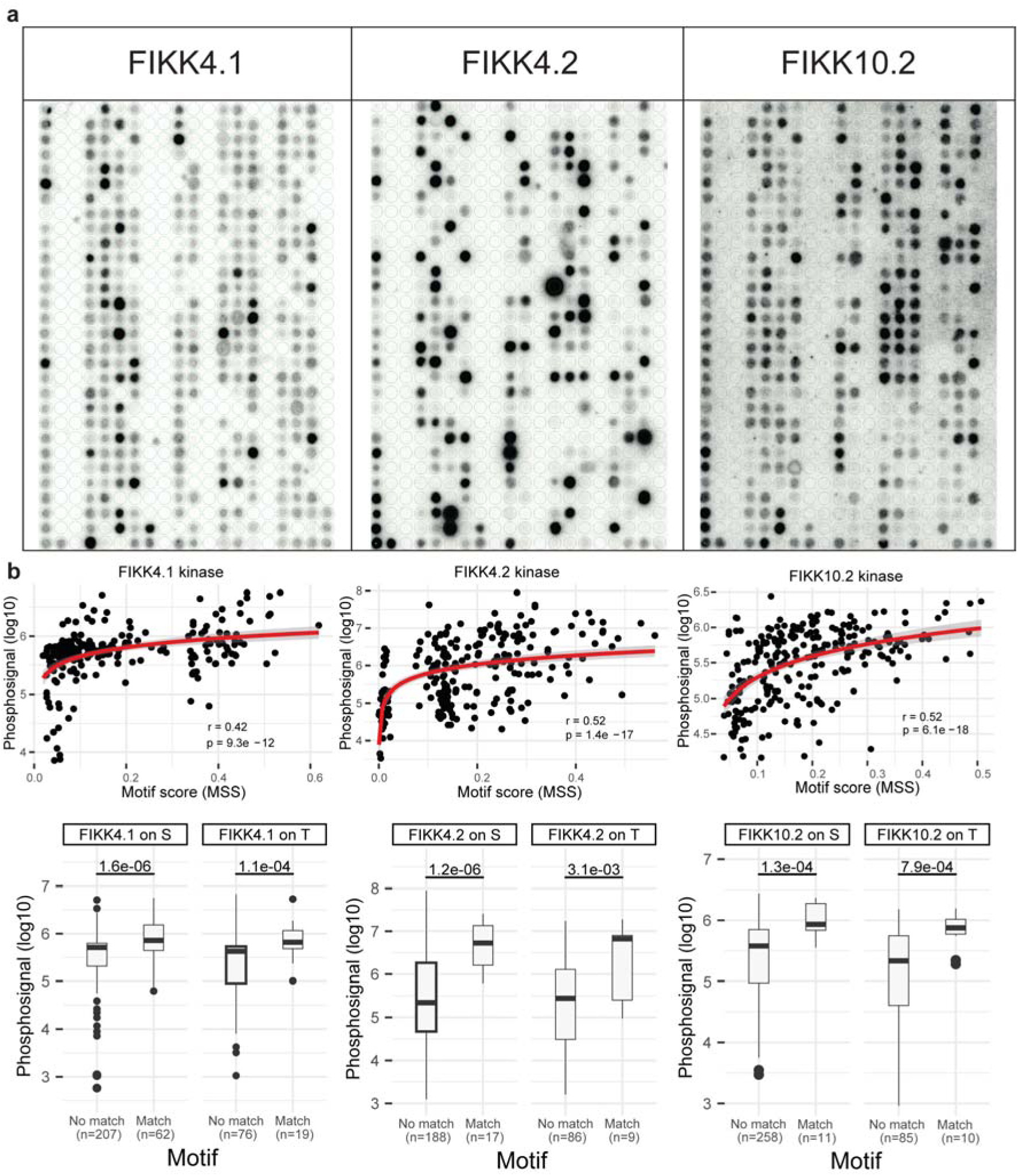
FIKK4.1, FIKK4.2 and FIKK10.2 activity on the phosphoproteome peptides libraries. **a**, Phosphorimager scans of phosphorscreen exposed to phosphoproteome peptide membranes incubated with either recombinant FIKK4.1, FIKK4.2 or FIKK10.2 kinase domains and [γ-32P]-ATP. **b,** Top: correlation of FIKK kinase activity on the phosphoproteome peptide membrane (log10-scaled) against the corresponding FIKK motif score (matrix similarity score) for each peptide. For FIKK 4.1, FIKK 4.2, and FIKK 10.2 kinases. Pearson’s correlation for the y = log(x) curve. Bottom: Difference in FIKK phosphorylation signal (log10-scaled) between peptides without or with a match to the corresponding FIKK motif, for peptides with an S or T phosphoacceptor, for FIKK 4.1, FIKK 4.2, and FIKK 10.2 kinases.

**Extended Data Fig. 13.**
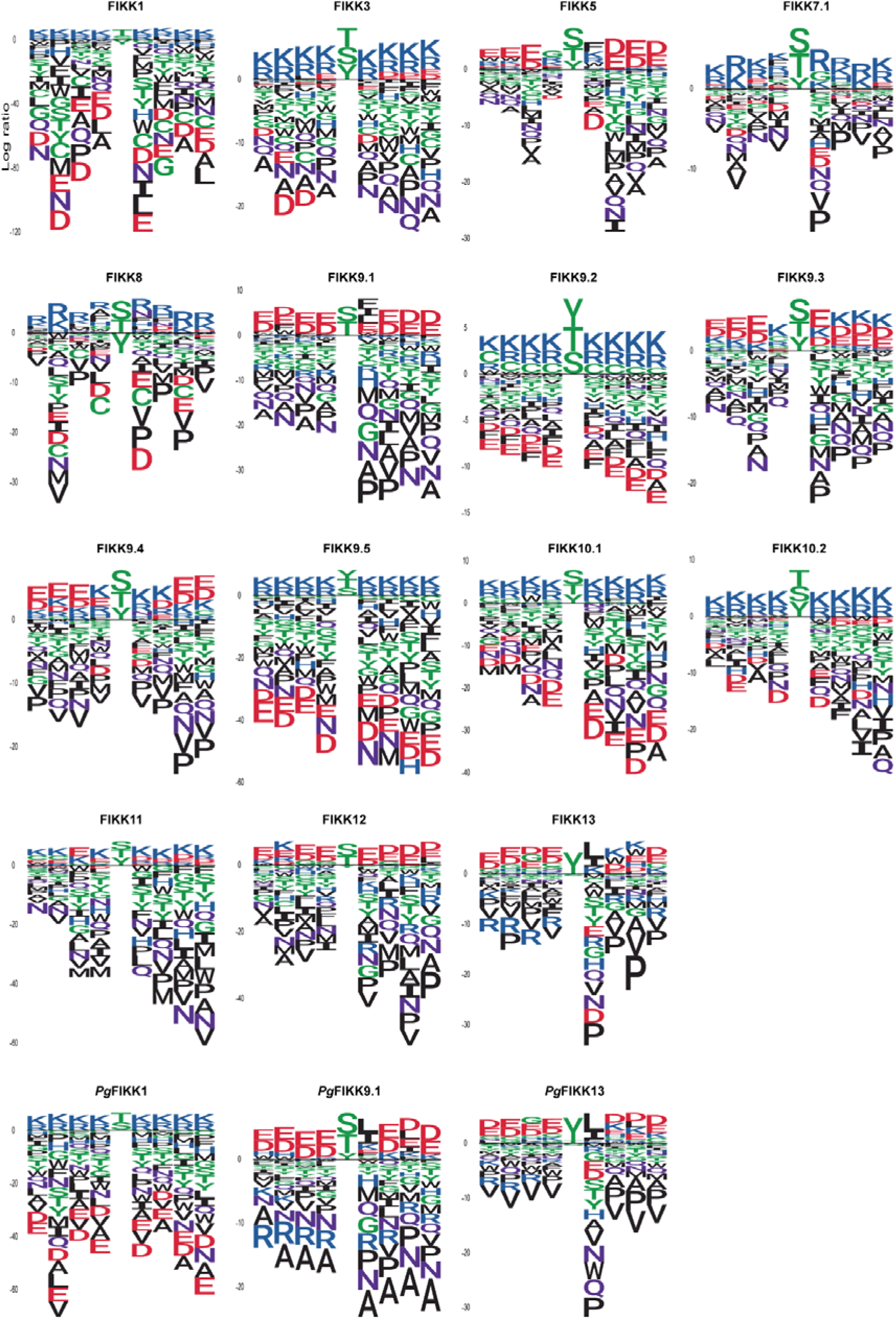
Log2 transformed PWM logos for all recombinant FIKK kinases tested. See Fig. 5f caption.

**Extended Data Fig. 14.**
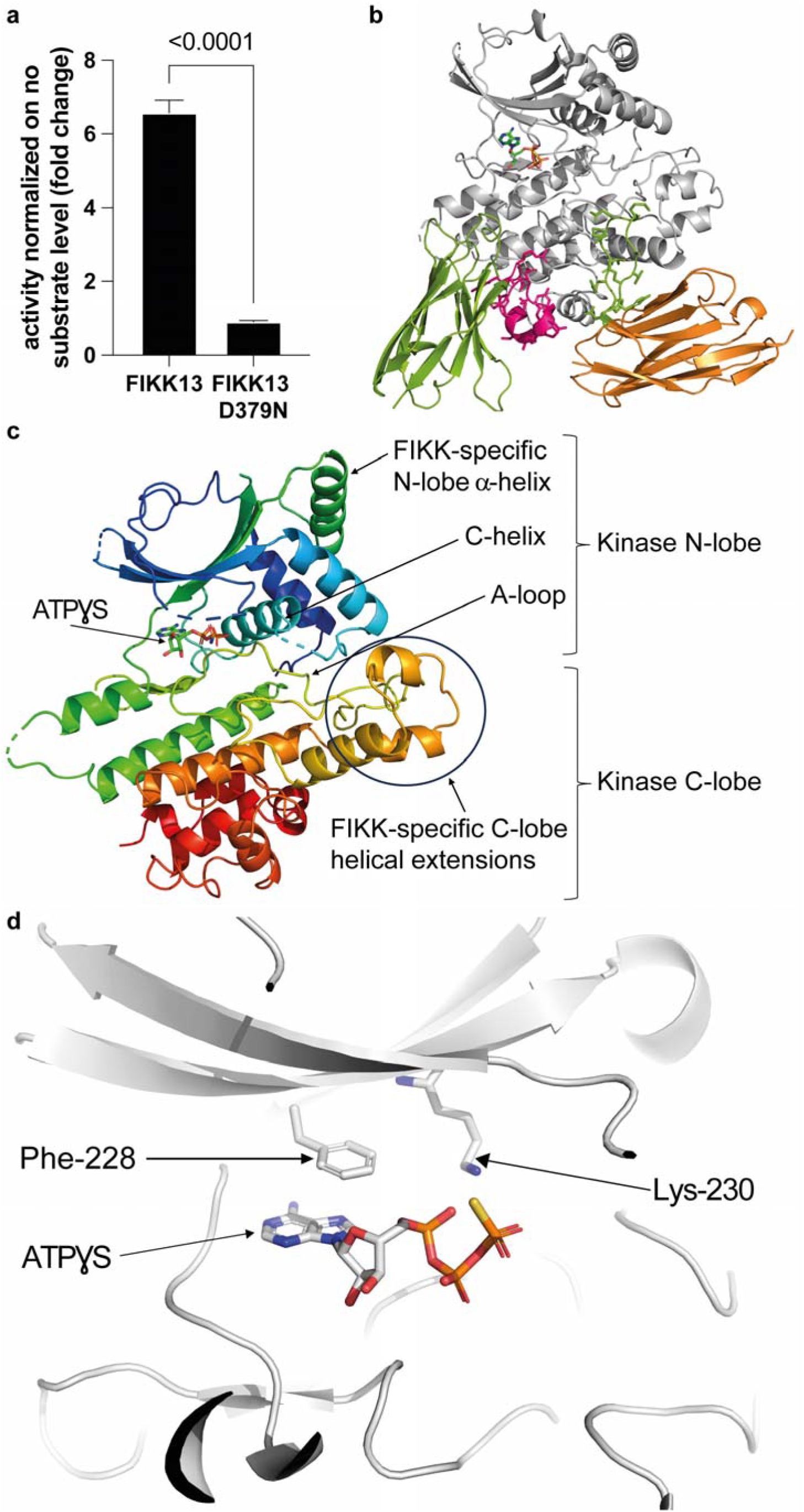
FIKK13 D379N dead mutant crystal structure informs on ATP binding. **a**, Comparison of FIKK13 wild type and FIKK13 D379N phosphorylating activity on cyclic peptide FIKK13_4 using the ADP-Glo assay. Statistical significance was determined using a two-tailed t-test. n=3 biological independent replicates. **b,** The FIKK13 kinase domain – in grey – bound to ATP[S and complexed with Nb9F10 (olive with its CDR3 in magenta) and Nb2G9 (orange with the CDR3 in olive wrapping around the C-lobe of the kinase) **c,** FIKK13 D379N crystal structure with ATP[S. The N-lobe of the FIKK kinases is compact with more features than ePKs including two α-helices packed on top of the conserved C-helix. The A-helix, rarely observed in kinase structures apart from the defining cAMP-dependent kinase PKA^51^, marks the beginning of the N-lobe with the conserved Trp-162 (Extended Data Fig. 18) buried in a pocket between the narrow ends of the aligned A and B-helices positioned on top of the C-helix. The arrangement is capped by an FIKK-specific α-helix inserted between the β4 and β5 strands. The mainly α-helical C-lobe contains, compared to ePKs, three additional α-helices inserted after the activation loop (A-loop). These helices contact the A-loop directly and may restrict its ability to change conformation upon phosphorylation as observed in a number of ePKs^59^. The catalytic machinery of FIKK13 kinase domain is conserved from ePKs with minor changes; the HRD motif where the Asp acts as a general base during phospho-transfer, is conserved as ^377^HLD^379^. The DFG motif, which can switch between the active “DFG-in” and inactive “DFG-out” conformation^60^, is present in FIKK13 as ^398^DLS^400^, although conserved as DFG in FIKK1 and FIKK9.1 (Extended Data Fig. 18) and adopts the “DFG-in” conformation in the FIKK13 kinase domain structure. **d,** Close-up representation of the FIKK13 kinase domain ATP-binding pocket containing the ATP-analogue ATP[S, focusing on the F-I-K-K motif. The size and hydrophobicity of Phe-228 restrict the volume of the ATP-binding pocket while the Lys-230 coordinates the α– and β-phosphates of the nucleotide and forms a salt bridge with Glu-261 of the C-helix which is a hallmark of active ePKs^51^. Taken together, the first experimentally determined structure of a FIKK kinase reveals strong resemblance to ePKs with conservation of the essential elements for catalysis. However, FIKK-specific features, such as the evolution of additional α-helices in both the N– and C-lobe could point to differences in its regulation.

**Extended Data Fig. 15.**
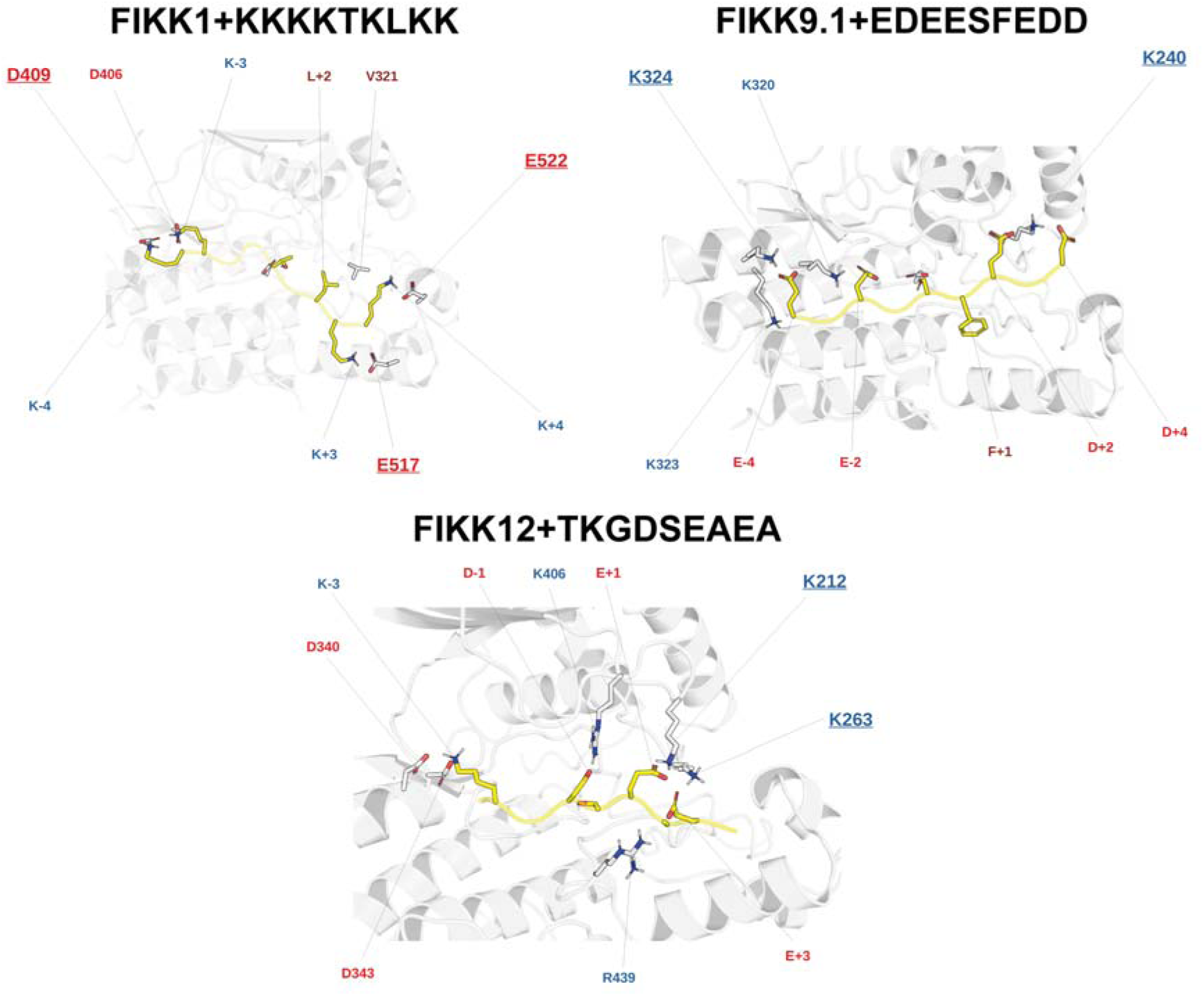
Target peptides of FIKK1, FIKK9.1, or FIKK12 modelled into the substrate-binding groove of the FIKK AF2 structures. (see Methods). Peptides may correspond to a likely target peptide of the kinase, or idealised targets based on the results of the OPAL arrays. FIKK kinase domain coloured in grey and the substrate peptide is coloured in yellow. Negatively charged amino acids are coloured in red, positively charged amino acids are coloured in blue, hydrophobic amino acids are coloured in brown.

**Extended Data Fig. 16.**
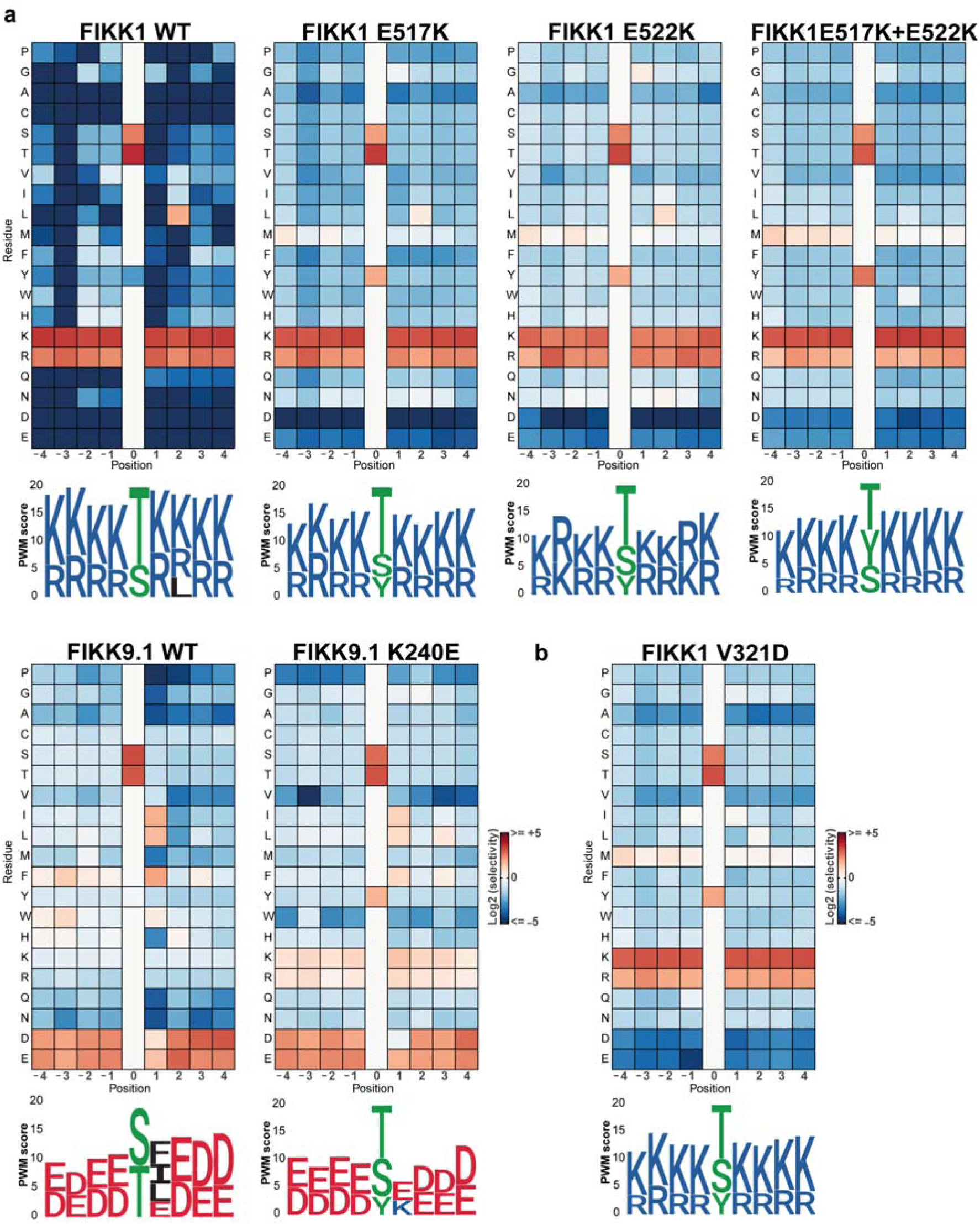
Substrate specificity assessment of FIKK1 and FIKK9.1 kinase mutants using OPAL arrays. **a**, FIKK1 and FIKK9.1 wild type and mutant phosphorylation activity on OPAL membranes represented as heatmaps (see Fig. 3bi caption). Below is represented the PWM logos (see Fig. 3bii caption). **b,** FIKK1 V321D mutant phosphorylation activity on OPAL membranes represented as a heatmap with PWM logo below.

**Extended Data Fig. 17.**
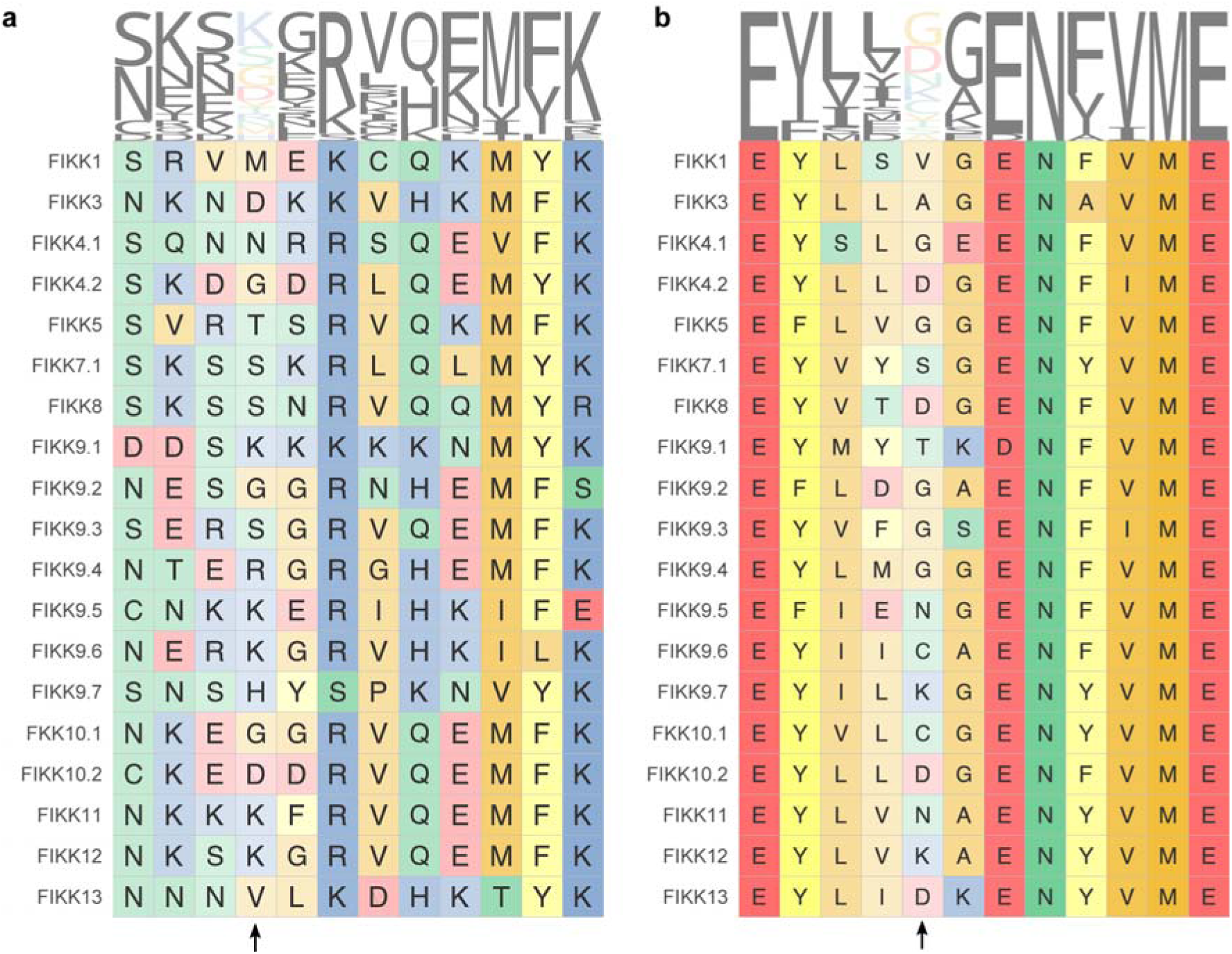
Sequence conservation of FIKK specificity determinants. **a**, Conservation of the region surrounding FIKK12 K212 between FIKK paralogues in *P. falciparum*. The position containing FIKK12 K212 is labelled with an arrow. **b,** Conservation of the region surrounding FIKK12 K263 between FIKK paralogues in *P. falciparum*. The position containing FIKK12 K263 is labelled with an arrow.

**Extended Data Fig. 18.**
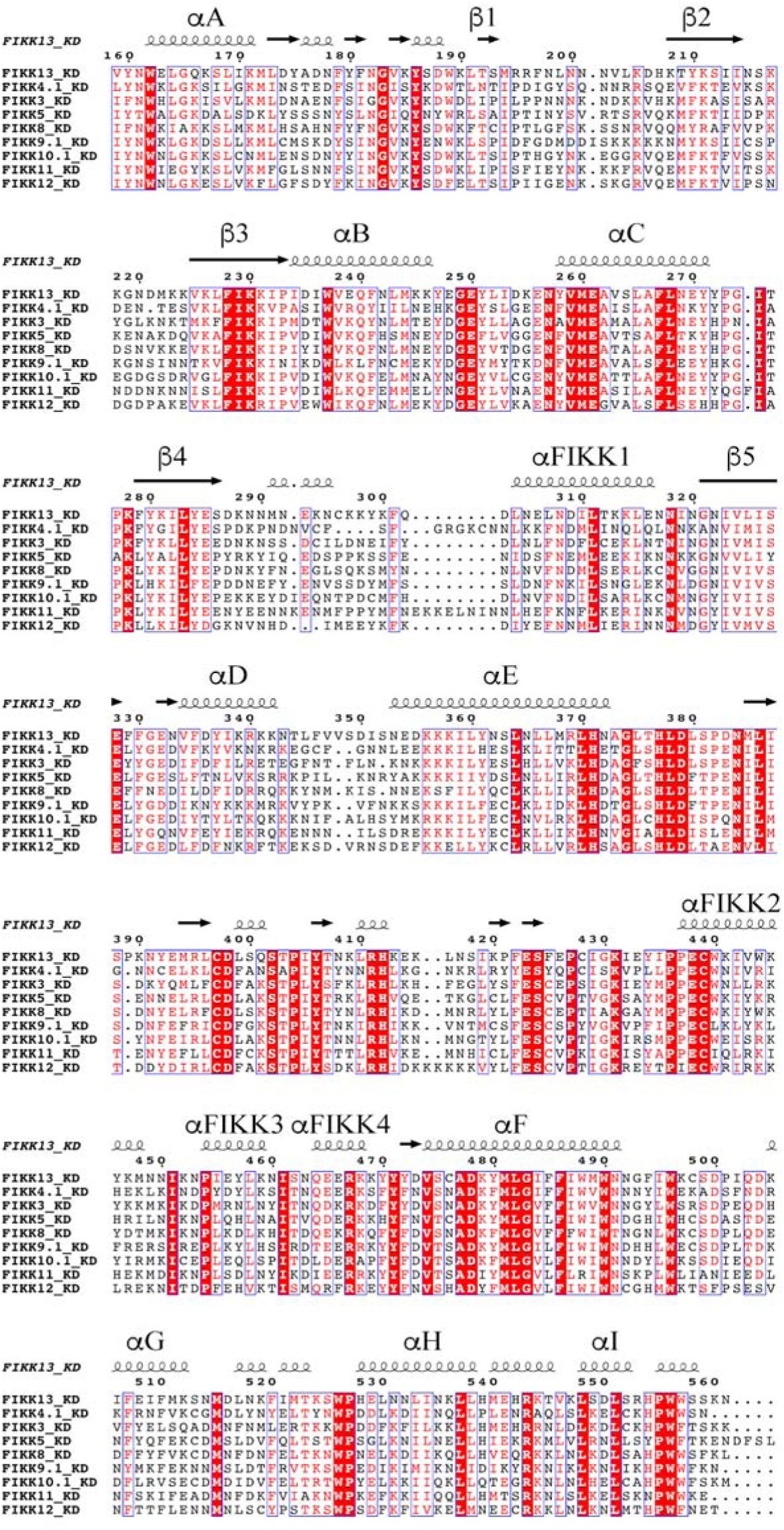
Multiple sequence alignment of various kinase domains. Alignment generated using ESPrit 3.0^68^. The secondary elements in FIKK13 are shown above the alignment. The [-helices and β-strands corresponding to ePKs are labelled. The [FIKK (1-4) are additional alpha-helices found in the FIKK family of kinases, but not in ePKs.

**Extended Data Fig. 19.**
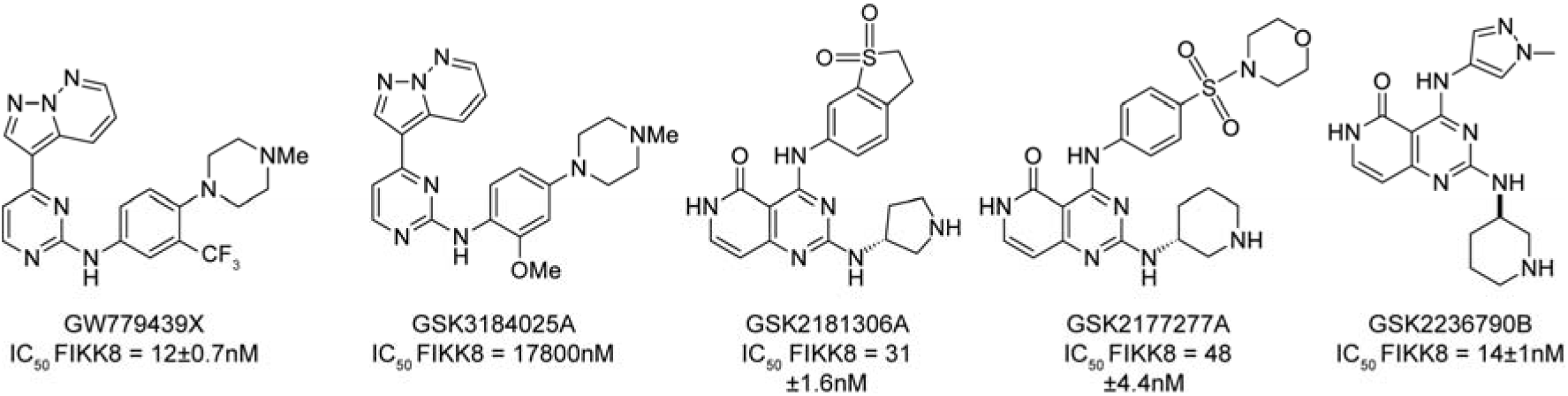
Structure-Activity Relationship assay identifies closely related compounds with different behaviours towards recombinant FIKK8 kinase domain. 333 compounds were identified from the three original PKIS chemical templates. IC50 on recombinant FIKK8 kinase domain was measured in biological triplicate for each one of the compounds and are indicated here for the selected ones ± SD.

**Extended Data Fig. 20.**
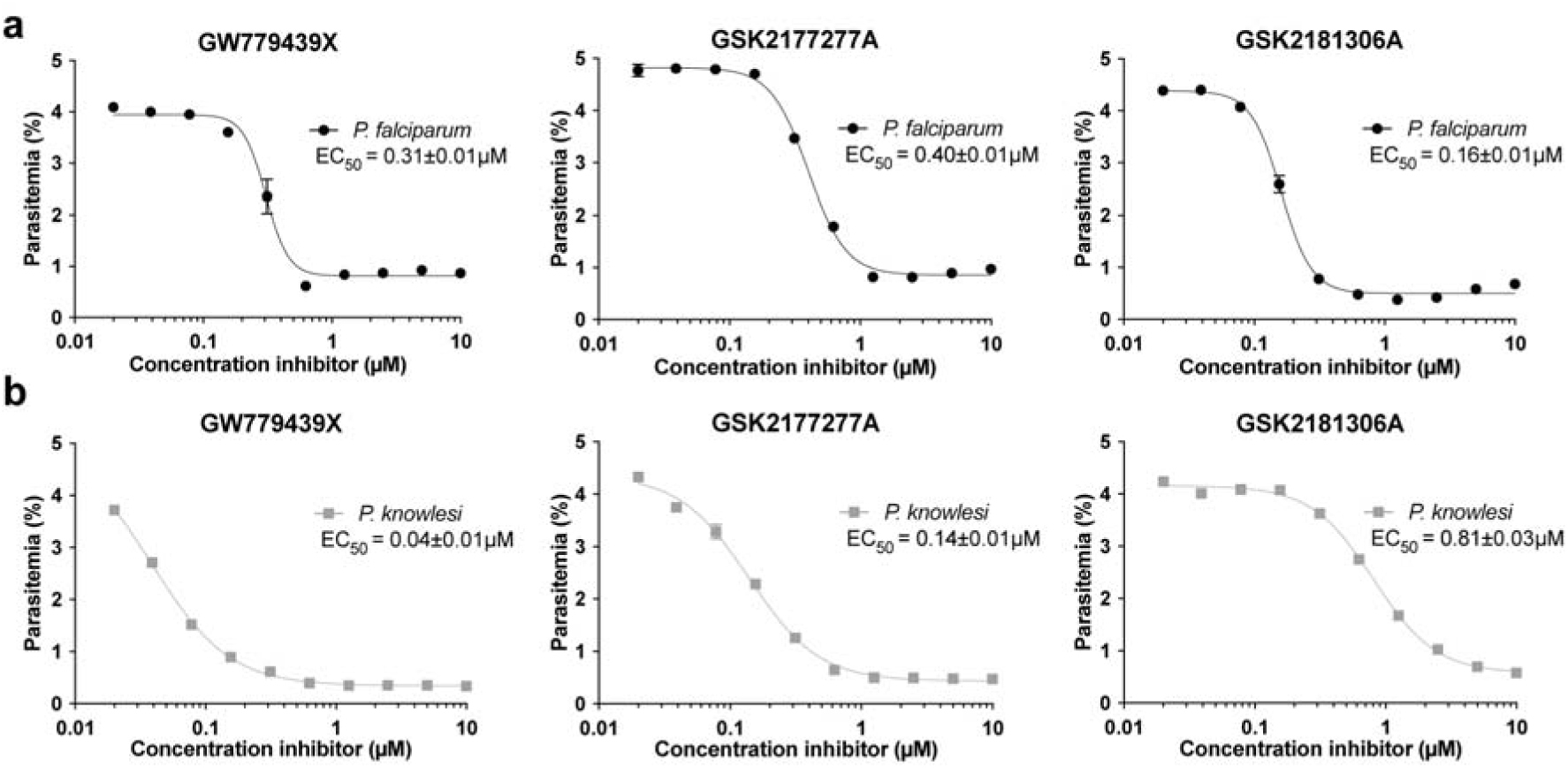
The three most potent *in vitro* FIKK inhibitors kill *Plasmodium* parasites in culture. Half maximal effective concentration (EC_50_) *in vitro* determination for GW779439X, GSK2177277A and GSK2181306A towards *P. falciparum* **a** and *P. knowlesi* **b** parasites. Parasitemia was assessed by flow cytometry using SYBR Green staining of the parasite nucleus after a 72 hours incubation period in the presence of different concentrations of compounds (parasitemia indicated in Supplementary Table 15). EC_50_s were determined using a four-parameter dose-response model with the software PRISM. Data are shown as the mean±SEM for 3 biological replicates. Curves represent dose-response curves of FIKK inhibitors inhibition of *P. falciparum* (black) and *P. knowlesi* (grey).

**Extended Data Fig. 21.**
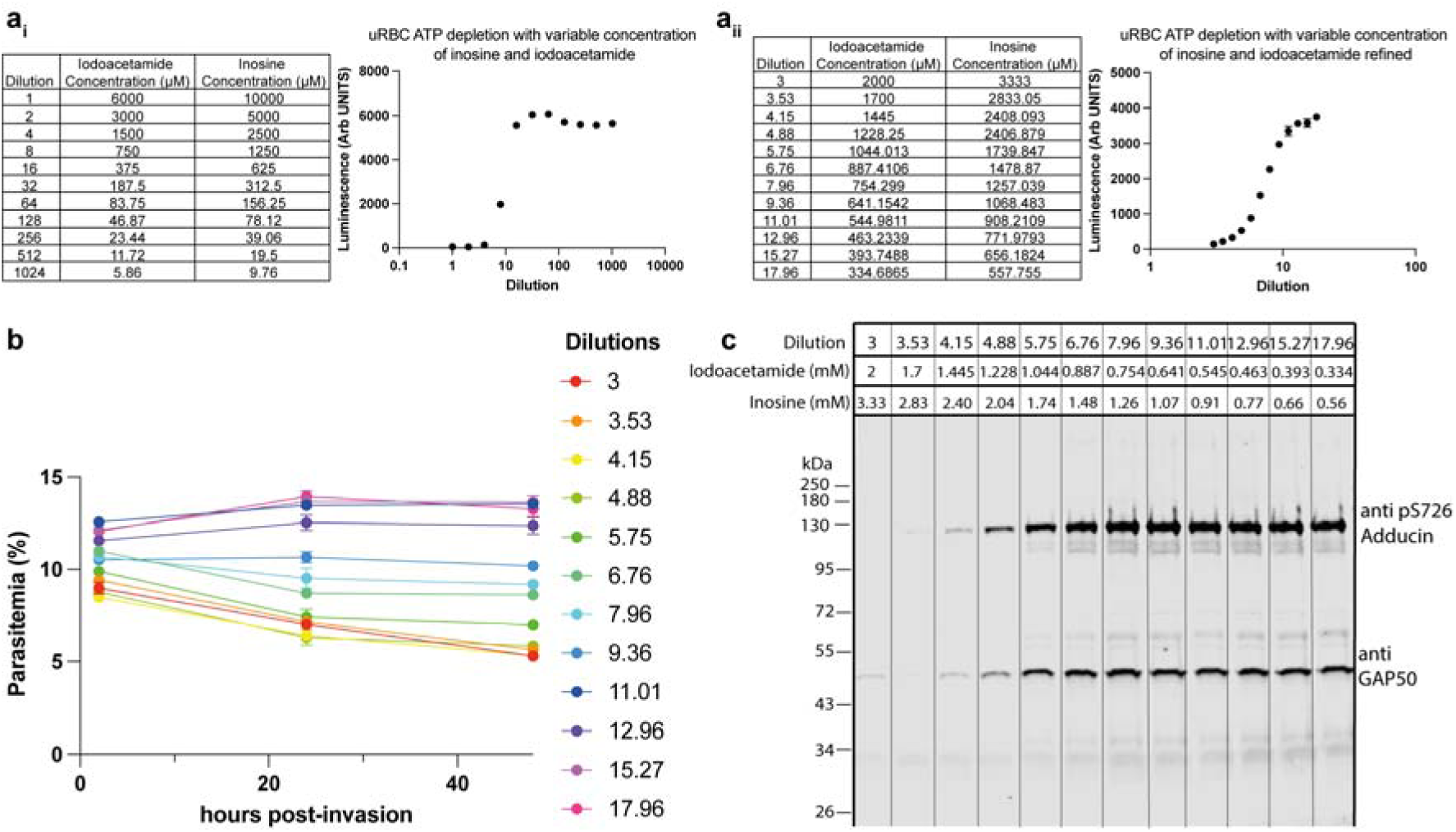
Optimisation of ATP-depletion conditions. **a**, Measurement of intra-erythrocytic ATP concentrations in uRBC using the CellTiter-Glo→ luminescence assay (Promega). (**_i_**) Luminescence, relative to intra-erythrocytic ATP concentration, was measured in uRBC pre-treated with iodoacetamide and inosine concentrations ranging from 6000µM to 5.86µM and 10000µM to 9.76µM respectively. (**_ii_**) Luminescence measured in uRBC pre-treated with iodoacetamide and inosine concentrations ranging from 2000µM to 334.7µM and 3333µM to 557.8µM respectively corresponding to dilution 3 to 17.96 from (**a_i_**). n = 3 biological replicates for both (**_i_**) and (**_ii_**). **b,** Parasitemia assessed for NF54 iRBCs pre-treated with different concentrations of iodoacetamide and inosine corresponding to dilution 3 to 17.96 from (**a_ii_**). Parasitemia was assessed by flow cytometry using SYBR green staining and numerical values of percentages is provided in Supplementary Table 16. n = 3 biological replicates. **c,** Western blot investigating adducin S726 phosphorylation in NF54 iRBCs pre-treated with different concentrations of iodoacetamide and inosine corresponding to dilution 3 to 17.96 from (**a_ii_**). Anti-GAP50 antibody is used here to investigate viability of the parasite.

